# Accurate, fast and memory efficient quantification of immune cell phenotypes in cytometry using machine learning

**DOI:** 10.1101/2024.07.26.605341

**Authors:** Tarik Exner, Nicolaj S. Hackert, Frederic Pohl, Görkem Osmanusta, Fee Schmitt, Hanns-Martin Lorenz, Guido Wabnitz, Georg Schett, Frederik Graw, Jörg Henes, Ricardo Grieshaber-Bouyer

## Abstract

To achieve accurate and reproducible cytometry data analysis, we benchmarked 19 machine learning algorithms for supervised and unsupervised cell classification. The underlying data encompassed 138 million cells from seven independent datasets including conventional flow cytometry, spectral flow cytometry and mass cytometry. We found that tree-based classifiers and in particular Decision Trees, outperformed other approaches in classification accuracy, speed and memory use. High accuracy was achieved even for cell populations rarer than 1% using decision trees. We validated our decision tree-based approach in a clinical setting using diagnostic blood T cell phenotyping of 107 patients. Automatic quantification of CD4 helper T cell phenotypes achieved 99 % accuracy compared to manual expert assessment. Finally, we combined automated data transformation, supervised and unsupervised gating, an application program interface and a user-friendly desktop-application into FACSPy and FACSPyUI, a fast and scalable open-source toolbox for the analysis and visualization of cytometry data.

## Introduction

Cytometry is an essential method for cell phenotyping and quantification and has been significantly advanced its methodology during the last years. Mass cytometry (CyTOF®) and spectral flow cytometry were developed to overcome spectral limitations of conventional flow cytometry, enabling simultaneous detection and analysis of a growing number of markers simultaneously on each single cell. The resulting data is high-dimensional; therefore, data analysis becomes increasingly challenging. The high dimensionality necessitates the use of sophisticated algorithms and techniques to navigate the data volume and recognize relevant structures.

Current analysis approaches using commercial software excel in user-friendliness and routine cytometry analysis tasks, such as manual gating, extraction of gate frequencies and absolute fluorescence values (e.g. FlowJo^1^, FCSExpress, CytoBank and others). Recently, the option of clustering and support for dimensionality reductions were added. However, these applications are typically closed source, associated with considerable costs and neither easily customizable nor extendable, rendering them inflexible when specific analysis tasks are demanded. On the other hand, specialized open-source software packages in Python and R offer advanced analysis capabilities but require a deep understanding of the programming language, posing a barrier for many users^2–5^.

While there are varieties of software packages with a potentially unlimited amount of analysis possibilities available, we identify four main challenges in the composition and organization of those tools, which are not solely specific to cytometry analysis: First, the landscape of analysis packages resembles a patchwork, where dedicated functionality is often not embedded into larger frameworks but rather function as isolated tools. While there can be advantages in specialized data structures found in individual packages, this makes portability and combinability difficult. Second, the development of computational solutions to analysis tasks, while it may be seen as automation, often requires secondary steps to embed them into analysis pipelines to unfold their full potential. Third, there are currently two main open-source programming languages for data analysis, namely Python and R, that often do not share the latest developments due to barriers for portability of code and objects between the two. While there have been attempts for increased interoperability (e.g. reticulate^6^ or rpy2 (https://rpy2.github.io)), there are several issues associated, such as computational overhead, difficulties in setup and loss of functionality of the respective objects during the conversion process^7^. Finally, all these software packages require expertise in programming in general and in specific, often hindering data analysis due to the individual lack of experience in that field. Here, we overcome these problems by developing a versatile, ready-to-use application programming interface (API), called FACSPy. We embed and automate common analysis tasks for cytometry analysis into a specialized, yet highly flexible framework. In FACSPy, we make extensive use of the *anndata* object as the underlying data structure due to its portability to other programming languages and compatibility with various pre-existing algorithms, including the scverse ecosystem^8^. During development of FACSPy, we focused extensively on providing a usable interface specifically equipped for the needs of cytometry analysis, while keeping the system flexible to be portable to other programming languages, integratable into other python frameworks and accommodate future algorithms. In addition, FACSPy is fully scriptable, allowing for easy automation of data analysis. We leverage supervised and unsupervised machine learning algorithms for the automation of cytometry gating and provide an extensive benchmark of suitable classifiers. Finally, we developed FACSPyUI, a desktop-app interface that includes the functionality of FACSPy but does not require pre-existing knowledge of programming.

In order to validate our system, we benchmark different approaches for automated cell quantification and validate the preferred approach, Decision Trees, in a clinical setting of human blood T cell phenotyping in 107 individuals.

## Results

### Benchmarking of supervised gating algorithms across different cytometry data entities

To determine suitable algorithms for supervised cell type classification, we tested 15 algorithms across seven different cytometry datasets (**Figure 1A**, see also **Methods**). Manual expert gating was defined as the ground truth and the classification accuracy was calculated based on concordance. In total, we analyzed 138 million cells from flow cytometry, spectral flow cytometry and mass cytometry containing both human and mouse immune cells (**Table 1**).

**Figure 1:**
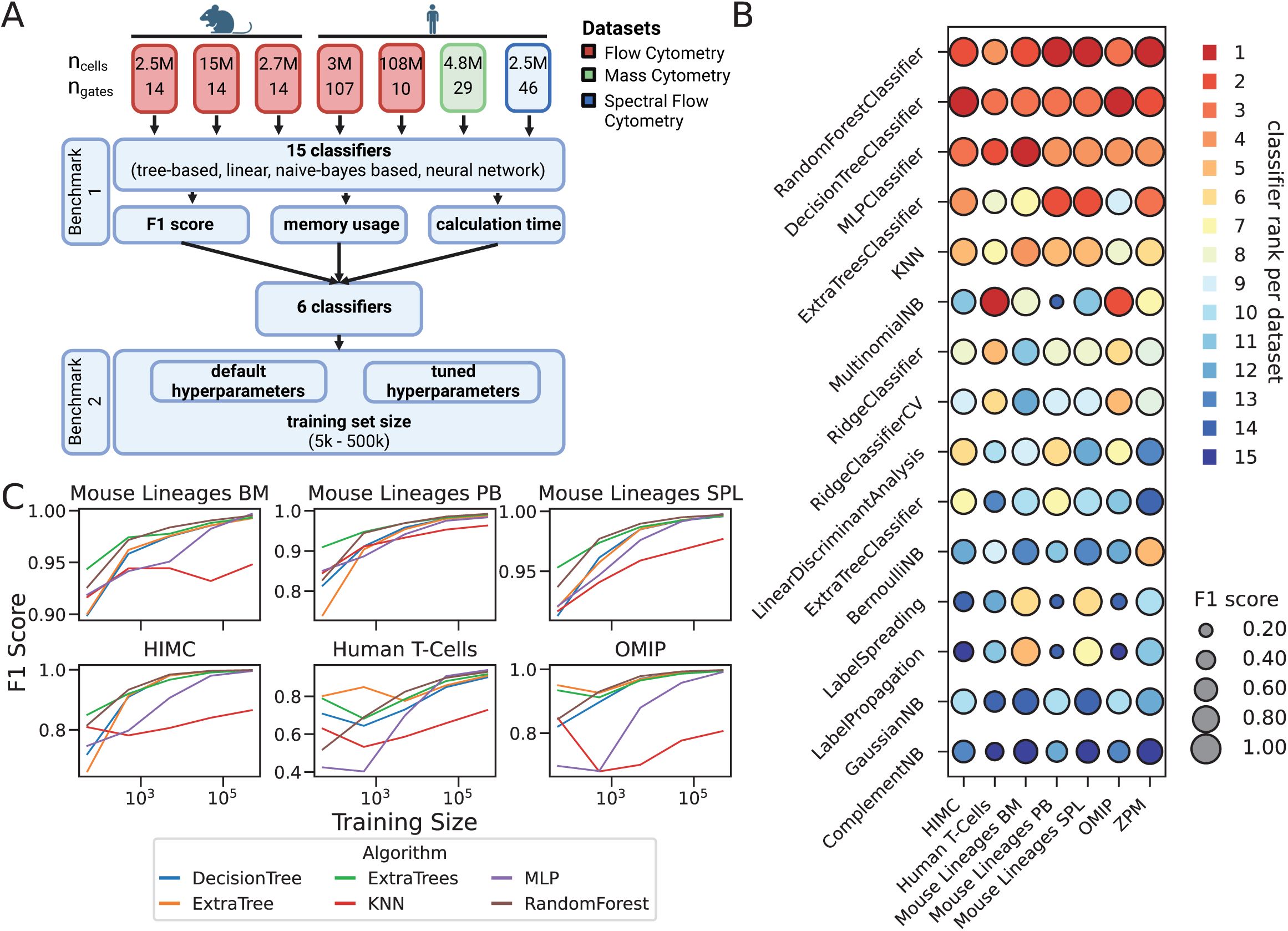
Benchmarking machine learning algorithms for cytometry data analysis. **A** Graphical abstract. Seven different cytometry datasets were used to compare the supervised classification algorithms using a subset of 10.000 cells of n-1 samples. The most promising classifiers based on F1-score (see **B**), memory usage and calculation time (compare **Extended Data Figure 2A and 2B**) were subjected to a second benchmark where the effect of hyperparameter tuning (compare **Extended Data Figure 2C**) and the training set size was evaluated. **B** Classifier comparison. The indicated classifiers were subjected to the benchmark on the indicated datasets as described above and scored using the F1-metric. The diameter of the circles represents the median F1-score per dataset, while the color code represents the relative rank of the classifier’s F1-score compared to all other algorithms for a given dataset. Classifiers are ordered in y-direction by the median F1-score across all datasets. **C** Effect of train set size on the classifier F1-score. Classifiers were instantiated with tuned hyperparameters and trained on the indicated event numbers ranging from 5×10¹ to 5×10⁵ events per training set. Maximum performance is reached at 500.000 events per training set, although convin-cing results are reached with much lower training set sizes for most of the algorithms.

**Table 1.** Datasets used in the study.

| **Dataset** | **Technology** | **Description** | **Panel size** | **Total samples** | **Total cells** |
| --- | --- | --- | --- | --- | --- |
| 1 | Flow cytometry | Bone marrow cells of C57BL/6 mice | 12 | 18 | 2,450,306 |
| 2 | Flow cytometry | Blood cells of C57BL/6 mice | 12 | 18 | 14,814,725 |
| 3 | Flow cytometry | Spleen cells of C57BL/6 mice | 12 | 18 | 2,753,919 |
| 4 | Flow cytometry | Human blood leukocytes | 17 | 14 | 3,016,910 |
| 5 | Flow cytometry | Human blood leukocytes | 7 | 107 | 108,045,599 |
| 6 | Spectral flow cytometry | Human blood leukocytes | 41 | 4 | 2,572,304 |
| 7 | Mass cytometry | Human blood leukocytes | 42 | 19 | 4,750,000 |
| **Total** |  |  |  | **198** | **138,403,763** |

Across all seven datasets, tree-based methods showed the highest classification accuracy, followed by the KNN-Classifier and the Multi-layer-perceptron (**Figure 1B**). The RandomForestClassifier showed the highest median F1 score (0.98), followed by the DecisionTreeClassifier (0.97) and ExtraTreesClassifier (0.94), when trained on 9000 events. In contrast, linear methods and naïve bayes based methods performed substantially worse, with median F1 scores ranging between 0.27-0.92 (**Figure 1B**).

Next, we benchmarked the performance of these algorithms according to calculation time and memory consumption. Of the algorithms with high accuracy, the KNN classifier exhibited prohibitively large compute time needed for predicting the validation samples **(Extended Data Figure 2A**). While highly accurate and fast, the RandomForestClassifier and ExtraTreesClassifier displayed varying memory use dependent on the panel and sample size **(Extended Data Figure 2B**). Thus, the DecisionTree Classifier, ExtraTreeClassifier and MLPClassifier outperformed other algorithms with respect to accuracy and computing performance.

### Effect of hyperparameter tuning and sample size on classification accuracy

We next selected the tree-based classifiers, the multi-layer-perceptron and the KNN-based classifier for further evaluation and tested the impact of hyperparameter tuning (also see **Methods**) on classification accuracy and performance. Hyperparameter tuning significantly increased the median F1 score (0.88 to 0.94) for the algorithms (**Extended Data Figure 2C, Supplementary Figure 1A–7A**), in particular for ExtraTree-(0.66 to 0.96) and ExtraTrees-Classifier (0.93 to 0.96).

We then sought to explore the necessary sample size needed for the classifier training. Classification accuracy of tuned classifiers was tested as a function of the number of training cells. Accurate cell type classification was already feasible with a training set consisting of 500 cells, resulting in an F1-score of 0.78 – 0.94 and reached a plateau at 5000 events for most cell types to provide an accurate result with a median F1 score of 0.97 across tree-based classifiers (**Figure 1C**). A further increase of the number of training examples led to a higher overall accuracy in the validation datasets (**Figure 1C, Supplementary Figure 1A–7A**).

Together, these results demonstrate that DecisionTreeClassifiers are fast and accurate for supervised cell classification in cytometry data.

### Supervised gating accurately identifies CD4 helper T cell quantities in a clinical setting

To validate the practical applicability, we tested if supervised gating could accurately quantify T cell phenotypes in peripheral blood mononuclear cells. The dataset consisted of 107 flow cytometry samples of human peripheral blood cells used for clinical immune phenotyping. Based on our benchmarking results, we trained a hyperparameter tuned Decision Tree Classifier on the cells of 8 samples from the dataset. We found high concordance of the quantified cell populations in the ungated samples as evidenced by the F1-scores (**Figure 2B**) and the frequency of parents of a manually drawn expert gate (left) and the AI-gated cells (right) (**Figure 2C and 2D**). We next subjected the other six datasets to the same analysis and also found highly comparable results between the manually defined expert gates and the gates reproduced by machine learning **(Supplementary Figure S8-S13)**. Thus, accurate, fast and memory efficient quantification of immune cell phenotypes in cytometry can be achieved using machine learning.

**Figure 2:**
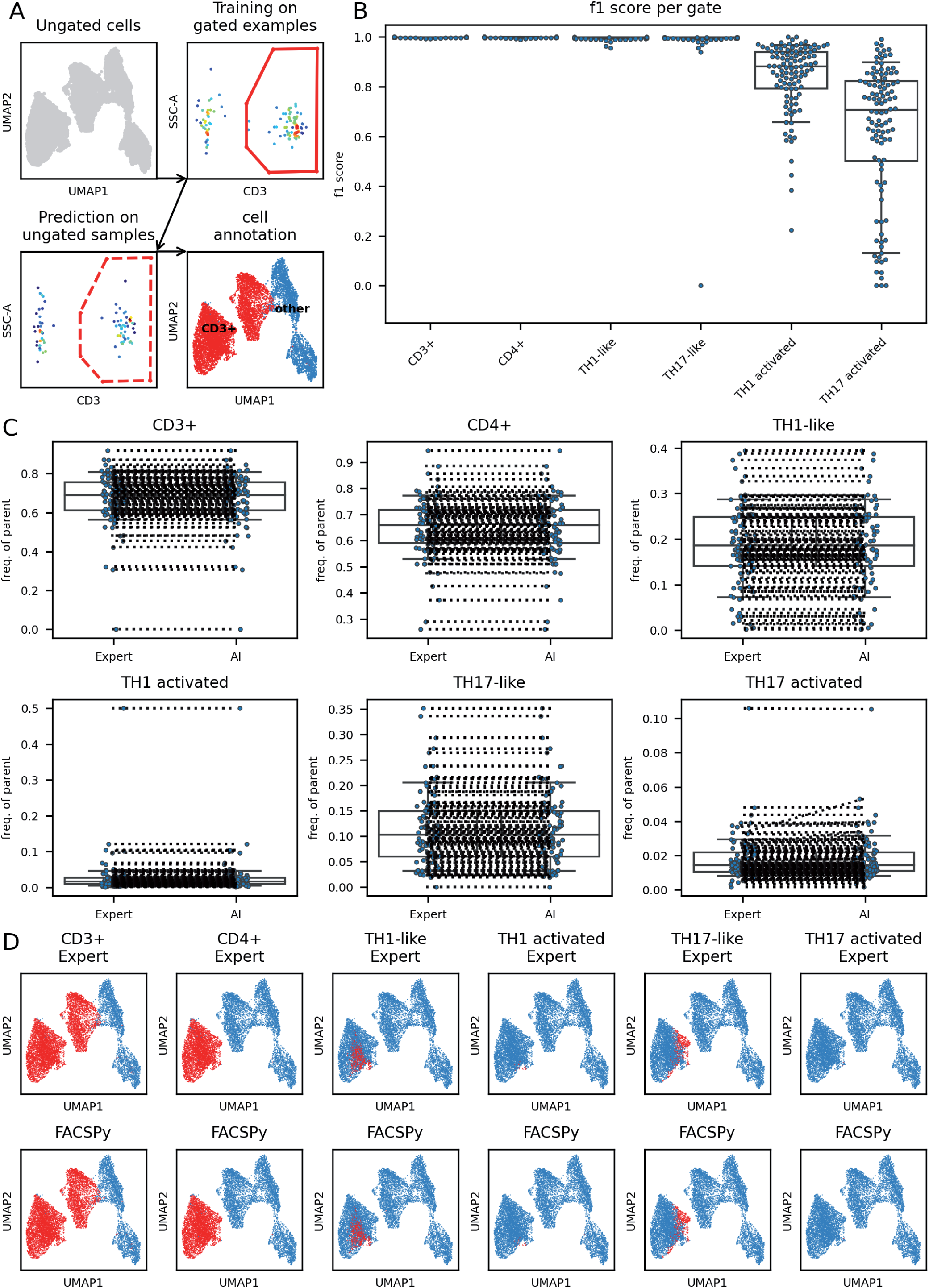
Supervised gating for cell identification. The dataset consisted of 107 samples of human peripheral blood cells (flow cytometry; Dataset 5). **A** Graphical abstract. Example samples are used to train a classifier using the detected marker expression and the gates. The classifier is then used to classify previously ungated cells. **B** A Decision Tree classifier with tuned hyperparameters (compare **Supplementary Figure S6**) was trained on eight samples of the dataset. The remaining samples were predicted. F1 scores are plotted on the y-axis respective to the individual gates (x-axis). Each dot corresponds to one sample that was not used for training. **C** High concordance of the frequency of parents of a manually drawn expert gate (left) and the AI-gated cells (right). Dotted lines connect corresponding samples. D High concordance of cell classification on a single cell level. Shown is a UMAP embedding colored for the indicated cell type in red. Top row: cells classified by manual expert gating. Bottom row: cells classified by supervised classifica-tion.

### Semi-supervised gating identifies populations of interest without selection bias

Supervised, bi-axial gating strategies rely on manually, and therefore subjectively, drawn gates and a-priori information. To overcome this limitation, we tested a semi-supervised gating strategy, which utilizes user-defined marker expressions for each cell type. Using unsupervised clustering, cells are divided into communities, which are subsequently assessed for the median cluster-specific marker expression. By defining markers with positive expression in the respective cell types, communities fulfilling this criterion are selected and classified to be member of a specific cell type (**Figure 3A**).

**Figure 3:**
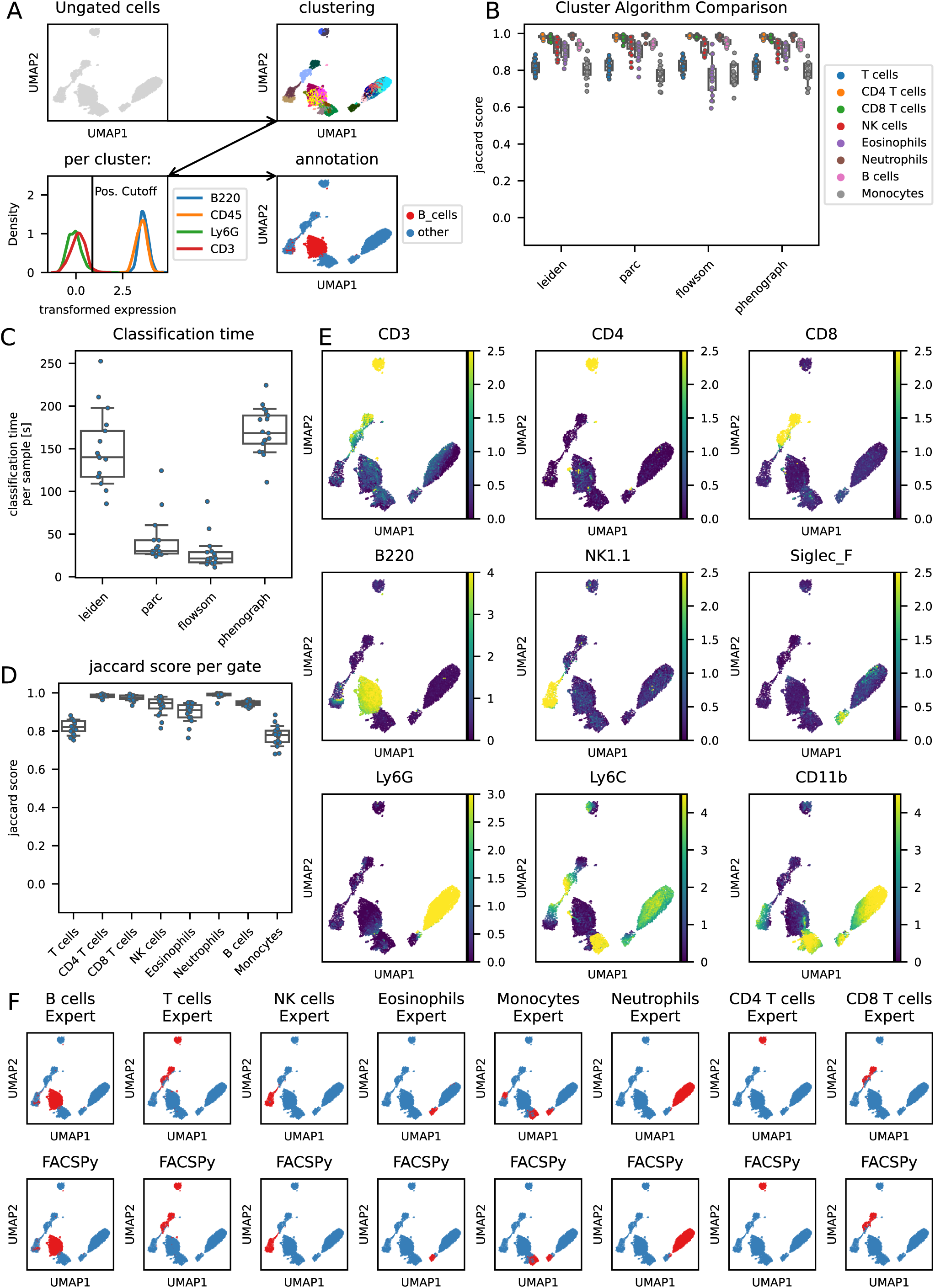
Semi-supervised gating. **A** Graphical abstract: ungated cells are clustered and each cluster is inspected for its median marker expression. Marker intensity is divided into positive, low, intermediate or high expression and cells are classified based on the user defined cell characterization. **B** Algorithm comparison. The indicated clustering algorithms were used and the marker expressions per cluster were used to gate cells as described in A. **C** Algorithm benchmark. Clustering time was compared for the indicated algorithms. The PARC algorithm showed comparable calculation times compared to FlowSOM, while maintaining high accuracy. **D** PARC algorithm was used with tuned hyperparameters (compare **Supplementary Figure S22**), and accuracy scores are plotted per gate. Each dot represents a single sample analyzed. **E** Marker expression on UMAP embedding. **F** High concordance of cell classification on a single cell level. UMAP embeddings are colored for the indicated cell type (red) for either manual expert gating (top row) or semi-supervised determined cell types (bottom row).

As a control experiment, we calculated jaccard scores (defined as the size of the intersection divided by the size of the union of two label sets) of populations that have been gated with alternative gating strategies but were aimed at quantifying the same cell type. First, we compared a gate which is drawn to identify marker-positive cells to gating strategies, which first use exclusion of other cell types. We identified contaminating cell types when using only the cell-type specific marker, which could be excluded using the exclusion gating strategies. Using a cluster-based-approach, cell communities with only a fraction of marker-expressing cells were identified to be cell-type negative by the unsupervised approach and therefore matched the exclusion gating strategies better, without supplying marker negativity in the semi-supervised gating strategy **(Extended Data Figure 3B**).

Cell classification via unsupervised machine learning poses challenges when dealing with markers with continuous, as opposed to bimodal, expression. Therefore, as a second approach, we compared different quadrant-gatings of populations, whose marker expressions were continuous. A continuous marker expression suggests that no clear separating line in two dimensions can be defined that would accurately classify the events. As expected, the different gating strategies resulted in vastly different cell assignments as quantified by the jaccard score and visualized in UMAP space **(Supplementary Figure S14)**.

These experiments suggested that manual gating in cytometry data does not define an unambiguous ground truth. Therefore, we chose jaccard indices instead of F1-scores as a metric to quantify the quality of cell identification.

Using our datasets with manual gating as a control, we compared different algorithms for community detection. We find that graph-based algorithms (phenograph, leiden, PARC) were on par with the widely used FlowSOM clustering in detecting biologically relevant cell type clusters reflecting cell identity (**Figure 3B, Supplementary Figures 15B–20B**). Usage of the PARC algorithm resulted in a more rapid clustering compared to the leiden- and phenograph implementations (**Figure 3C, Supplementary Figures 15C–20C**). Hyperparameter tuning did not have striking effects on the overall jaccard index of this approach (**Supplementary Figure 21-27**), although a high clustering resolution could improve the performance in some datasets **(Supplementary Figure 24-27**).

We applied this approach to all seven datasets and found a very high concordance of cells identified by manual gating as judged by the jaccard score and visual inspection of the datasets on a single cell level based on their marker expressions (**Figure 3D–F, Supplementary Figures 15D–F to 20D–F**).

### FACSPy is an end-to-end analysis platform

To enable end-to-end cytometry data analysis with full machine learning and automation implementation, we developed the *FACSPy* platform. The cytometry data are automatically annotated using provided metadata and panel information, optionally gated using a provided FlowJo® workspace, and saved as an .h5ad file. This file can be read using different *anndata* implementations in R, python or Julia, allowing for the use of analysis steps which are not currently implemented in python (**Figure 4A**). FACSPy is deliberately designed to store analyses for multiple populations within the same *anndata* object, allowing for comparative and parallel analysis of different populations without the need for data subsets. (**Figure 4B**).

**Figure 4:**
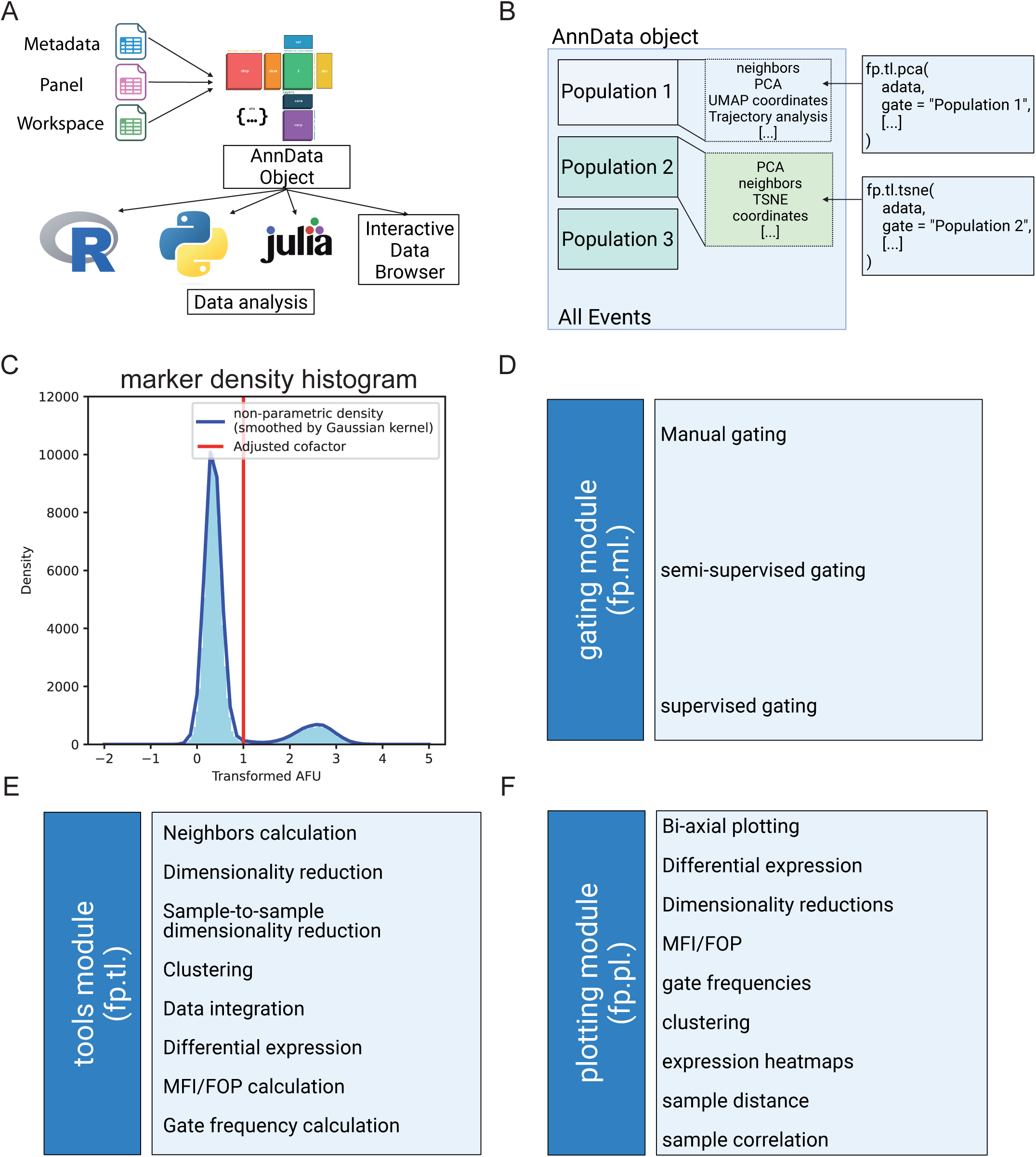
Graphical abstract of the FACSPy API. **A** Automated data annotation. Metadata, the panel information and a workspace are concatenated into an AnnData object containing the compensated cytometry data. Leveraging the inter-operability of AnnData objects between different programming languages, the analysis can be performed in python, R, Julia or the FACSPyUI (Interactive Data Browser). **B** As cytometry data store multiple populations of interest, FACSPy stores population specific analysis within the same anndata object. **C** Automated data transformation. Histogram analysis per channel is performed to determine the cutoff of a bimodal distribution which is in turn used for arcsinh transformation. **D** The tools module. This module offers analysis functionality such as the calculation of neighbors, dimensionality reduction or differential expression. Data integration and expression recalculati-on can be performed. **E** The gating module. Gating can be accomplished using semi-supervised approaches and a user-defined marker expression table or supervised using a gating strategy provided for example samples. Manual gating strategies are automati-cally imported or realized via FACSPyUI. **F** The plotting module. A dedicated module is used for data visualization, including classic bi-axial plotting and frequency plots, low-dimensional median fluorescence intensity plots or high-dimensional plots including dimen-sionality reductions and expression heatmaps.

The API further includes the newly developed approaches for automated cofactor detection (**Figure 4C, Extended Data Figure 1 and Methods**), while populations of interest are defined using manual predefined gating strategies or leveraging the API-provided supervised or semi-supervised approaches (**Figure 4D**).

The *tools* module (**Figure 4E and Extended Data Table 2**) offers a broad functionality with common tasks of single cell analysis, such as differential expression analysis, dimensionality reduction and data integration. Furthermore, as cytometry analysis is frequently carried out on a large scale covering potentially hundreds of samples over a wide range of experimental conditions, we implemented dedicated functions for the analysis of sample-to-sample relationships, such as samplewise dimensionality reductions, sample correlations and expression analyses per sample.

In order to provide easily accessible data visualization, we implemented a dedicated *plotting* module (**Figure 4F and Extended Data Table 1**) that covers the most frequently used plot-types in cytometry analysis as high-level functions. While this module produces publication-ready plots out of the box, the user is still enabled to access the respective raw data leading to the plot directly from the function, if he wishes to use other third-party plotting utilities. Further, the plotting module produces plots that are seamlessly integratable into the widely used *matplotlib/seaborn* ecosystem, so that raw plots produced by FACSPy can easily be modified using the normal *matplotlib* syntax. This implementation further highlights the above-mentioned flexibility while still providing a reliable toolbox for the tasks at hand.

In order to build on pre-existing syntax and infrastructure, FACSPy leverages the *scanpy*^9^ notation for functions which ensures a quick understanding of the API for users already familiar with single cell analysis in python using *scanpy*.

### FACSPyUI is a desktop app enabling interactive data analysis

To enable users without programming experience to leverage the FACSPy API, we developed a desktop app interface called FACSPyUI for interactive data analysis.

FACSPyUI is implemented as a PyQt5 app that can be run locally. The backend uses FACSPy exclusively (**Figure 5A**) and implements the same steps to create a dataset as described above. To create a dataset, accompanying metadata and the panel information can either be loaded from a tabular file or entered directly into the app. The transformation method can be selected and adjusted using an intuitive graphical user interface (GUI). Finally, the dataset is saved to a hard drive or kept in memory for concurrent analysis.

**Figure 5:**
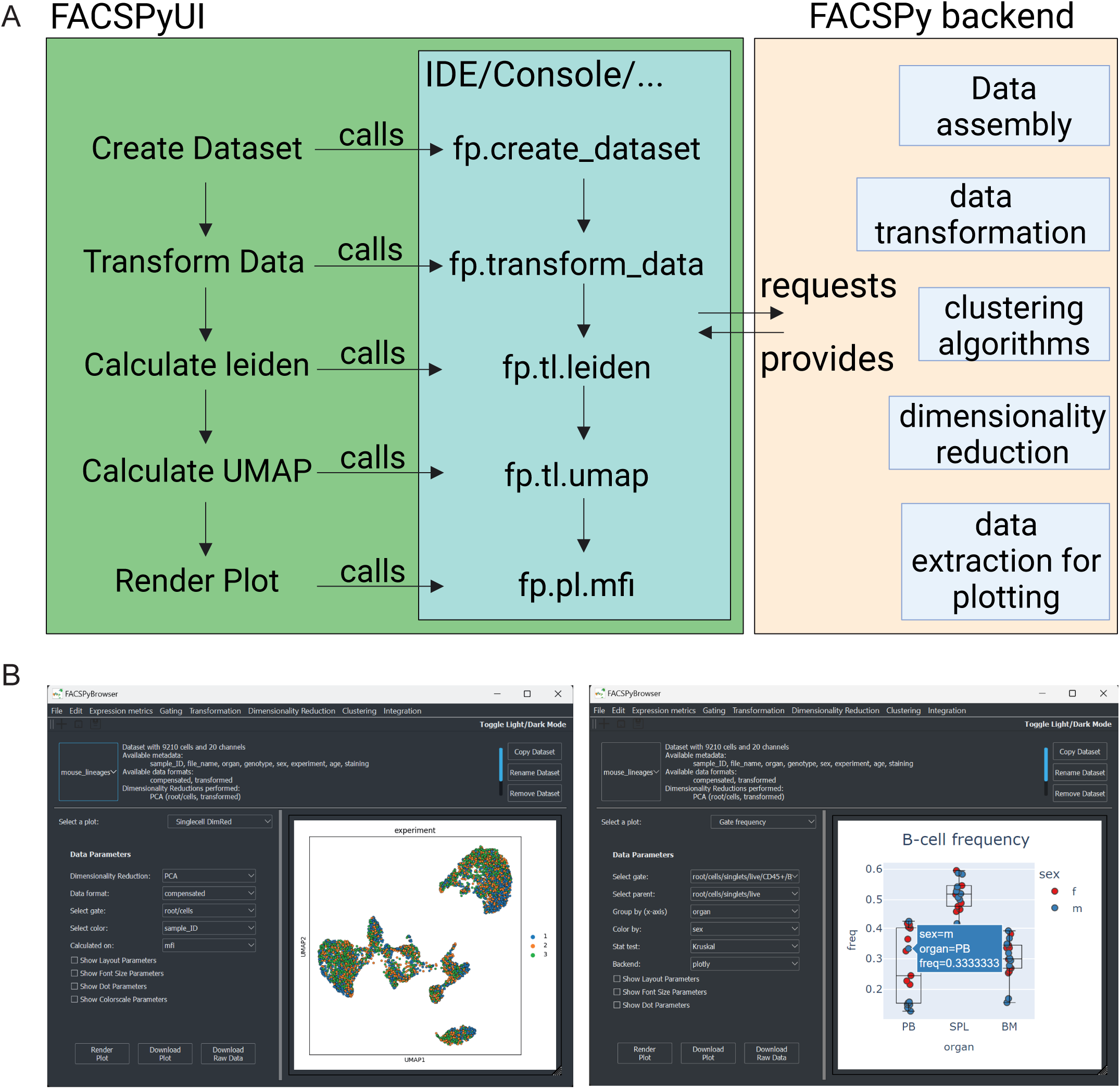
A desktop app interface for programming free data analysis – FACSPyUI. **A** Graphical abstract. Internally, the FACSPyUI (green) uses the same functions that are used by the user when using the API via a console or an integrated development environment (IDE, light blue). The functions then use the FACSPy-backend (orange) to run calculations, assemble data or extract data for plotting. **B** Data analysis. The app is divided into multiple tabs covering implemented analysis steps. Publication-ready plots can be assembled and exported using matplotlib (left side) or can be rendered via the interactive plotly backend (right side).

The dataset analysis modules are again divided into multiple submenus covering utilities for quality control as well as calculation and display of key cytometry metrics (e.g. median fluorescence intensity or gate frequencies), visualization of the data via dimensionality reduction or clustering, among others. Clusters are obtained by the four FACSPy-implemented clustering algorithms parc^10^, FlowSOM^2,11^, leiden^12^ and phenograph^13^. Dimensionality reductions (UMAP^14,15^, tSNE, PCA, DiffusionMap^16–18^) can be calculated from within the app. Tools for data visualization also include the calculation and display of expression heatmaps, sample correlation analysis and sample distance calculations. Furthermore, cluster frequencies and cluster distributions in connection with the provided metadata and the marker expression per cluster are provided via interactive plots. The plots can be generated as publication-ready plots using the *matplotlib* backend or rendered in an interactive fashion using *plotly* (**Figure 5B**).

In order to increase the flexibility for plotting, pre-analyzed raw data of the plots can be exported at any time and analyzed or plotted using other third-party software. Furthermore, datasets can be subset, exported and loaded on demand to enable following analyses via the API or using other third-party libraries from other programming languages as described above.

## Discussion

As cytometry datasets increase in complexity and in sample size, data analysis becomes an increasingly complex task. Strategies for **automated data handling**, **cell type identification** and **-quantification** and **downstream analysis** are therefore crucial for the efficient analysis of highly dimensional datasets^19–22^.

Automation is an increasingly requested feature in data analysis. While the automatization of data analysis not only increases the speed of the analysis process, it also reduces the amount of manual, and therefore subjective, user input and therefore improves reproducibility of the analysis. In order to facilitate automation of cytometry analysis, we implemented algorithmic solutions for two key steps: data transformation and gating of cytometry events.

As described above and in the **Methods** section, data transformation is automated using curve-sketching of the channel histogram, enabling the user to exert the asinh transformation, which is widely used in cytometry analysis.

In order to automate the gating of cytometry events, we implemented a supervised cell gating approach that uses pre-defined gating strategies that are subsequently learned by the machine and applied on previously ungated samples. Supervised methods are a good choice when inter-sample variability is low and sample sizes are large, for example in diagnostic settings that are usually aimed at quantifying known cell types. We first benchmarked different machine learning algorithms for supervised cell gating according to manually gated ground truth data and found that tree-based methods outperformed other approaches for automated cell quantification with respect to accuracy, speed and memory efficiency. This is intuitive, as the process of decision trees instinctively mimics the decision making in a hierarchical, biaxial gating strategy. The described functionality was implemented in FACSPy by the *supervisedGating* class and allows for the automated training, hyperparameter tuning and prediction of ungated cells.

On the other hand, the curiosity driven nature of basic science requires approaches for unbiased cell classification to detect new phenotypes and cell heterogeneity. In order to account for this type of cell identification, we implemented a semi-supervised gating approach where the user only provides a characterization of the populations of interest in the form of marker expression combinations. The use of unsupervised clustering facilitates the grouping of cells into communities based on all marker proteins, not just the markers used in a biaxial gating plot. The obvious difference to classical gating is that a cell is not classified based on its individual marker expression in a two-dimensional biaxial gate but rather based on the membership of a cell to a multi-dimensionally defined community fulfilling cell-type-specific criteria. Expectedly, given the dissimilarities of both approaches, we found significant differences comparing the semi-supervised assignment of cell type labels to manual gating strategies. While this discrepancy has to be kept in mind, we note that comparable manual gating strategies of the same cell type and gates set for continuous marker expressions also resulted in considerably different gating results **(Extended Data Figure 3 and Supplementary Figure S14)**. Taken together, we propose the usage of unsupervised clustering with concurrent semi-supervised cell annotation for the unbiased and automated gating of cytometry events. We benchmarked different clustering algorithms and found that graph-based algorithms such as PARC, leiden and phenograph were equally performant compared to the FlowSOM algorithm, which is based on a self-organizing map.

We introduce FACSPy as an API for an end-to-end cytometry analysis platform combining key techniques of cytometry analysis in a ready-to-use manner. In addition, we developed FACSPyUI that represents a desktop app interface of FACSPy, enabling users to use the API without prior programming experience.

FACSPy is embedded into the scverse ecosystem^8^ and leverages the *anndata*^23^ object, which has experienced immense popularity among single cell analysis enthusiasts since its creation. The advantages over conventional tabular data are numerous, but the widespread use and flexible data annotation with manifold data types makes it an ideal platform for any single cell-based analysis. We extend on similar approaches^5,24,25^ by providing an end-to-end analysis platform covering cytometry-specific analysis and plotting methods without the use of specialized data structures.

One practical problem in single-cell-analysis is the frequent data subsetting of individual populations of interest with concurrent analysis thereof. We leverage the above-mentioned flexibility of the *anndata* object and limit analysis steps to a user-defined population of interest that is provided as a *gate* parameter to the respective function calls. Corresponding data are automatically added to the *anndata* object in a correctly re-indexed fashion, thereby leaving cells not belonging to the population of interest untouched but still available. This approach completely removes the necessity for manual data subsetting and keeping multiple copies and data subsets of the data in one analysis session.

Finally, we sought to apply the API to a desktop-app based interface, in order to enable users without programming knowledge to leverage FACSPy. The app transfers the functionality provided by FACSPy to an interactive interface that allows for the rapid and intuitive data exploration of cytometry data. While other similar tools exist^26,27^, we made the deliberate choice of building our application on top of a flexible API, therefore minimizing compatibility and flexibility issues that would arise in a completely standalone application.

## Materials and methods

### Datasets

Three flow cytometry datasets consisted of cell lineage analysis of C57BL/6 bone marrow, peripheral blood and spleen. One of the diagnostic flow cytometry datasets consisted of a T-cell lineage panel of 13 human donors and a second consisted of a CD4 T-cell phenotyping panel of 107 human samples. The CyTOF dataset of the Human Immune Monitoring Center (HIMC) was assembled from 19 samples of human donors (FlowRepository FR-FCM-ZYAJ)^28^. The spectral flow cytometry dataset contained 4 samples from human donors (FlowRepository FR-FCM-Z2QV)^29^. Cytometry panels are listed in **Supplementary table 1–5**.

### Mouse samples

C57BL/6J mice were housed under SPF-conditions with a 12h light/dark cycle, a temperature of 22 ± 2 °C, a humidity of 50-60% and food and water available ad libitum. Blood was drawn by cardiac puncture under general, isofluorane-induced anaesthesia followed by cervical dislocation. Spleens were removed and bone marrow was extracted by flushing with RPMI (Gibco Cat. No. #21875-034) supplemented with 10% heat-inactivated FBS (PAN Biotech #3302/P101102) and 1% GlutaMAX (Gibco #35050-061). Spleens were mechanically disintegrated and filtered using a 40 µm filter. Peripheral blood erythrocytes were lysed by two concurrent cycles of ACK lysis (Lonza #10-548E) for three minutes each at room temperature, followed by centrifugation at 300xg for 5 min. Erythrocytes from the spleen and bone marrow were lysed using ACK lysis for 1 min at room temperature, followed by a centrifugation at 300xg for 5 min. Cells were counted and 10^6^ cells were resuspended in 50 µl FACS buffer (2% FBS, 5 mM EDTA, 0.1% sodium azide) containing the antibody panel **(Supplementary Table 1)**.

### Flow Cytometry

Samples were acquired on a Symphony A3 flow cytometer (BD BioSciences). At least 50.000 live cells per sample were recorded. FCS files were exported (v. 3.1) and gated and compensated using FlowJo (v10.8.0). Gating strategies are supplied in **Extended Data Figure 4-8**.

### Cofactor calculation

The automated cofactor calculation is an algorithm that defines the fluorescence intensity separating positive from negative events using histogram data. The transformation cofactor is then adjusted iteratively, so that this fluorescence intensity value is transformed to approximately 0.88 using the arcsinh function. First, expression data for each channel and sample are collected and transformed using a starting cofactor of 200 for flow cytometry data. Subsequently, a kernel density estimate is computed using convolution (FFT). The resulting histogram is inspected as follows: for control samples, peaks at increasing signal intensity values are detected using the *scipy*^30^ implementation in the function *find_peaks*. The inflection point of the right side of the first peak (i.e. corresponding to background signal) is calculated using the first and second derivative of the histogram function. A tangent line is drawn to the inflection point and the root of the tangent line is used to adjust the cofactor by sinh(root) * cofactor (**Figure 4C**). This process is repeated for n times, where n represents a user-definable iteration number (default setting = 2). For stained samples, peaks are detected as described above. The first two peaks with the highest density are then subset and the left peak is inspected as follows: In the first iteration, the base of the peak is found using the *find_peaks* function of *scipy*. As the left peak represents unstained cells in a stained sample, its prominence can be shallow. Therefore, the drawing of a tangent line often resulted in a significant overestimation of the cofactor adjustment (data not shown). After adjusting the cofactor using sinh(peak_base)*cofactor, the iterative process using the tangent line at the inflection point as described above is computed for n-1 times where n represents a user-definable iteration number. This approach allows for a rapid detection of the relevant cofactors without the need for testing for the optimal cofactor (as implemented in e.g. FlowVS^31^ or FlowTrans^32^). Cofactors are calculated for each sample and channel separately and the median of all calculated cofactors per channel is used for the final data transformation.

### Supervised gating – implementation details

Gated samples are defined by the user. The corresponding gate matrix as well as the fluorescence intensity values are subsequently scaled and sampled (see below) as per the user’s request. A classifier (DecisionTree, ExtraTree, RandomForest, ExtraTrees, MLP or KNN) is trained with the scikit-learn syntax and implementation, using the fluorescence intensity values as input and the gate matrix as output. The trained classifier is finally used to predict the gate matrix of ungated samples.

### Supervised gating – data sampling

Data sampling is used to avoid overfitting of the classifier and to match the respective class-counts. Cell classification in cytometry is a multi-label machine learning problem where cells are included or excluded in n different gates (labels). However, individual gates can be dependent on each other (e.g. parent gates). While this is per se not problematic, size matching of the individual label classes becomes unfeasible, as increasing the number of a cell population also increases the number of examples of the parent gate labels. In order to circumvent that problem, we developed an algorithm that first binarizes the gating strategy and transforms the multi-label classification problem into a single-label classification. The training dataset is then sampled for each individual gating strategy rather than individual gates, where randomized removal of examples is used for downsampling and the application of gaussian noise is used as a technique for data augmentation. This technique allowed a noticeable increase in the recall score of very rare gates (**Supplementary Figures 1B–7B**), while keeping the F1-metric comparable for more frequent gates (data not shown).

The gating strategy is stored as a Boolean sparse matrix g x c where g denotes the gates and c denotes the cells. The matrix is then transformed into its integer representation where every row (sequence of Booleans) is treated as a bit sequence and is converted into a 64bit integer by calculating the dot-product of the binary sequence with an array storing the powers of two in a decrementing order. As python does not readily support integers greater than 64 bits, gating strategies with more than 64 gates get split into n matrices of shape (g / 64) x c, where g denotes the gates and c denotes the cells. Each matrix is then transformed as described above and combined in an integer array. The bit representation integers are then mapped to unique integers in order to allow classification of gating strategies with more than 64 gates, which results in two or more bit representing integers per cell.

The sampling strategy denotes how many cells per individual gate are to be sampled. First, all user defined gates are reconstructed from the gating strategies and constructed into their binary representations as described above. Importantly, upsampling is only provided for user-defined gates. Cells that do not match any user-defined gating strategy (e.g. cells that are present in two or more gates at the same time) are not excluded but also not upsampled. While these non-predefined gating strategies are often heterogeneous and frequent (up to 5-20x of user-defined gating strategies), their respective cell count per gate were generally low (<<1% of all cells, data not shown). For each class, cell numbers are matched to the sampling strategy by either randomly selecting a subset of cells for downsampling or the use of gaussian noise or the SMOTE algorithms^33^ for upsampling.

### Classifier benchmark and hyperparameter tuning

To evaluate the accuracy of supervised cell classification, different datasets were assembled using FACSPy (**Table 1**). The compensated expression data were arcsinh-transformed using manually determined cofactors and subsequently Z-scaled using the StandardScaler implementation of *scikit-learn*^34^. Expression data served as input X, whereas the class labels were provided as a Boolean matrix of shape g x c, where g denotes the gates and c denotes the cells.

In a first experiment, 26 classifiers as implemented in the *scikit-learn* library were selected, of which 11 were excluded from further experiments due to an error if an empty class was present. The remaining 15 classifiers were subjected to our benchmark, among them tree-based methods (Random Forest, Decision Tree, Extra Trees, Extra Tree), neural networks (multi-layer perceptron), naïve bayes based classifiers and linear classifiers (**Figure 1 and Extended Data Figure 2**). In order to test the performance of the described classifiers, one sample of each dataset was reserved as a validation dataset and 10.000 events of the remaining n-1 samples were randomly selected, split into a training- and test-set (ratio: 0.1) and used for training. This process was repeated for all samples of a given dataset. Finally, the trained classifiers were used to predict the gate matrix of the validation dataset and the F1-score between the predictions and the original gating strategy of the validation sample was computed for each column g of the gate matrix separately. The top six algorithms (Random Forest Classifier, Decision Tree Classifier, Extra Trees Classifier, Extra Tree Classifier, multi-layer-perceptron and the k-nearest neighbor classifier were selected and used for further characterization. In order to test the effect of increasing sample sizes, the classifier was trained on n-1 samples as described above with ascending training sizes, reaching from 50 to 500,000 cells. The F1-score was calculated again on all gates of the classified validation set of the remaining sample. Hyperparameter tuning was conducted using the RandomizedHalvingSearch implementation of *scikit-learn*, where two thirds of possible parameter combinations were removed at each iteration (for a detailed list of tested parameters refer to the code documented). The best parameters were obtained using 5-fold cross validation. These parameters were then used to repeat the analysis as described above. Memory consumption was measured using the *memory-profiler* package. Analysis time was quantified using the built-in *time* module in python. For the visualization in **Extended Figure 2C**, the circle size was normalized to 0, corresponding to the minimum F1-score per classifier and dataset, and 1, corresponding to an F1-score of 1 and plotted for the untuned and tuned classifier. In order to preserve the global change in F1-score, the semicircles were colored according to the median F1-score for each the untuned and tuned classifier.

### Semi-supervised gating – implementation details

The gating strategy for the semi-supervised gating is supplied by the user in the form of a population name, the parent population and markers describing the population of interest (positive markers, negative markers or marker intensity of ordinal scale from low, int and high). Importantly, this user-supplied gating strategy is built around logic and does not rely on fixed signal intensity cutoffs to be applicable. The cells of the parent population are subsequently clustered using one of the implemented clustering algorithms (leiden, parc, phenograph or FlowSOM). For each cluster, the marker expression is evaluated and compared with the user-supplied markers specific for the respective population. The matching clusters are then assigned the name of the population of interest. Marker positivity is defined of a median expression of the respective marker per cluster of above 0.88 (arcsinh transformed space), where negativity is defined as a median expression below that value. The sensitivity parameter can be used to adjust the positivity threshold where the threshold is defined as arcsinh(1) – (0.1*log10(sensitivity)). The cutoffs for low, intermediate and high marker expression are calculated per parent population as quantiles over all cells, which can be further defined by the user (defaults: low: 0-33^rd^ quantile, intermediate: 33^rd^ - 66^th^ quantile, high: 66^th^ quantile - 1).

### Algorithm comparison of semi-supervised gating - benchmark

To compare the performance of different algorithms, datasets were first assembled and arcsinh-transformed using manually defined cofactors in FACSPy. The transformed data were clustered using the leiden^12^ algorithm as implemented in the scanpy package^9^, the parc algorithm^10^, the phenograph algorithm^13^ or FlowSOM^2^ with a fixed metacluster number of 30. First, the classifiers were run with standard settings (**Figure 3B, Supplementary Figures 8B–13B**) and the jaccard index was computed as the intersection of positively quantified events by manual and semi-supervised gating divided by the size of their union. The effect of cluster resolution as evaluated for the graph-based clustering methods leiden, parc and phenograph **(Supplementary Figures 21-27, A–C**), by repeating the above-described experiment with different parameters for cluster resolution. For FlowSOM clustering, the number of metaclusters was tested **(Supplementary Figures 21–27, D**). Memory consumption was measured using the memory-profiler package. Analysis time was quantified using the built-in time module in python (**Figure 3C, Supplementary Figures 21-27**).

### FACSPy design and architecture

The FACSPy architecture is built around the AnnData^8,23^ object, which stores user-defined metadata (obs), panel information (var), an optional workspace containing the gating information (obsm) and other associated data (uns) together with the cytometry expression data (layers). While there are no strictly mandatory supplementary files, metadata as well as additional panel information can be provided using tabular data. First, FCS files are read-in and compensated using a provided workspace^5^, a compensation matrix extracted from the FCS file, or a separate matrix supplied by the user using the *flowio* (https://github.com/whitews/FlowIO) and *flowkit*^5^ package. The data are subsequently annotated using the provided metadata, gated, and saved as an .h5ad file. This file can be read using different anndata implementations in R, python and Julia, allowing for the use of analysis steps which are not currently implemented in python (**Figure 4A**). FACSPy is deliberately designed to store analyses for multiple populations within the same *anndata* object (**Figure 4B**), allowing for comparative and parallel analysis of different populations without the need for data subsets. Populations of interest are defined using manual predefined gating strategies or leveraging the API-provided supervised or semi-supervised approaches (**Figure 4D**). The data can be transformed using the hyperlog, logicle, log or arcsinh transformation of which the latter requires channel-specific cofactors for optimal performance^31,35,36^. These cofactors can either be supplied as determined manually or calculated automatically from the channel histogram (**Figure 4C).** The transformed data are subsequently stored as a different layer within the AnnData object, enabling parallel analysis on compensated and transformed data. The *tools* module offers distinct analysis capabilities for data integration, dimensionality reduction such as UMAP^14,15^, TSNE^37,38^, PCA and DiffusionMaps^16–18^ and differential expression, among others (**Figure 4E and Extended Data Table 2**). A dedicated *plotting* module is used for data visualization and is implemented to fulfil the requirements of plotting data from different populations and different data origins (e.g. compensated or transformed data; **Figure 4F and Extended Data Table 1**). For each function, a *gate* has to be provided in order to specify the population of interest to be analyzed. This gate can either be supplied as a singular string, pointing to the end of a gating strategy (e.g. “T_cells” being short for root/singlets/live/T_cells) or as a partial or full gating path. A dedicated settings class allows, among other features, for the definition of a default gate for rapid concurrent analysis steps and the definition of gate aliases to increase the user-friendliness and replace long gating paths. In order to build on pre-existing syntax and infrastructure, FACSPy leverages the scanpy^9^ notation for functions which aims to ensure a quick understanding of the API for users already familiar with single cell analysis in python using scanpy.

## Code availability

The FACSPy API and the FACSPyUI are publicly available at https://github.com/rgb-lab/FACSPy and https://github.com/rgb-lab/FACSPyUI, respectively. The code to reproduce all figures and data generated in this paper is publicly available at https://github.com/rgb-lab/CytoGateBenchmark. All data generated have been deposited at https://n-hackert.github.io/facspy_websupps and can be visualized interactively.

## Acknowledgements

T.E. was supported by an MD/PhD fellowship from the Medical Faculty of Heidelberg. This work was supported by grants to R.G.-B. from Deutsche Forschungsgemeinschaft (DFG, GR 5979/2-1, 517717827), Else Kröner-Fresenius-Stiftung (2022_EKEA.72), state of Baden-Wuerttemberg within the Centers for Personalized Medicine Baden-Wuerttemberg (ZPM) and a research grant from the German Society for Rheumatology (DGRh). Figures 1A, 4A and 5A were created with BioRender.com. The authors acknowledge support by the state of Baden-Württemberg through bwHPC and the German Research Foundation (DFG) through grant INST 35/1597-1 FUGG. We thank Andrei Stapran, Richard Huth and Caroline Pabst for extensive beta-testing of FACSPy during development.

## Supplementary Tables

**Extended Data Table 1:**
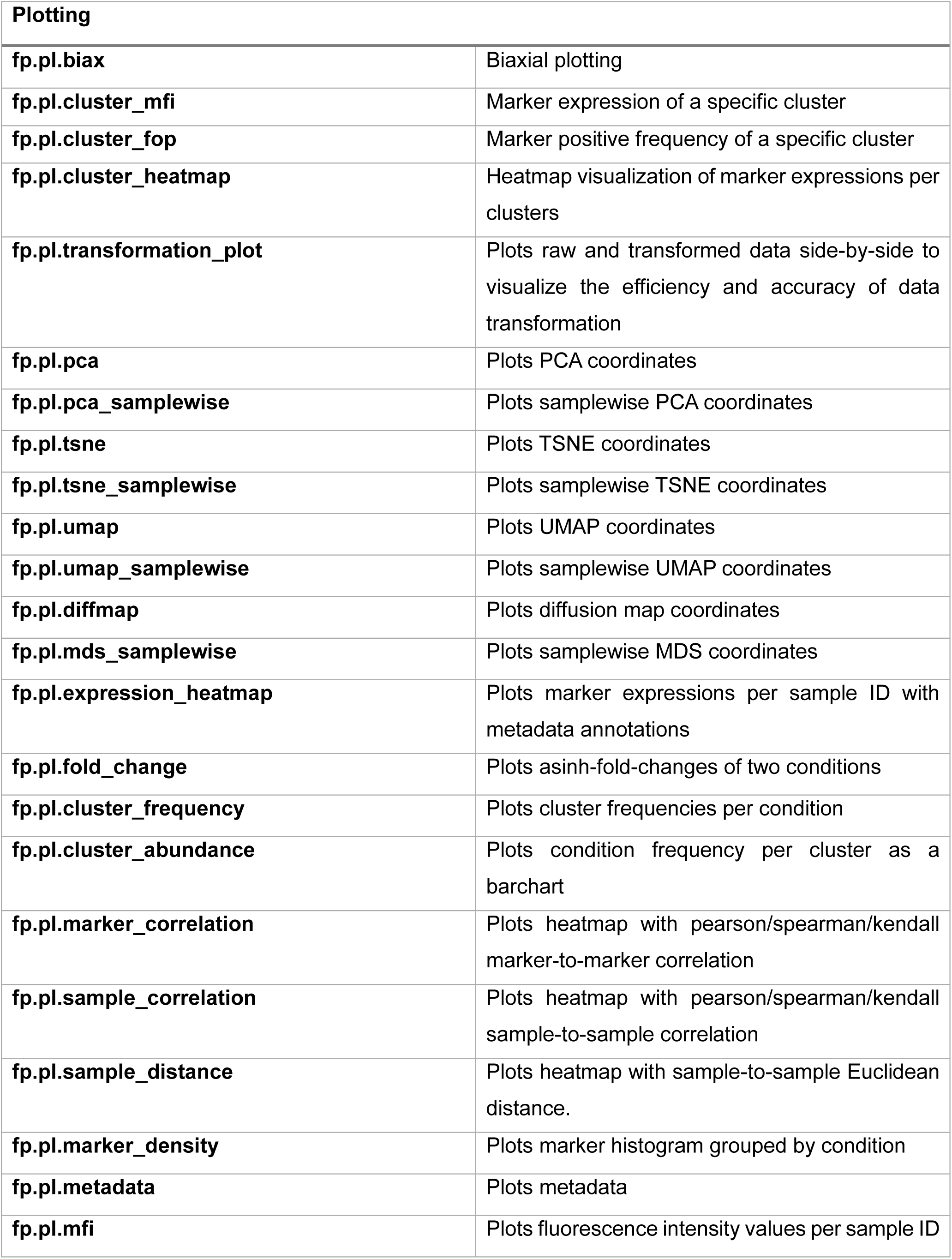
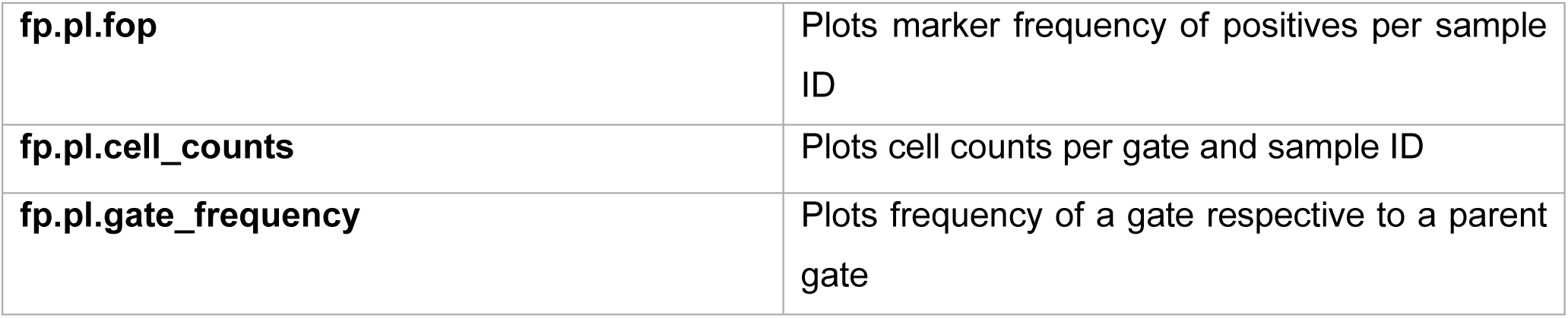
Overview of the plotting functionality of FACSPy.

**Extended Data Table 2:** Overview of calculation properties by FACSPy.

| Tools (.tl) |  |
| --- | --- |
| **fp.tl.pca** | Calculates Principal Component Analysis |
| **fp.tl.pca_samplewise** | Calculates Principal Component Analysis sample-to-sample |
| **fp.tl.umap** | Calculates UMAP |
| **fp.tl.umap_samplewise** | Calculates UMAP sample-to-sample |
| **fp.tl.tsne** | Calculates TSNE |
| **fp.tl.tsne_samplewise** | Calculates TSNE sample-to-sample |
| **fp.tl.diffmap** | Calculates DiffusionMaps |
| **fp.tl.mds_samplewise** | Calculates Multidimensional scaling sample-to-sample |
| **fp.tl.neighbors** | Calculates neighbors |
| **fp.tl.parc** | Calculates PARC clustering |
| **fp.tl.flowsom** | Calculates flowsom clustering |
| **fp.tl.leiden** | Calculates leiden clustering |
| **fp.tl.phenograph** | Calculates phenograph clustering |
| **fp.tl.fop** | Calculates frequency of positive events per marker |
| **fp.tl.mfi** | Calculates median/mean fluorescence intensity per marker |
| **fp.tl.gate_frequencies** | Calculates frequencies of gate-positive events |
| **fp.tl.scanorama** | Computes scanorama integrated embedding |
| **fp.tl.harmony** | Computes harmony integrated embedding |

---

**Extended Data Figure 1:**
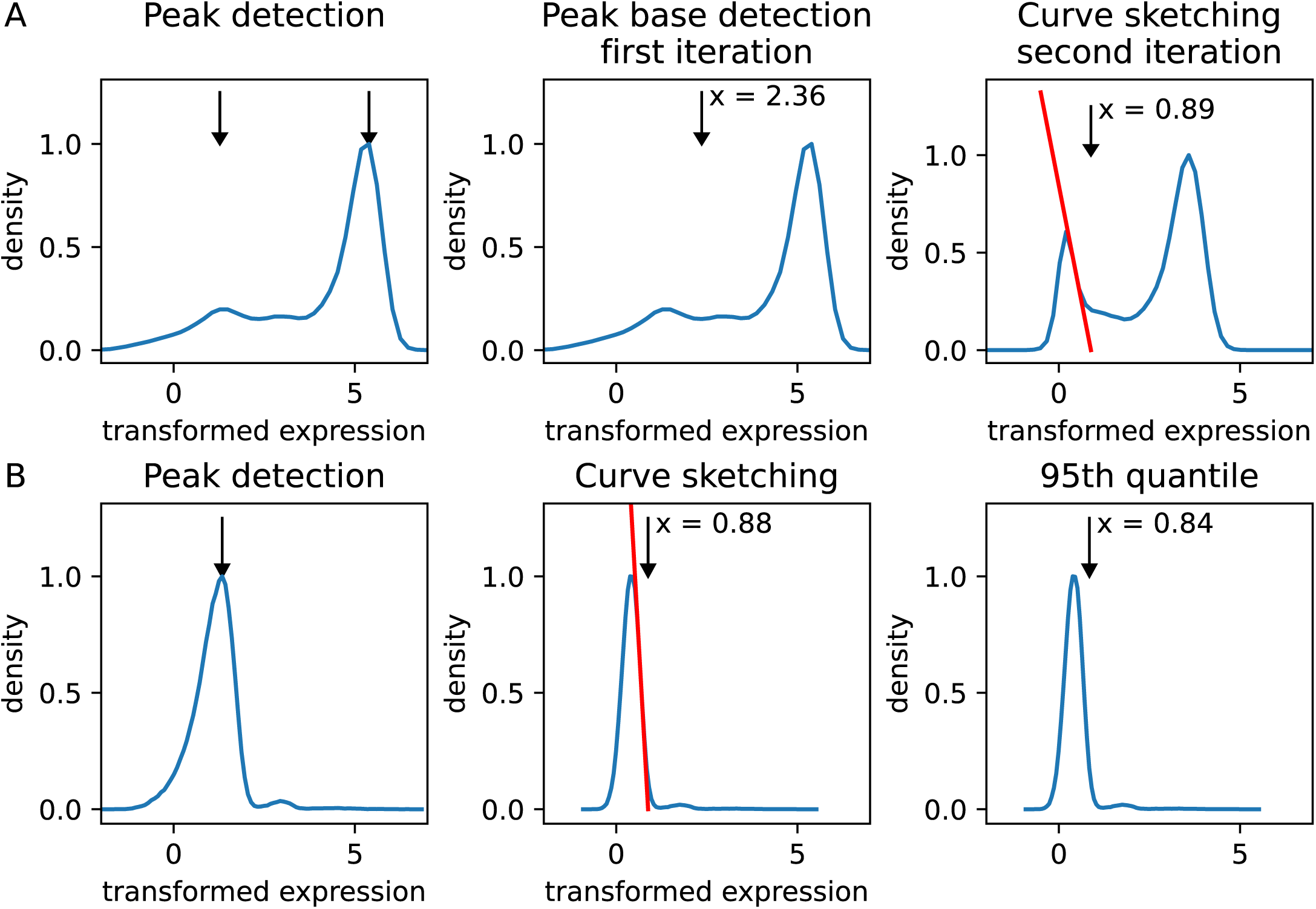
Automated cofactor detection. **A** Shown is a marker histogram of a stained flow cytometry sample. First, peaks are detected (left) and the base of the left peak is determined (mid). A tangent line is drawn to the turning point of the left peak and the root is used to adjust the cofactor (see Methods). This process is repeated using user-defined iterations to converge at ∼0.88. **B** Shown is a marker histogram of an unstained flow cytometry sample. First peaks are detected (left) and a tangent line is drawn to the left peak (middle). The root of the tangent line is used to adjust the cofactor as in a. In addition, the 95th quantile of the control sample is calculated. The final cofactor is then calculated as the mean of both methods.

**Extended Data Figure 2:**
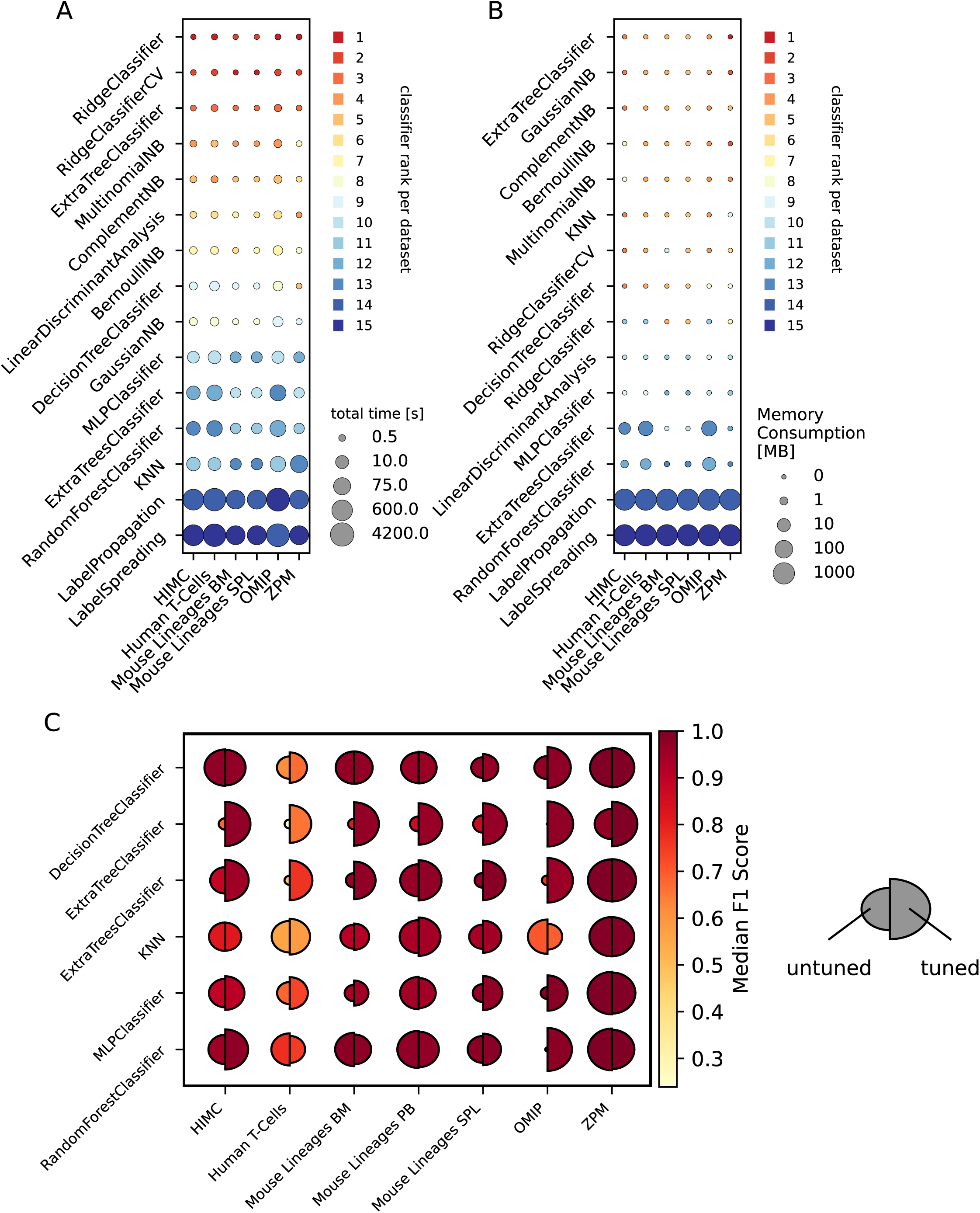
Benchmarking machine learning algorithms for cytometry data analysis. **A** Time comparison. Classifiers were subjected to the benchmark as described in Figure 1. The time for training and predicting the ungated sample was measured. The circle size indicates the time needed by the classifier, while the color code describes the relative rank of the classifier to all other algorithms for this indicated dataset. **B** Memory consumption. Classifiers were trained as described in Figure 1. The memory consumption during training was measured. The circle size indicates the consumed memory during training while the color encodes the relative rank of the indicated classifier among all algorithms used for the respective dataset. **C** Effect of hyperparameter tuning. Classifiers were either left untuned (left semicircle) or were tuned using the given training set and an internal test set. The tuned classifiers were subsequently subjected to the same training and validation procedure as described in Figure 1. For a detailed description on the circle sizes refer to the **Methods**. The color encodes the median F1-score for the indicated classifiers on the respective dataset.

**Extended Data Figure 3:**
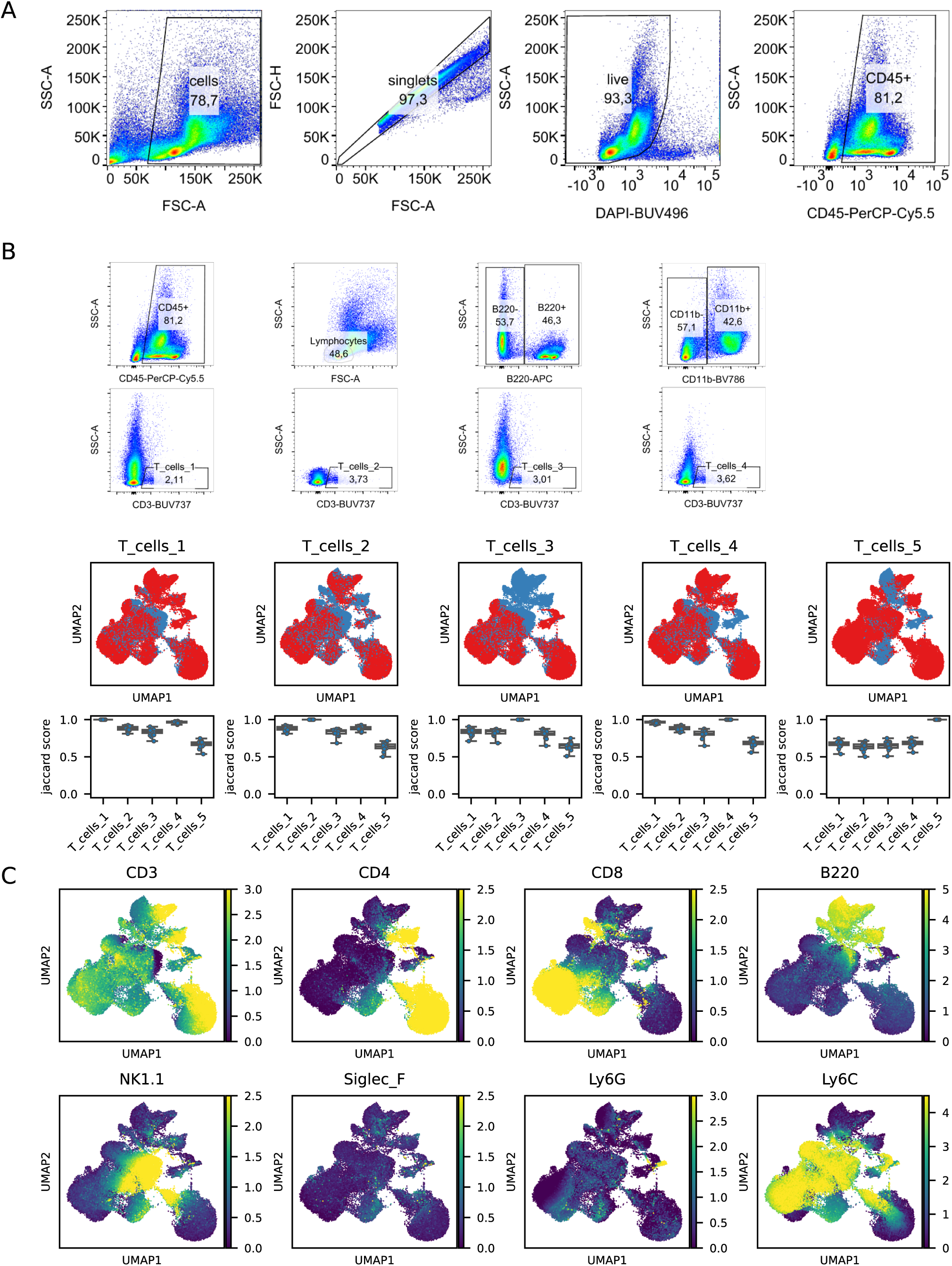
Comparability of differently drawn manual gates. **A** Gating strategy. **B Top**: Four different gates for T-cell gating were drawn (T_cells_1-4). T_cells_1 represents the single CD3-positivity gating while T_cells_2-4 used exclusion of granulocytes (second panel), B cells (third panel) or myeloid cells (fourth panel) before gating T-cells. The final T_cells gate is an exact copy between the four gating strategies. T_cells_5 denotes the classification by semi-supervised gating which used CD3 positivity and SSC low as input parameters to classify T-cells. **Mid**: UMAP representation of all cells belonging to one of the five gating stragies colored by the positive cells as identified by the respective gating (red: positive, blue: negative). Note that contaminating cell populations (e.g. CD3dim, B220+ and CD3dim, Ly6G+) cells were excluded by the semi-supervised approach and only in some exclusion gating strategies, while being included by the T_cells_1 gating. Similarly, CD3+ clusters are identified completely by the semi-supervised gating while the manual gating strategies often labelled cells as negative in the same cluster of cells (bottom left UMAP area). **Bottom**: Jaccard indices quantifying the overlap of identified T-cells of the different gating strategies. **C** UMAP representation as in B, colored by the indicated marker expressions.

**Extended Data Figure 4:**
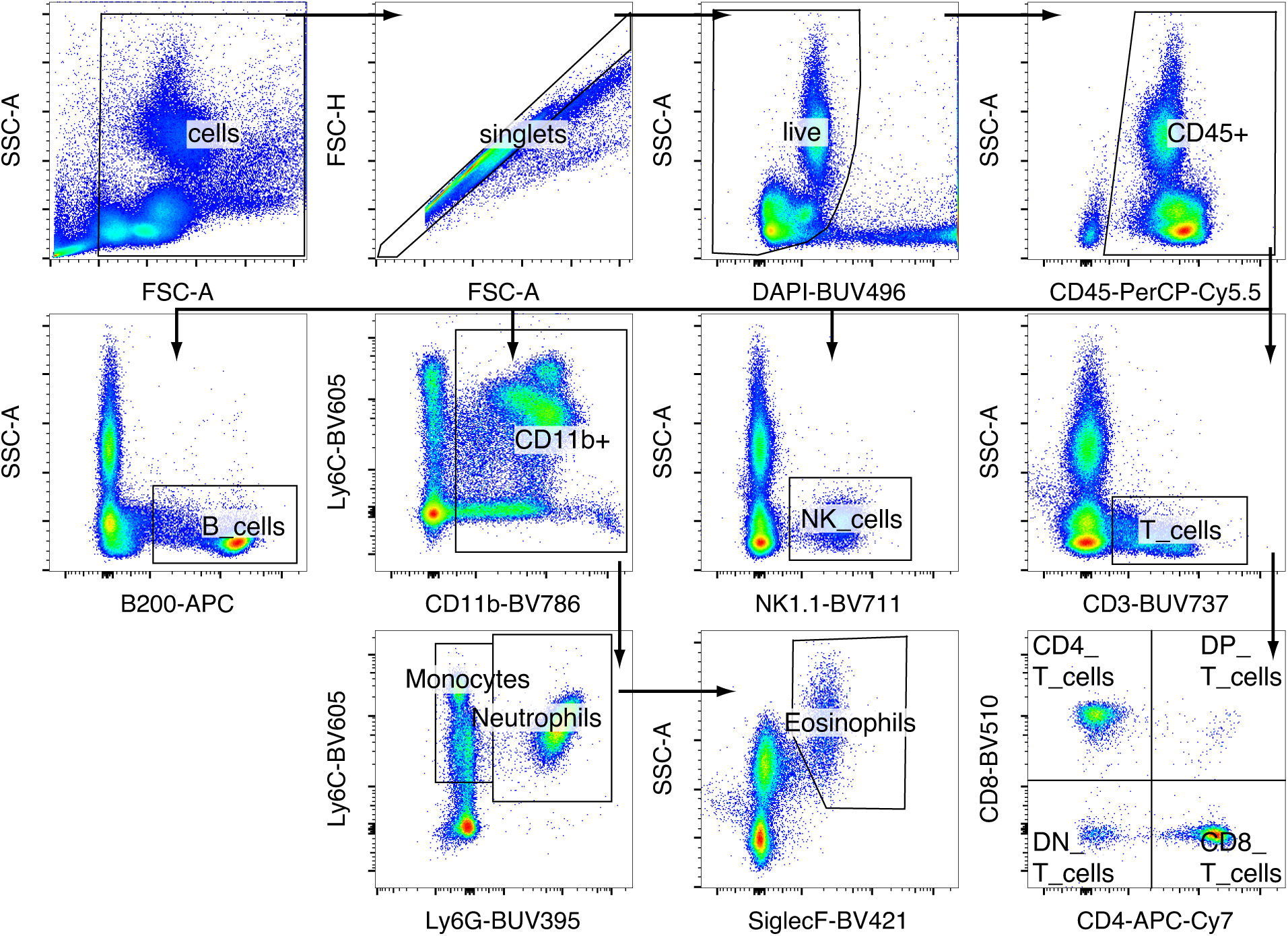
Gating strategy mouse lineages (dataset 1-3) Shown is the gating strategy used for datasets encompassing the mouse lineage data (datasets 1-3, compare Table 1). Shown is a sample from peripheral blood, the gating strategies for bone marrow and spleen were equivalent.

**Extended Data Figure 5:**
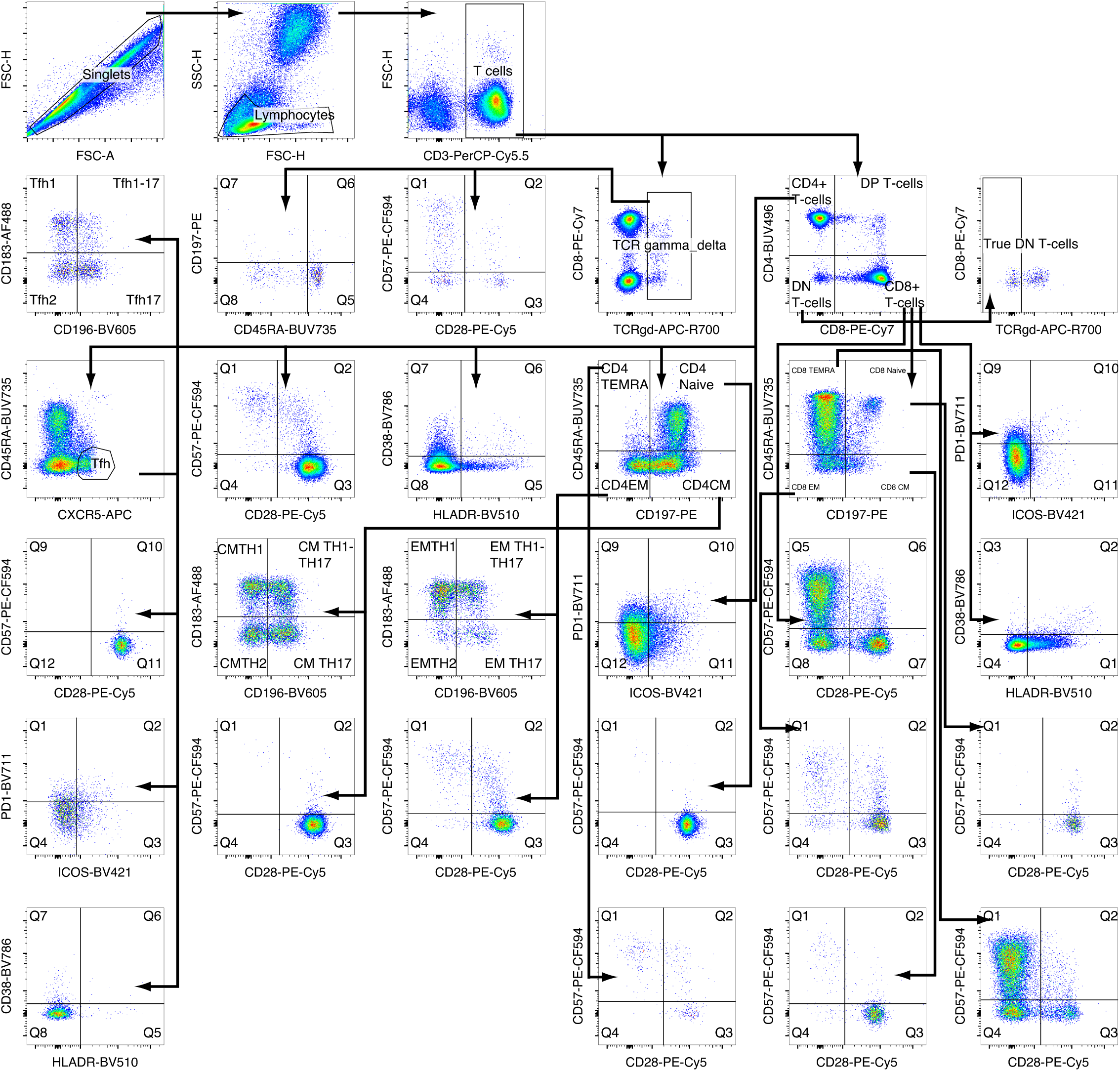
Gating strategy human T-cells (dataset 4) Shown is the gating strategy used for dataset 4 (compare Table 1)

**Extended Data Figure 6:**
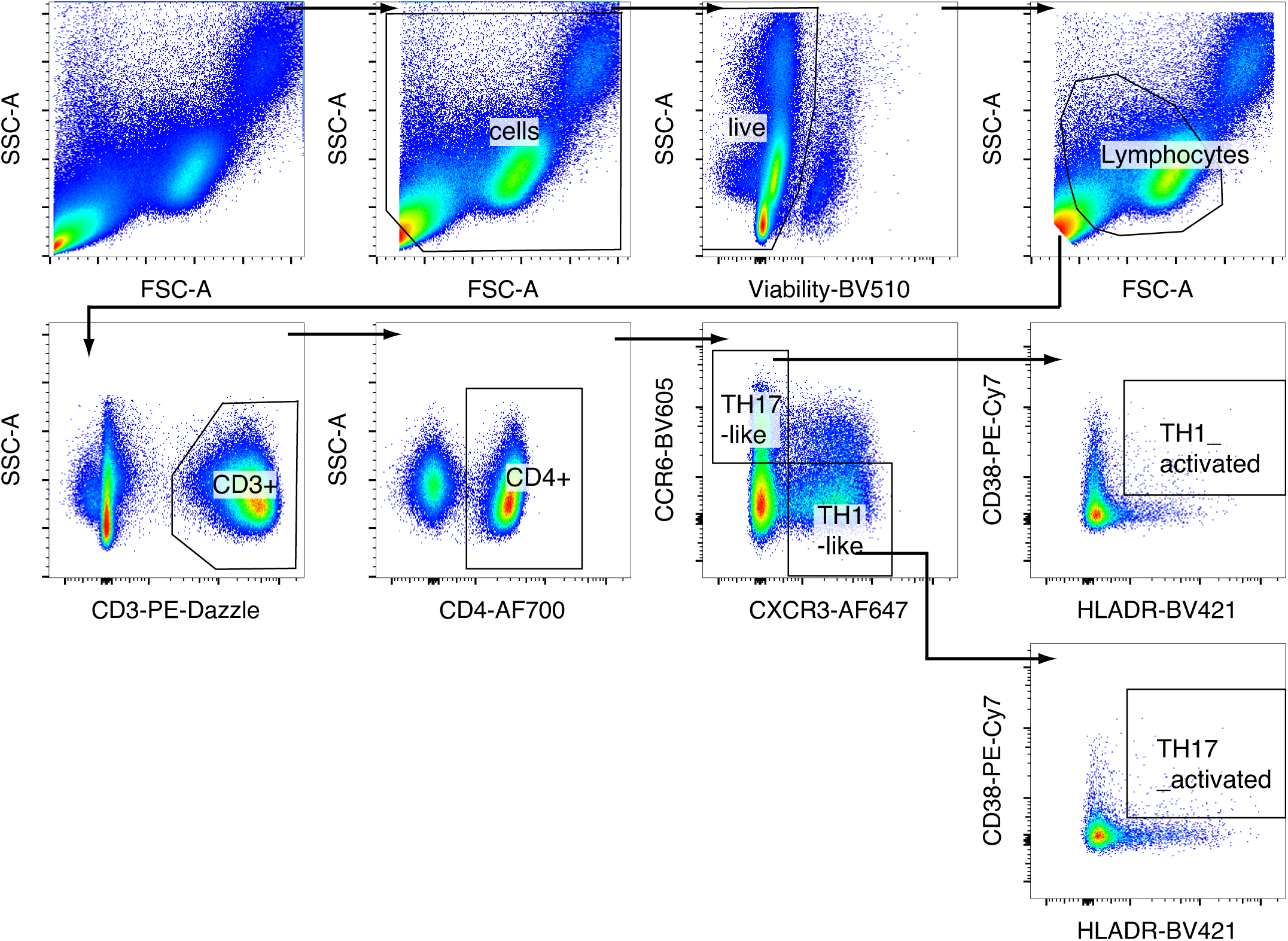
Gating strategy dataset 5. Shown is the gating strategy used for dataset 5 (compare Table 1).

**Extended Data Figure 7:**
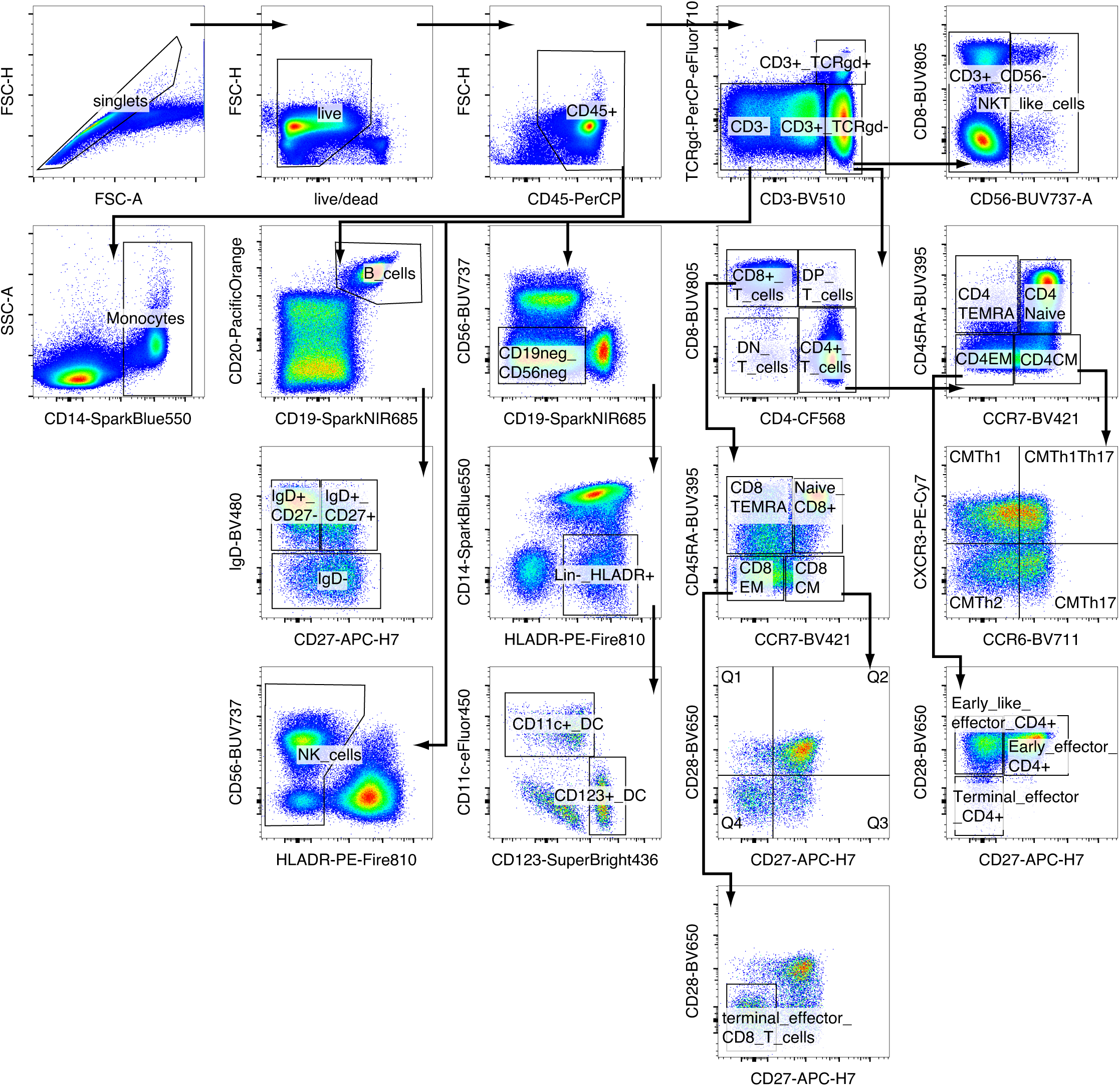
Gating strategy dataset 6. Shown is the gating strategy used for dataset 6 (compare Table 1).

**Extended Data Figure 8:**
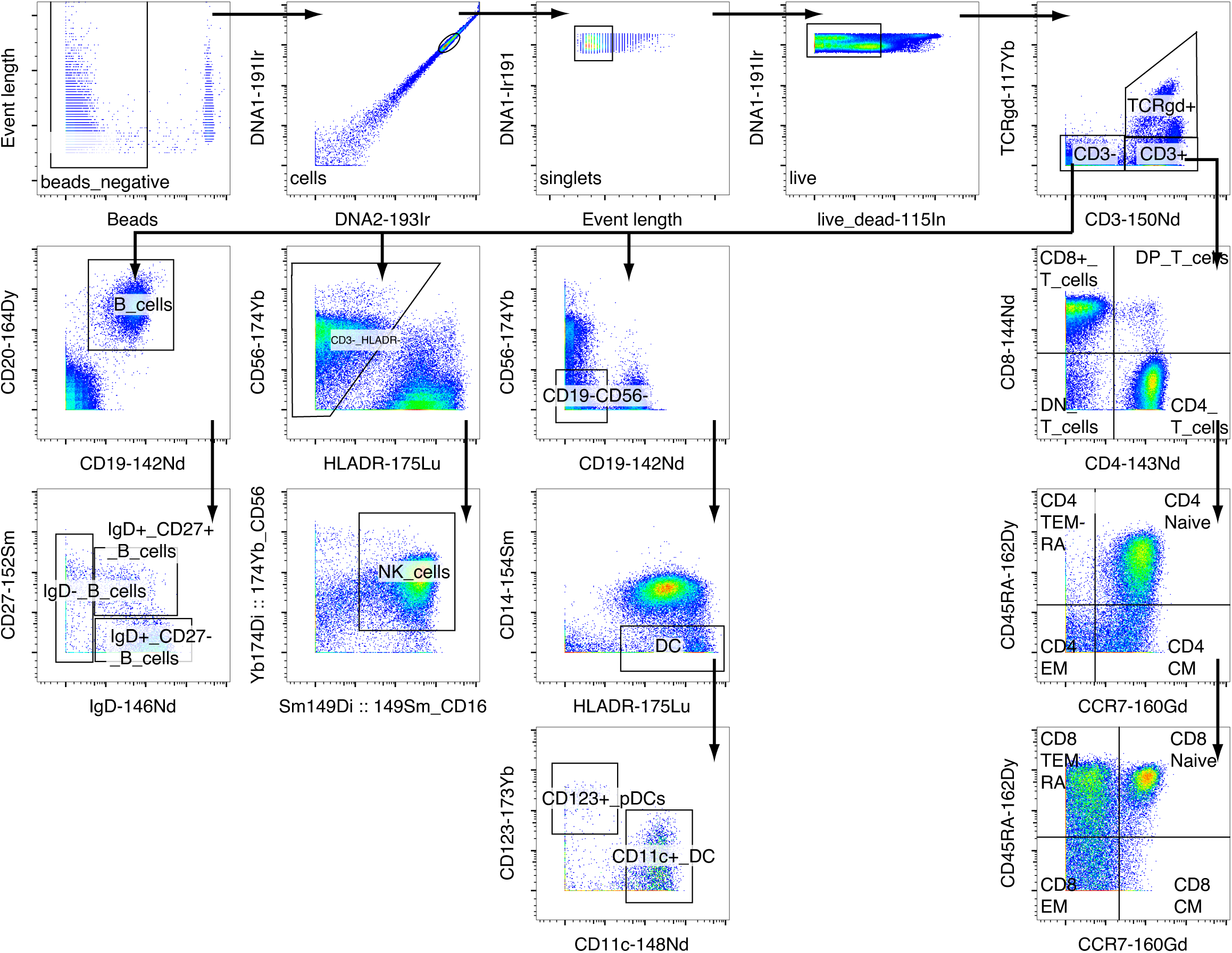
Gating strategy dataset 7. Shown is the gating strategy used for dataset 7 (compare Table 1).

