## Supplementary Material for "Accurate, fast and memory efficient quantification of immune cell phenotypes in cytometry using machine learning"

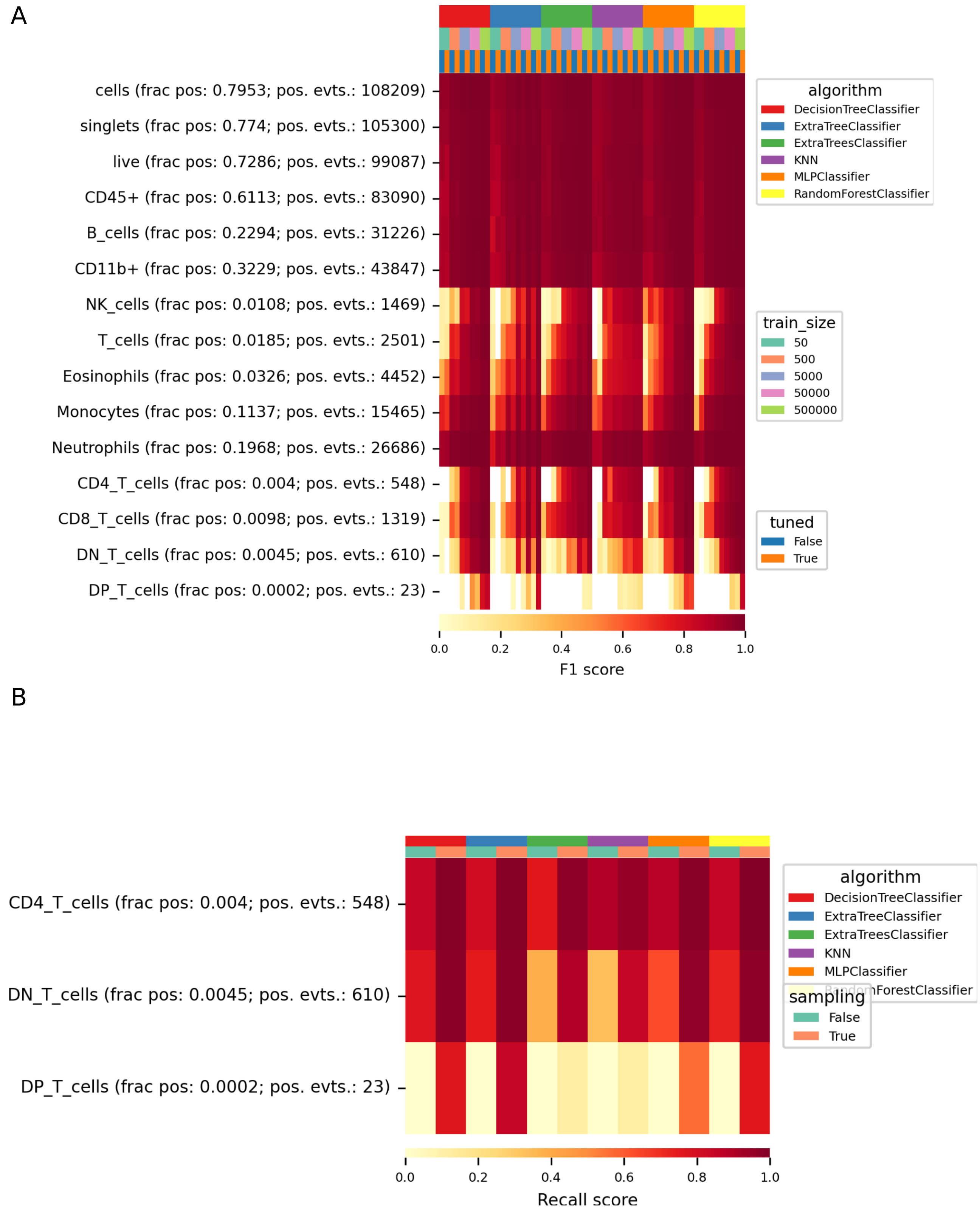


**Supplementary Figure S1: Classifier characterization on a murine flow cytometry dataset.** The dataset consisted of 18 samples of mouse bone marrow cells (flow cytometry; Dataset 1). **A** Classifiers were compared using the indicated train sizes and hyperparameter tuning (top column annotations). Shown are the mean F1 metrics per indicated gate. For each gate, the fraction of positive events (frac pos) as well as the total number of positive events (pos. evts.) are indicated. **B** Recall scores for gates below 0.005% after using a dedicated sampling strategy (sampling: True) compared to random subsampled data. MLP: multi-layer-perceptron, KNN: k-nearest-neighbor classifier

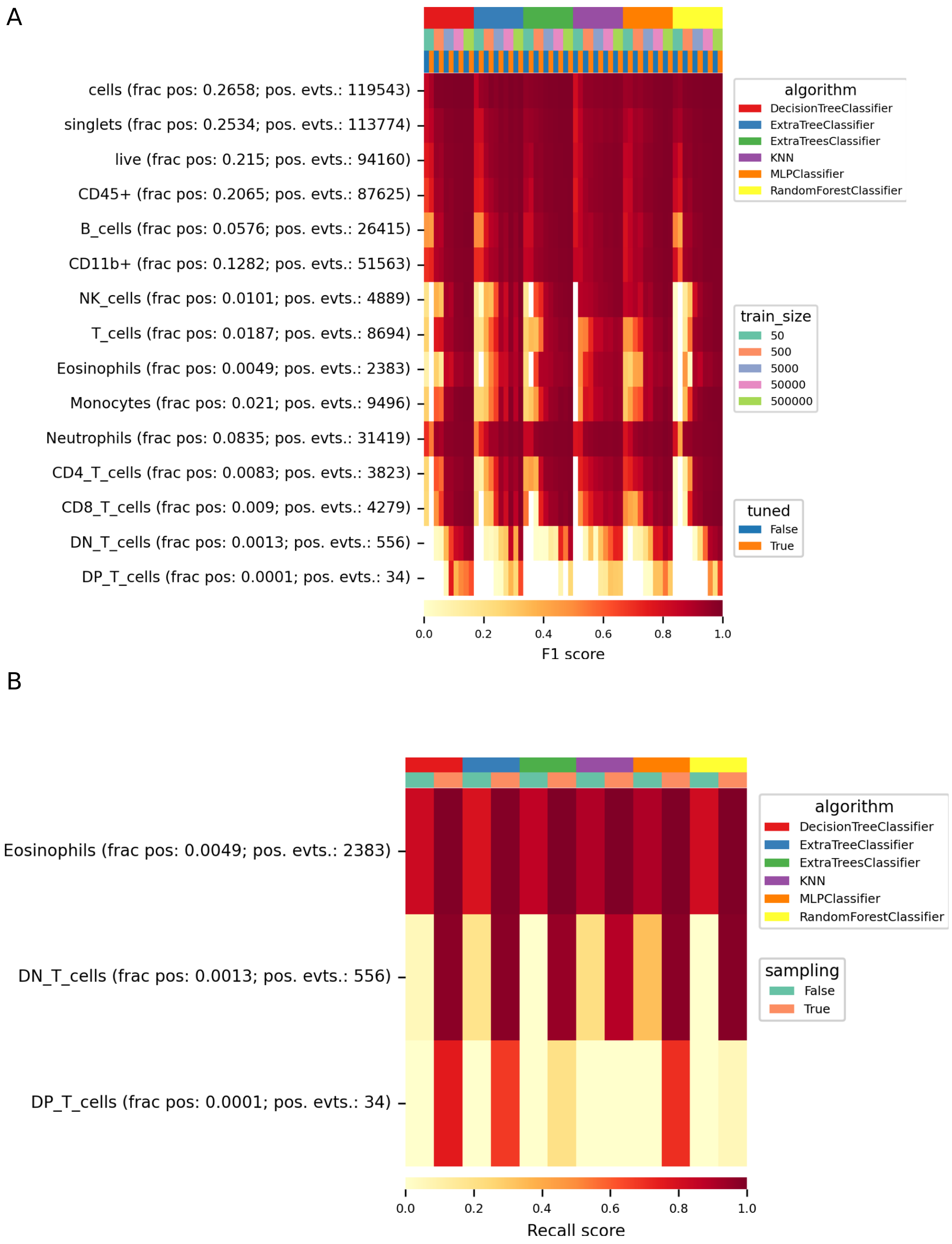


**Supplementary Figure S2: Classifier characterization on a murine flow cytometry dataset.** The dataset consisted of 18 samples of mouse peripheral blood cells (flow cytometry, Dataset 2). **A** Classifiers were compared using the indicated train sizes and hyperparameter tuning (top column annotations). Shown are the mean F1 metrics per indicated gate. For each gate, the fraction of positive events (frac pos) as well as the total number of positive events (pos. evts.) are indicated. **B** Recall scores for gates below 0.005% after using a dedicated sampling strategy (sampling: True) compared to random subsampled data. MLP: multi-layer-perceptron, KNN: k-nearest-neighbor classifier

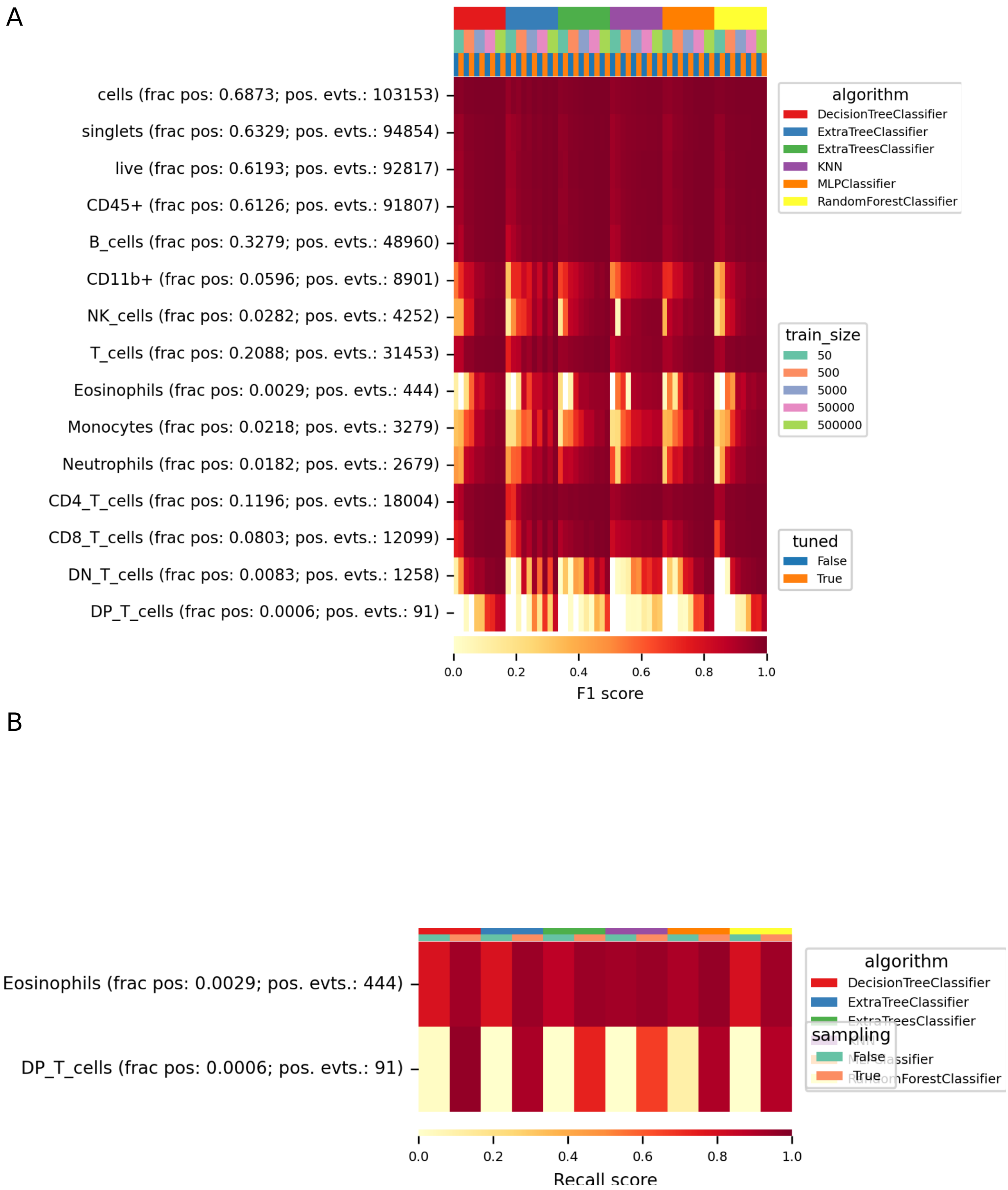


**Supplementary Figure S3: Classifier characterization on a murine flow cytometry dataset.** The dataset consisted of 18 samples of mouse spleen cells (flow cytometry; Dataset 3). **A** Classifiers were compared using the indicated train sizes and hyperparameter tuning (top column annotations). Shown are the mean F1 metrics per indicated gate. For each gate, the fraction of positive events (frac pos) as well as the total number of positive events (pos. evts.) are indicated. **B** Recall scores for gates below 0.005% after using a dedicated sampling strategy (sampling: True) compared to random subsampled data. MLP: multi-layer-perceptron, KNN: k-nearest-neighbor classifier

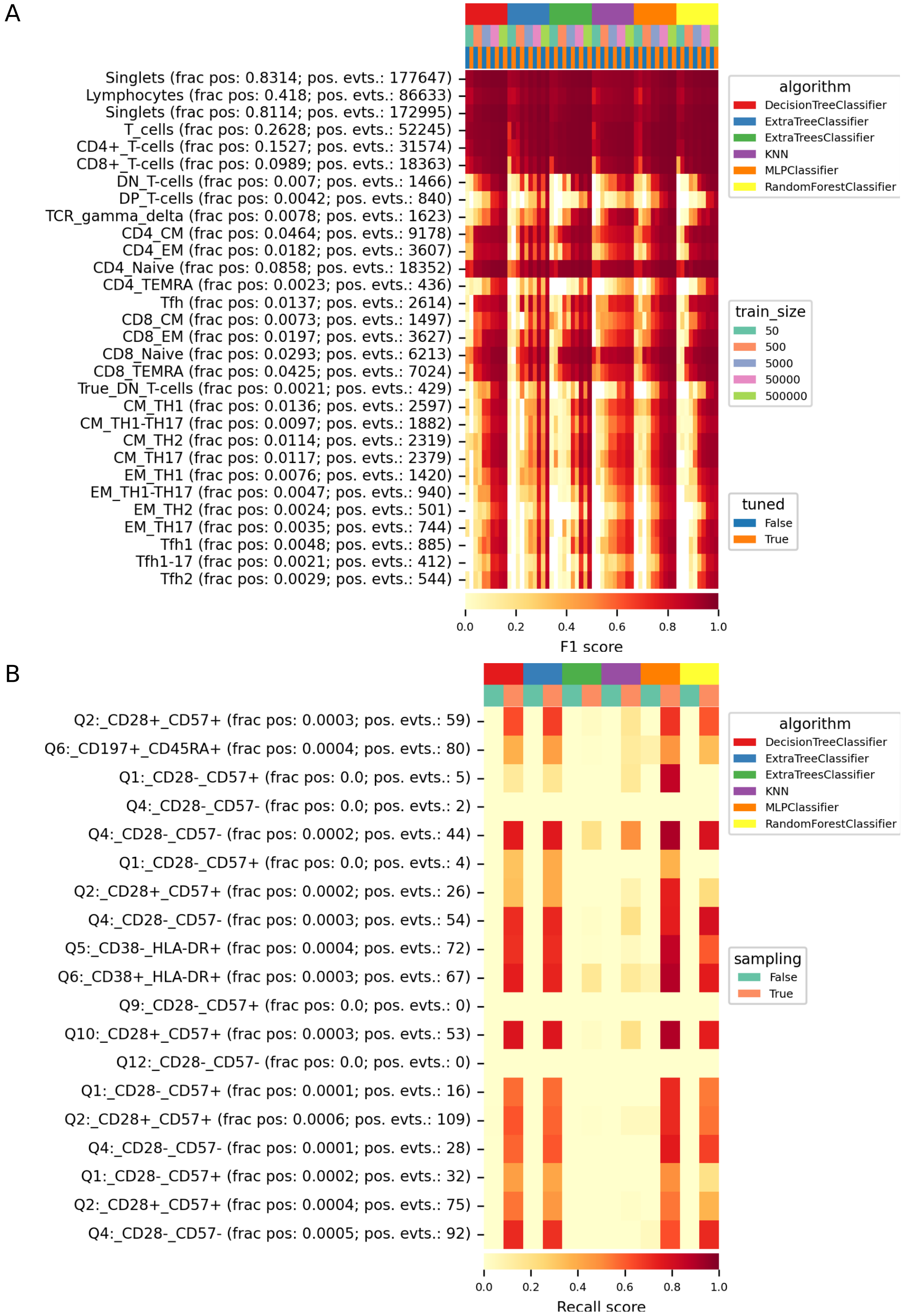


**Supplementary Figure S4: Classifier characterization on a human flow cytometry dataset.** The dataset consisted of 14 samples of human peripheral blood cells (flow cytometry; Dataset 4). **A** Classifiers were compared using the indicated train sizes and hyperparameter tuning (top column annotations). Shown are the mean F1 metrics per indicated gate. For each gate, the fraction of positive events (frac pos) as well as the total number of positive events (pos. evts.) are indicated. **B** Recall scores for gates below 0.005% after using a dedicated sampling strategy (sampling: True) compared to random subsampled data. MLP: multi-layer-perceptron, KNN: k-nearest-neighbor classifier

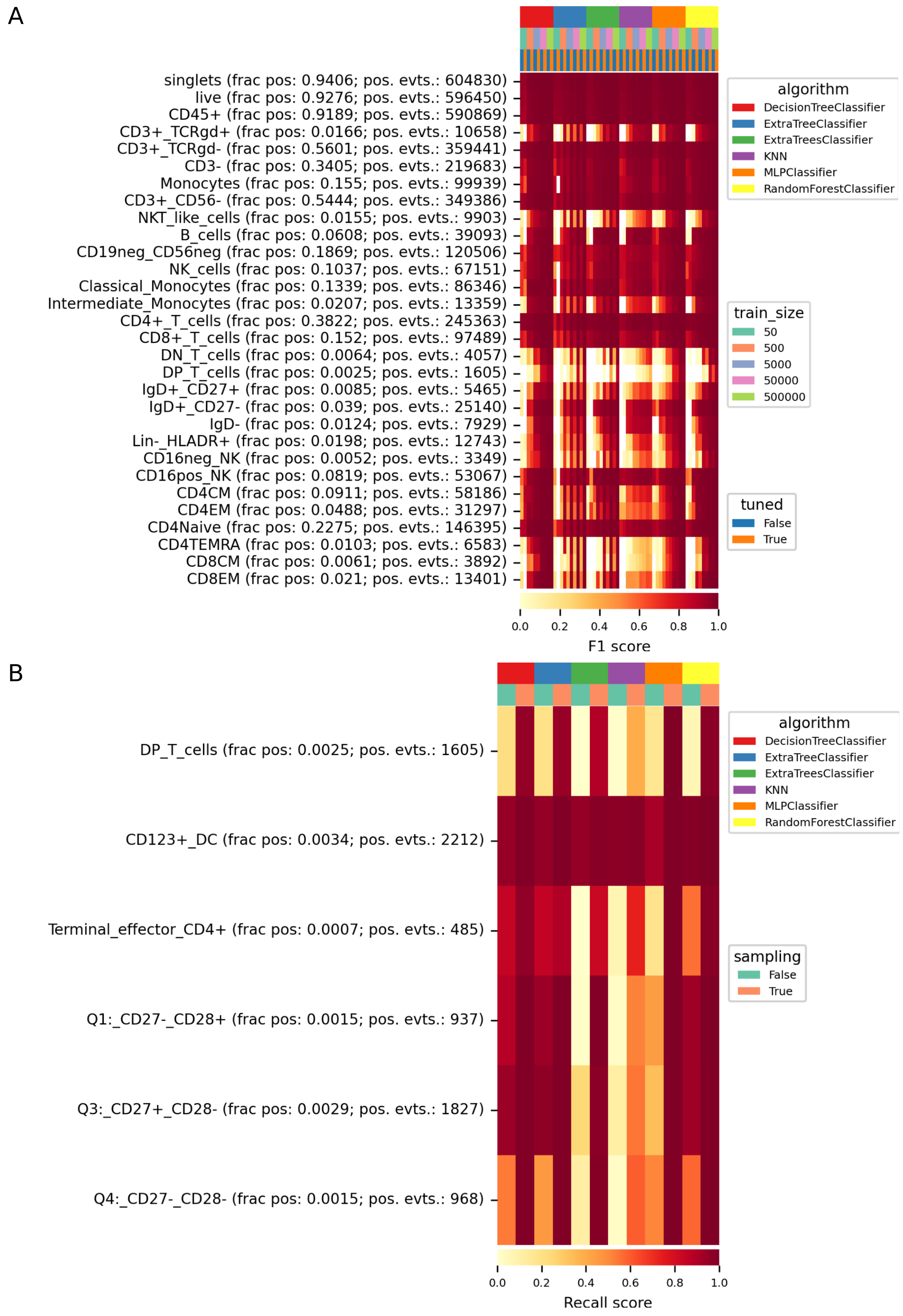


**Supplementary Figure S5: Classifier characterization on a human flow cytometry dataset.** The dataset consisted of 4 samples of human peripheral blood cells (flow cytometry; Dataset 6). **A** Classifiers were compared using the indicated train sizes and hyperparameter tuning (top column annotations). Shown are the mean F1 metrics per indicated gate. For each gate, the fraction of positive events (frac pos) as well as the total number of positive events (pos. evts.) are indicated. **B** Recall scores for gates below 0.005% after using a dedicated sampling strategy (sampling: True) compared to random subsampled data. MLP: multi-layer-perceptron, KNN: k-nearest-neighbor classifier

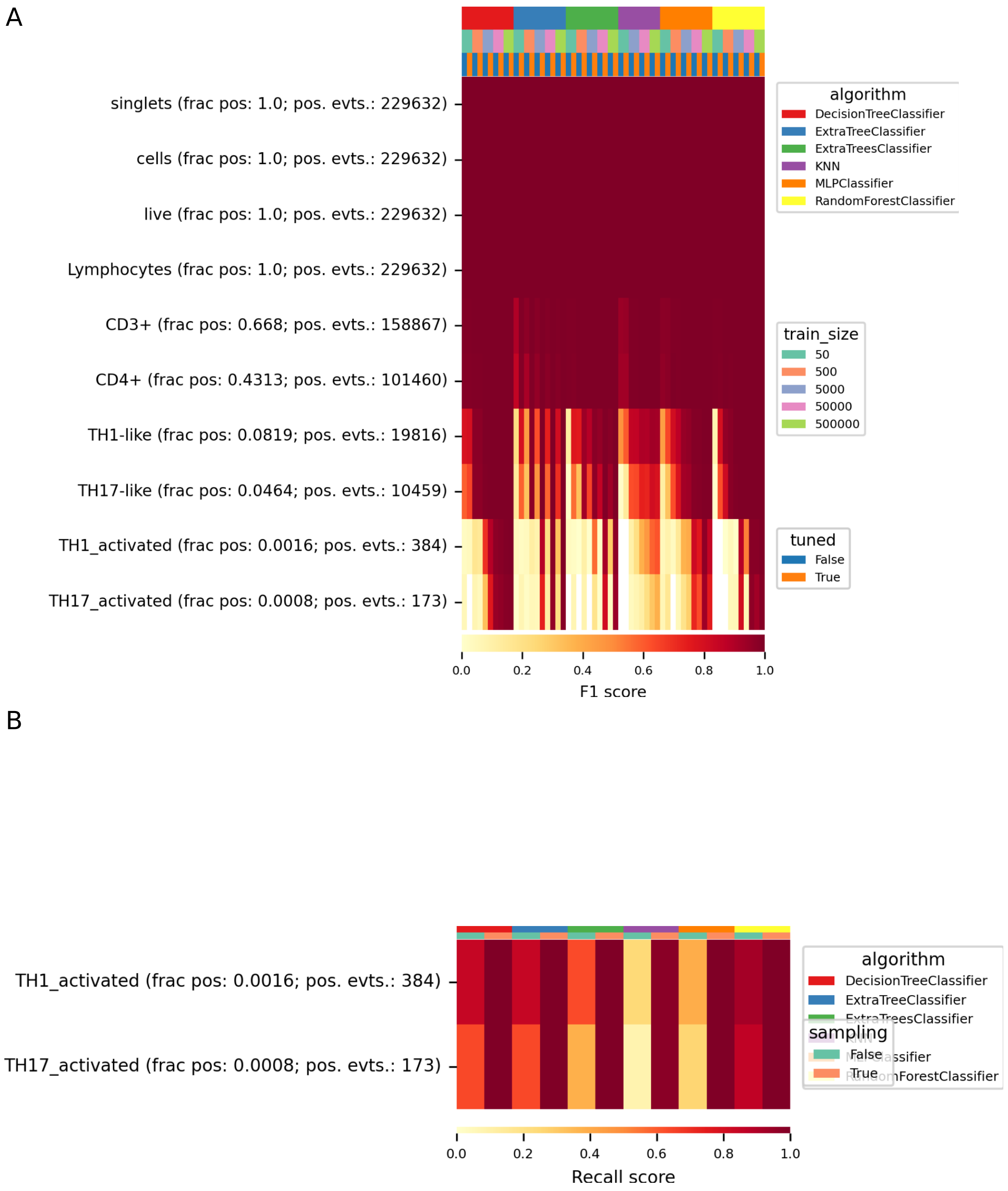


**Supplementary Figure S6: Classifier characterization on a human flow cytometry dataset.** The dataset consisted of 107 samples of human peripheral blood cells (flow cytometry; Dataset 5). **A** Classifiers were compared using the indicated train sizes and hyperparameter tuning (top column annotations). Shown are the mean F1 metrics per indicated gate. For each gate, the fraction of positive events (frac pos) as well as the total number of positive events (pos. evts.) are indicated. **B** Recall scores for gates below 0.005% after using a dedicated sampling strategy (sampling: True) compared to random subsampled data. MLP: multi-layer-perceptron, KNN: k-nearest-neighbor classifier

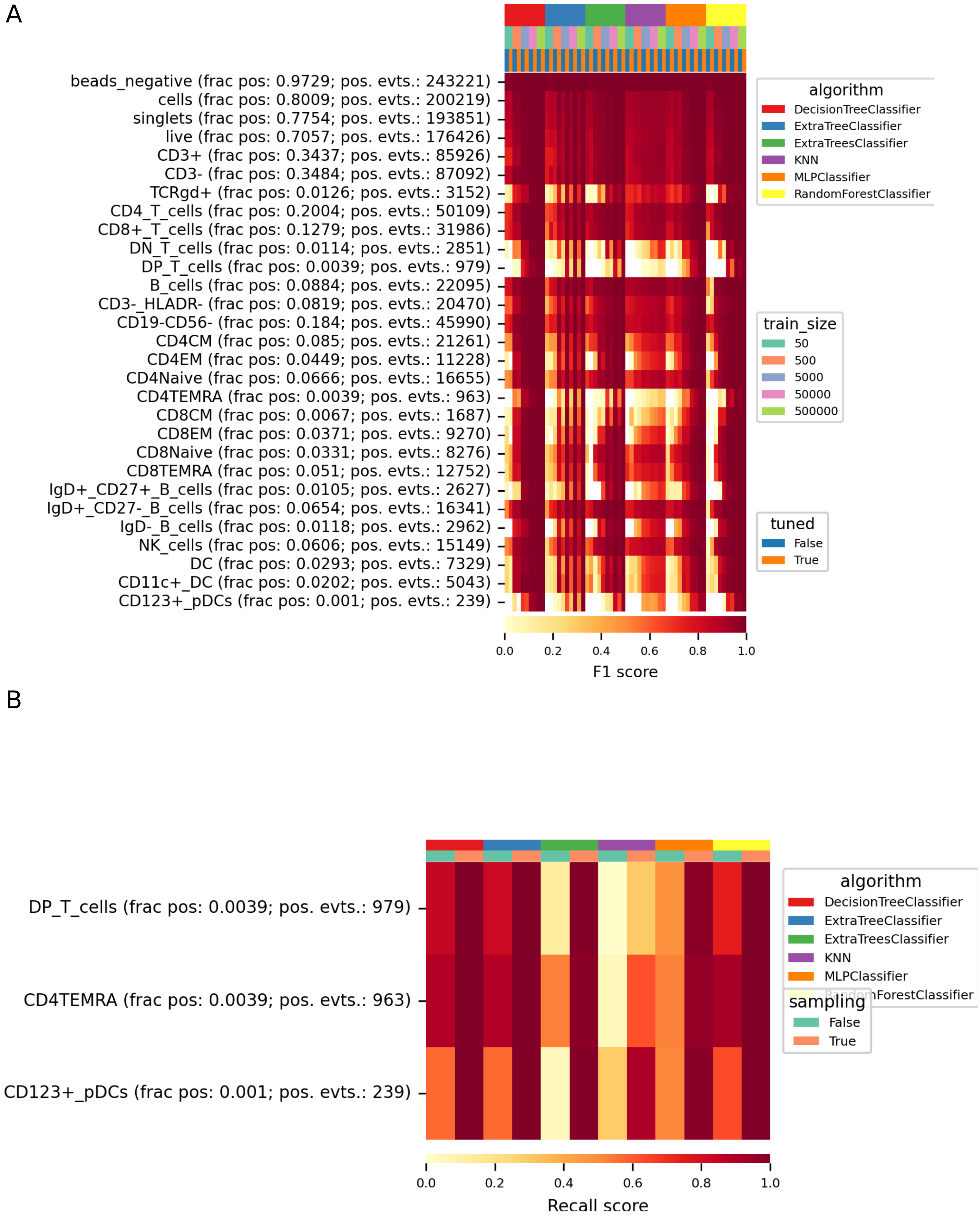


**Supplementary Figure S7: Classifier characterization on a human mass cytometry dataset.** The dataset consisted of 19 samples of human peripheral blood cells (mass cytometry; Dataset 7). **A** Classifiers were compared using the indicated train sizes and hyperparameter tuning (top column annotations). Shown are the mean F1 metrics per indicated gate. For each gate, the fraction of positive events (frac pos) as well as the total number of positive events (pos. evts.) are indicated. **B** Recall scores for gates below 0.005% after using a dedicated sampling strategy (sampling: True) compared to random subsampled data. MLP: multi-layer-perceptron, KNN: k-nearest-neighbor classifier

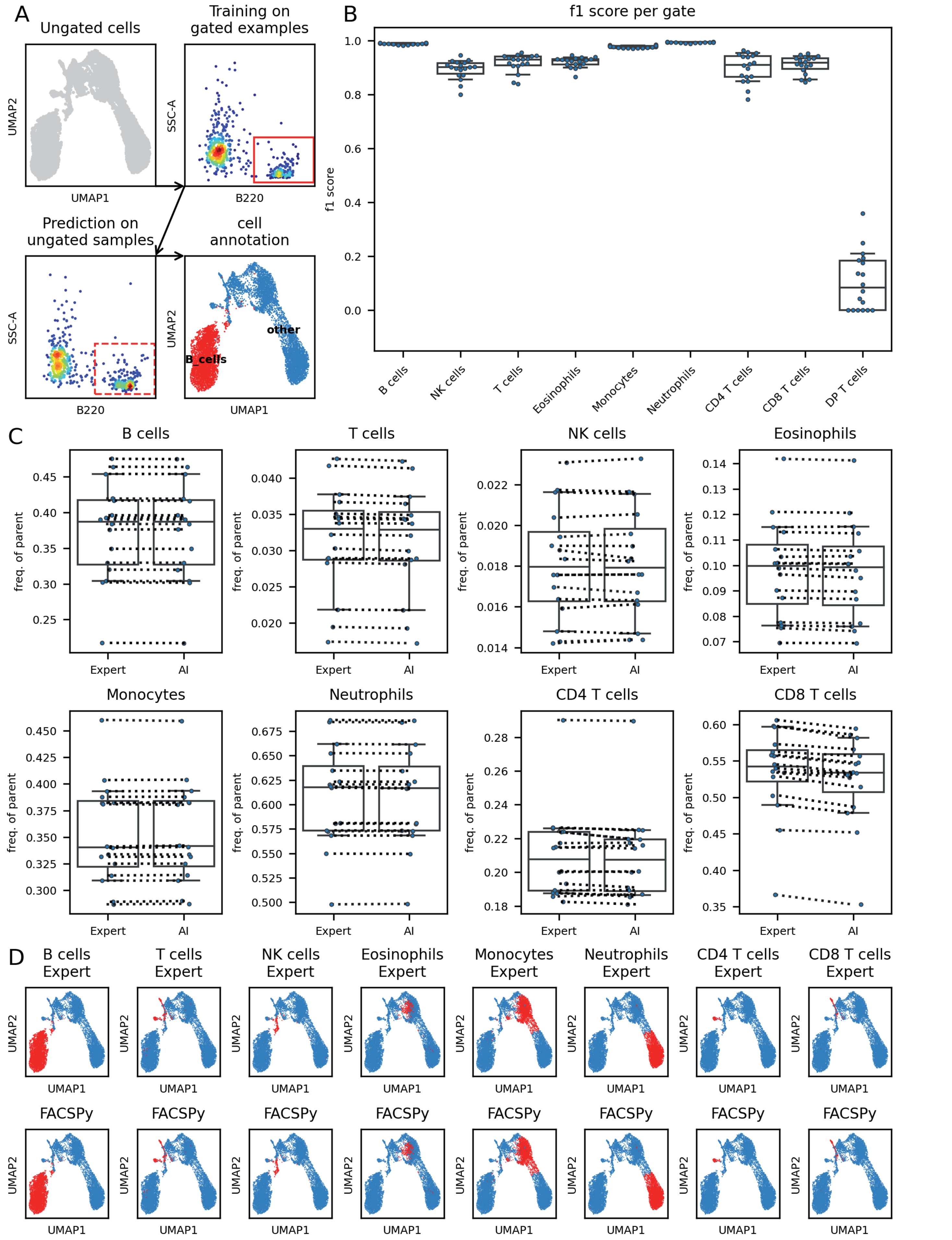


**Supplementary Figure S8: Classifier characterization on a murine flow cytometry dataset.** The dataset consisted of 18 samples of mouse bone marrow cells (flow cytometry; Dataset 1). **A** Graphical abstract. Example samples are used to train a classifier using the detected marker expression and the gates. The classifier is then used to classify previously ungated cells. **B** A classifier using tuned hyperparameters (compare **Supplementary Figure S1**) was used as in B. F1 scores are plotted on the y-axis respective to the individual gates (x-axis). **C** High concordance of the frequency of parents of a manually drawn expert gate (left) and the AI-gated cells (right). Dotted lines connect corresponding samples. **D** High concordance of cell classification on a single cell level. Shown is a UMAP embedding colored for the indicated cell type. **Top**: cells classified by manual expert gating. **Bottom**: cells classified by supervised classification. MLP: multi-layer perceptron, KNN: k-nearest-neighbor classifier

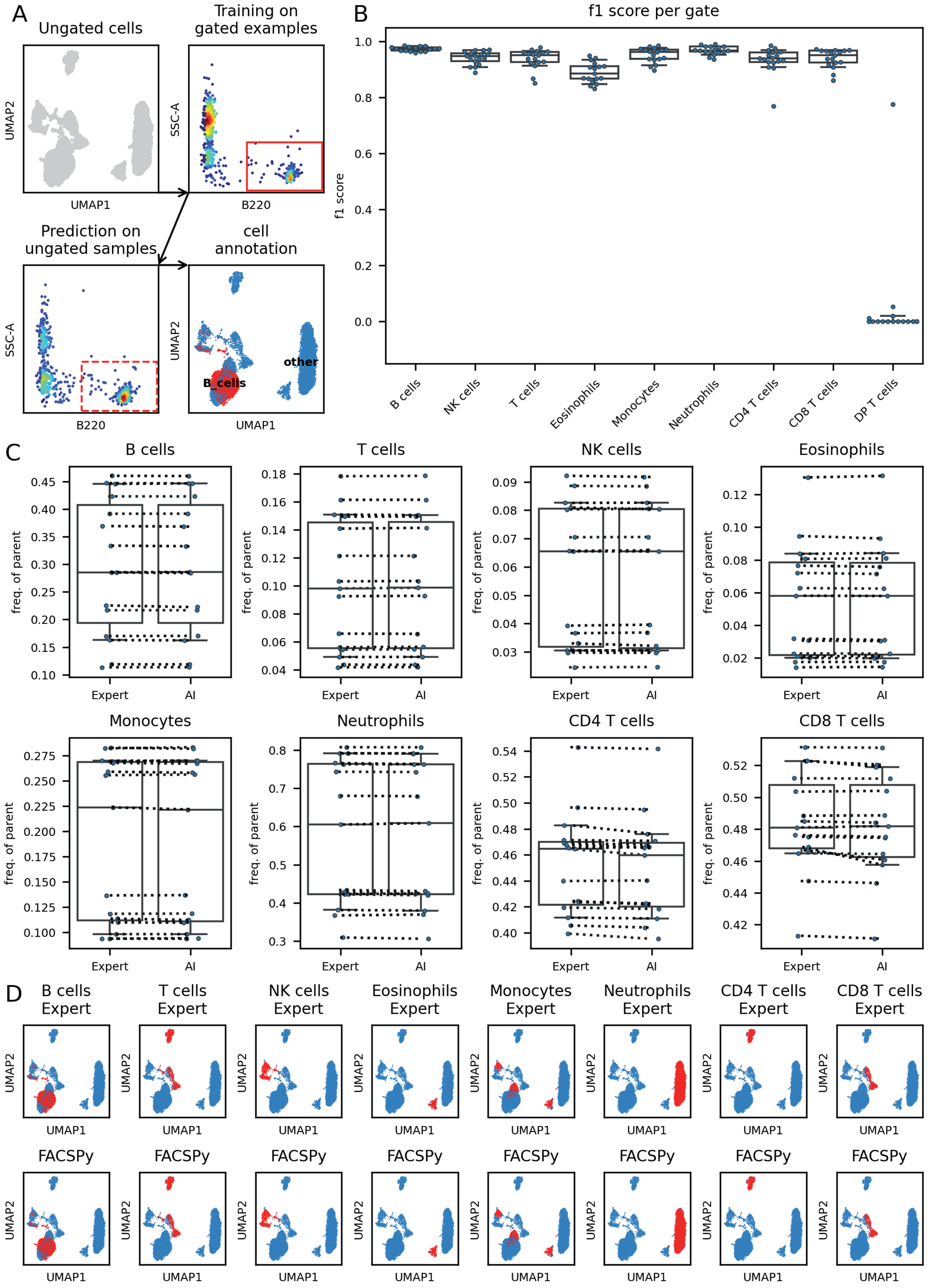


**Supplementary Figure S9: Classifier characterization on a murine flow cytometry dataset.** The dataset consisted of 18 samples of mouse peripheral blood cells (flow cytometry; Dataset 2). **A** Graphical abstract. Example samples are used to train a classifier using the detected marker expression and the gates. The classifier is then used to classify previously ungated cells. **B** A classifier using tuned hyperparameters (compare **Supplementary Figure S2**) was used as in B. F1 scores are plotted on the y-axis respective to the individual gates (x-axis). **C** High concordance of the frequency of parents of a manually drawn expert gate (left) and the AI-gated cells (right). Dotted lines connect corresponding samples. **D** High concordance of cell classification on a single cell level. Shown is a UMAP embedding colored for the indicated cell type. **Top**: cells classified by manual expert gating. **Bottom**: cells classified by supervised classification. MLP: multi-layer perceptron, KNN: k-nearest-neighbor classifier

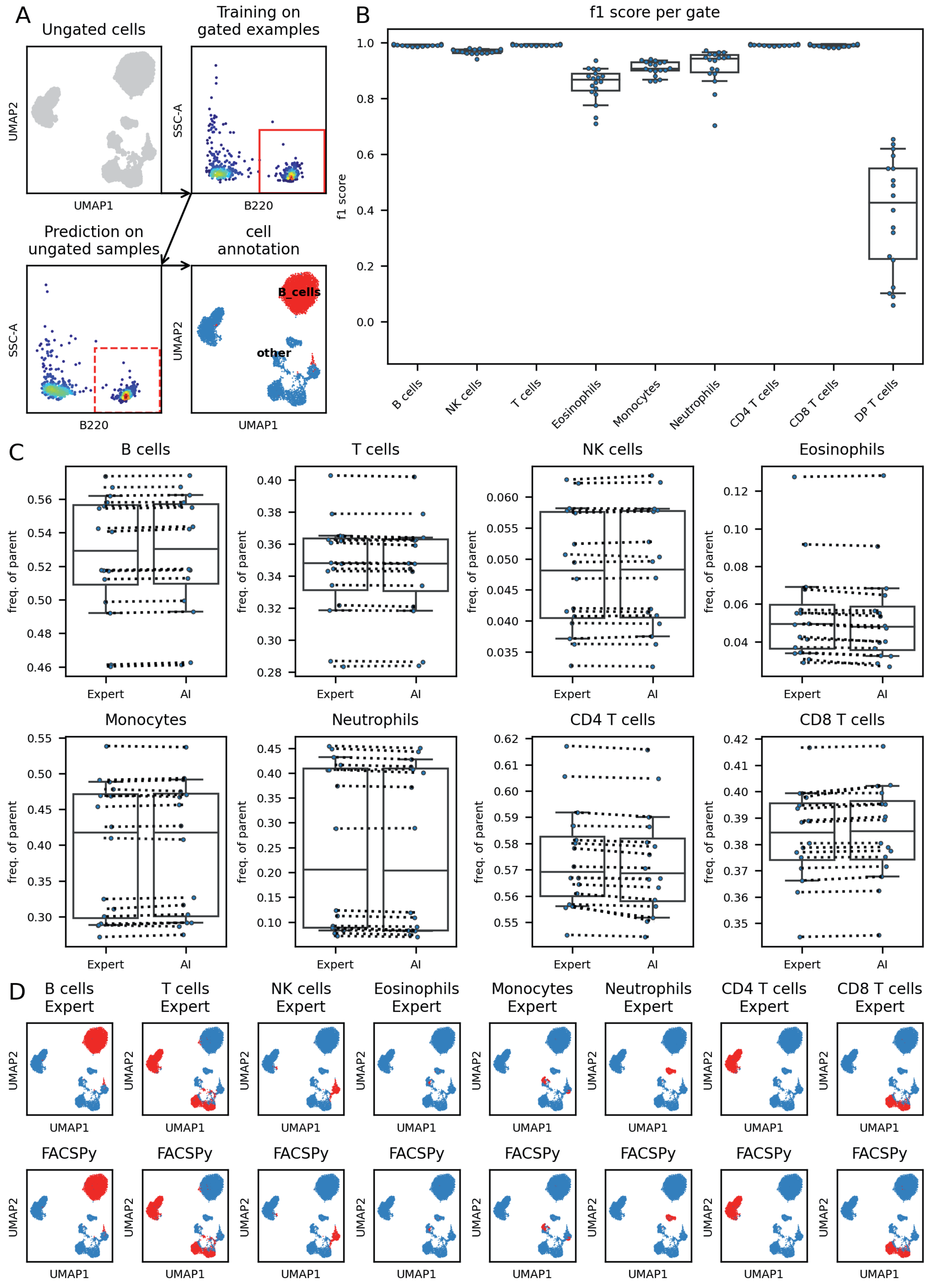


**Supplementary Figure S10: Classifier characterization on a murine flow cytometry dataset.** The dataset consisted of 18 samples of mouse spleen cells (flow cytometry; Dataset 3). **A** Graphical abstract. Example samples are used to train a classifier using the detected marker expression and the gates. The classifier is then used to classify previously ungated cells. **B** A classifier using tuned hyperparameters (compare **Supplementary Figure S3**) was used as in B. F1 scores are plotted on the y-axis respective to the individual gates (x-axis). **C** High concordance of the frequency of parents of a manually drawn expert gate (left) and the AI-gated cells (right). Dotted lines connect corresponding samples. **D** High concordance of cell classification on a single cell level. Shown is a UMAP embedding colored for the indicated cell type. **Top**: cells classified by manual expert gating. **Bottom**: cells classified by supervised classification. MLP: multi-layer perceptron, KNN: k-nearest-neighbor classifier

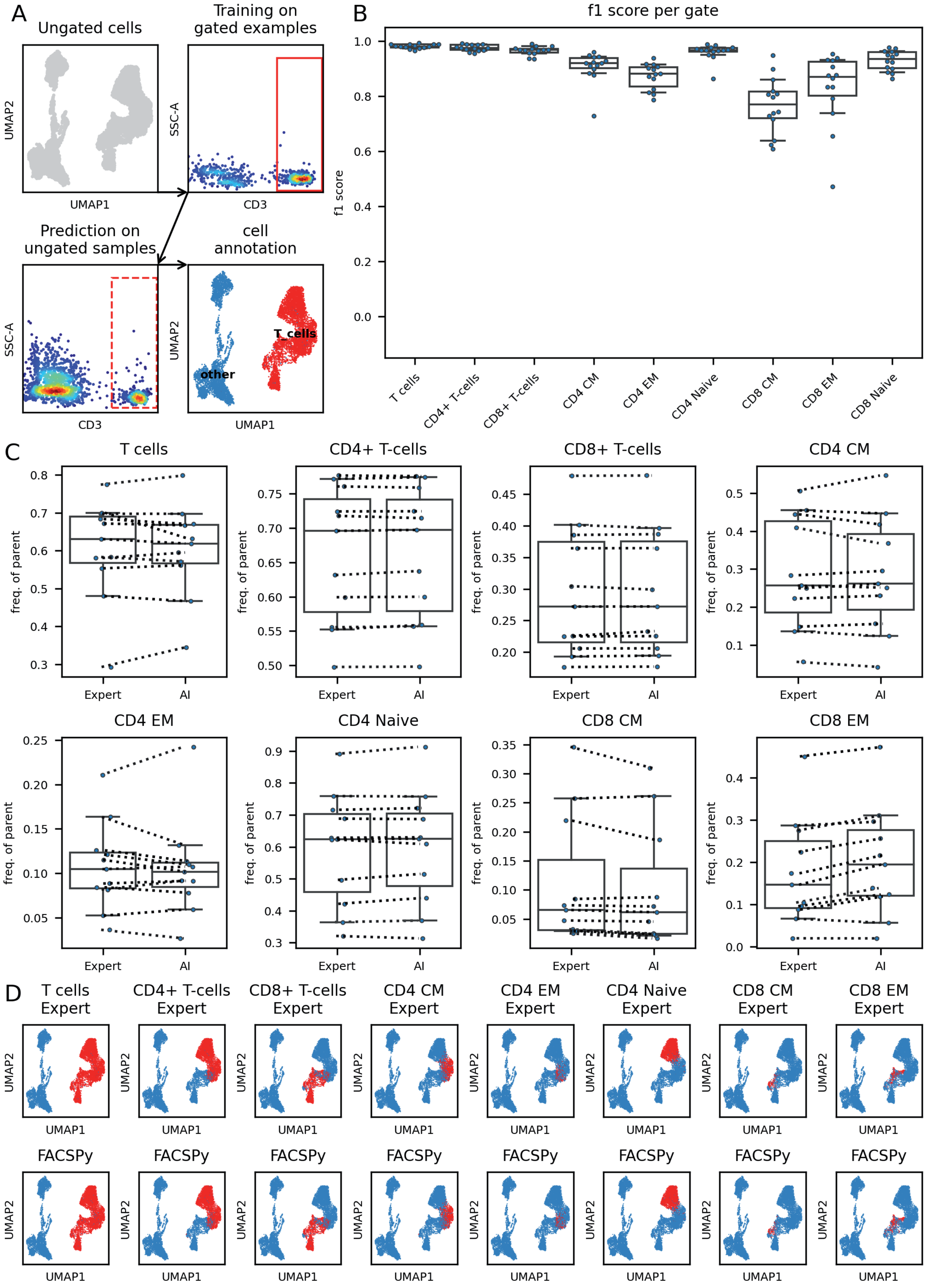


**Supplementary Figure S11: Classifier characterization on a human flow cytometry dataset.** The dataset consisted of 14 samples of human peripheral blood cells (flow cytometry; Dataset 4). **A** Graphical abstract. Example samples are used to train a classifier using the detected marker expression and the gates. The classifier is then used to classify previously ungated cells. **B** A classifier using tuned hyperparameters (compare **Supplementary Figure S4**) was used as in B. F1 scores are plotted on the y-axis respective to the individual gates (x-axis). **C** High concordance of the frequency of parents of a manually drawn expert gate (left) and the AI-gated cells (right). Dotted lines connect corresponding samples. **D** High concordance of cell classification on a single cell level. Shown is a UMAP embedding colored for the indicated cell type. **Top**: cells classified by manual expert gating. **Bottom**: cells classified by supervised classification. MLP: multi-layer perceptron, KNN: k-nearest-neighbor classifier

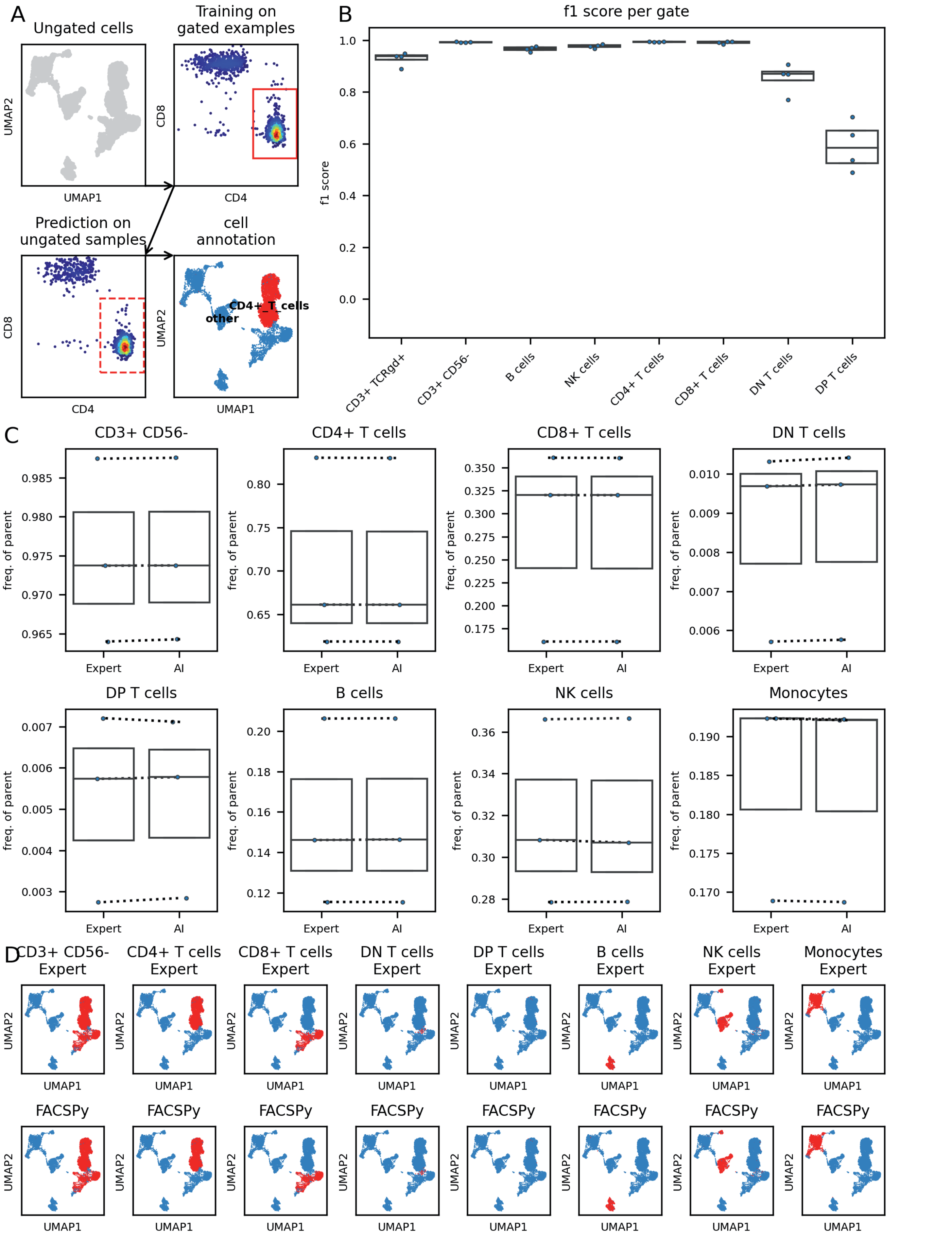


**Supplementary Figure S12: Classifier characterization on a human flow cytometry dataset.** The dataset consisted of 4 samples of human peripheral blood cells (spectral flow cytometry; Dataset 6). **A** Graphical abstract. Example samples are used to train a classifier using the detected marker expression and the gates. The classifier is then used to classify previously ungated cells. **B** A classifier using tuned hyperparameters (compare **Supplementary Figure S5**) was used as in B. F1 scores are plotted on the y-axis respective to the individual gates (x-axis). **C** High concordance of the frequency of parents of a manually drawn expert gate (left) and the AI-gated cells (right). Dotted lines connect corresponding samples. **D** High concordance of cell classification on a single cell level. Shown is a UMAP embedding colored for the indicated cell type. **Top**: cells classified by manual expert gating. **Bottom**: cells classified by supervised classification. MLP: multi-layer perceptron, KNN: k-nearest-neighbor classifier

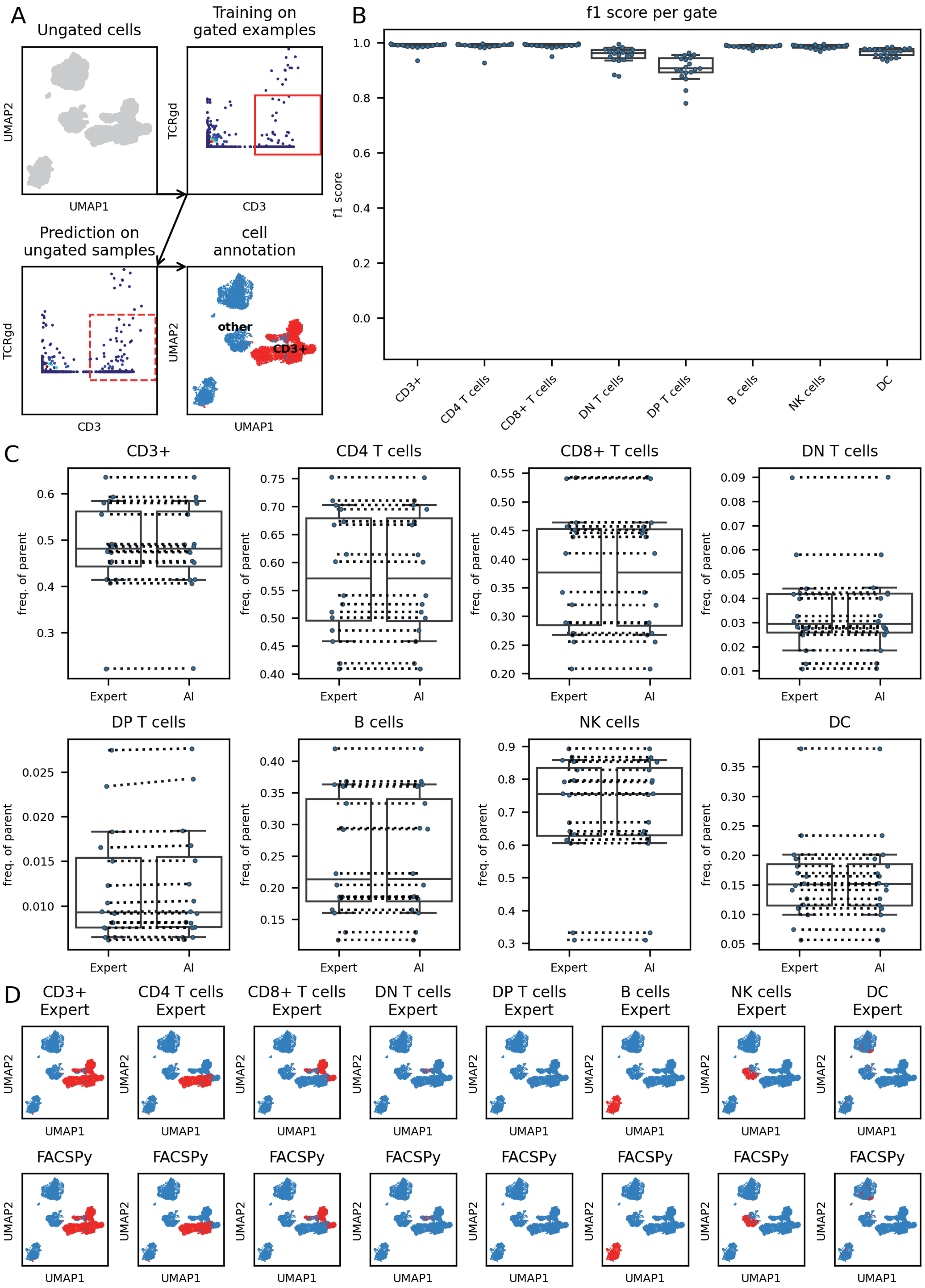


**Supplementary Figure S13: Classifier characterization on a human mass cytometry dataset.** The dataset consisted of 19 samples of human peripheral blood cells (mass cytometry; Dataset 7). **A** Graphical abstract. Example samples are used to train a classifier using the detected marker expression and the gates. The classifier is then used to classify previously ungated cells. **B** A classifier using tuned hyperparameters (compare **Supplementary Figure S7**) was used as in B. F1 scores are plotted on the y-axis respective to the individual gates (x-axis). **C** High concordance of the frequency of parents of a manually drawn expert gate (left) and the AI-gated cells (right). Dotted lines connect corresponding samples. **D** High concordance of cell classification on a single cell level. Shown is a UMAP embedding colored for the indicated cell type. **Top**: cells classified by manual expert gating. **Bottom**: cells classified by supervised classification. MLP: multi-layer perceptron, KNN: k-nearest-neighbor classifier

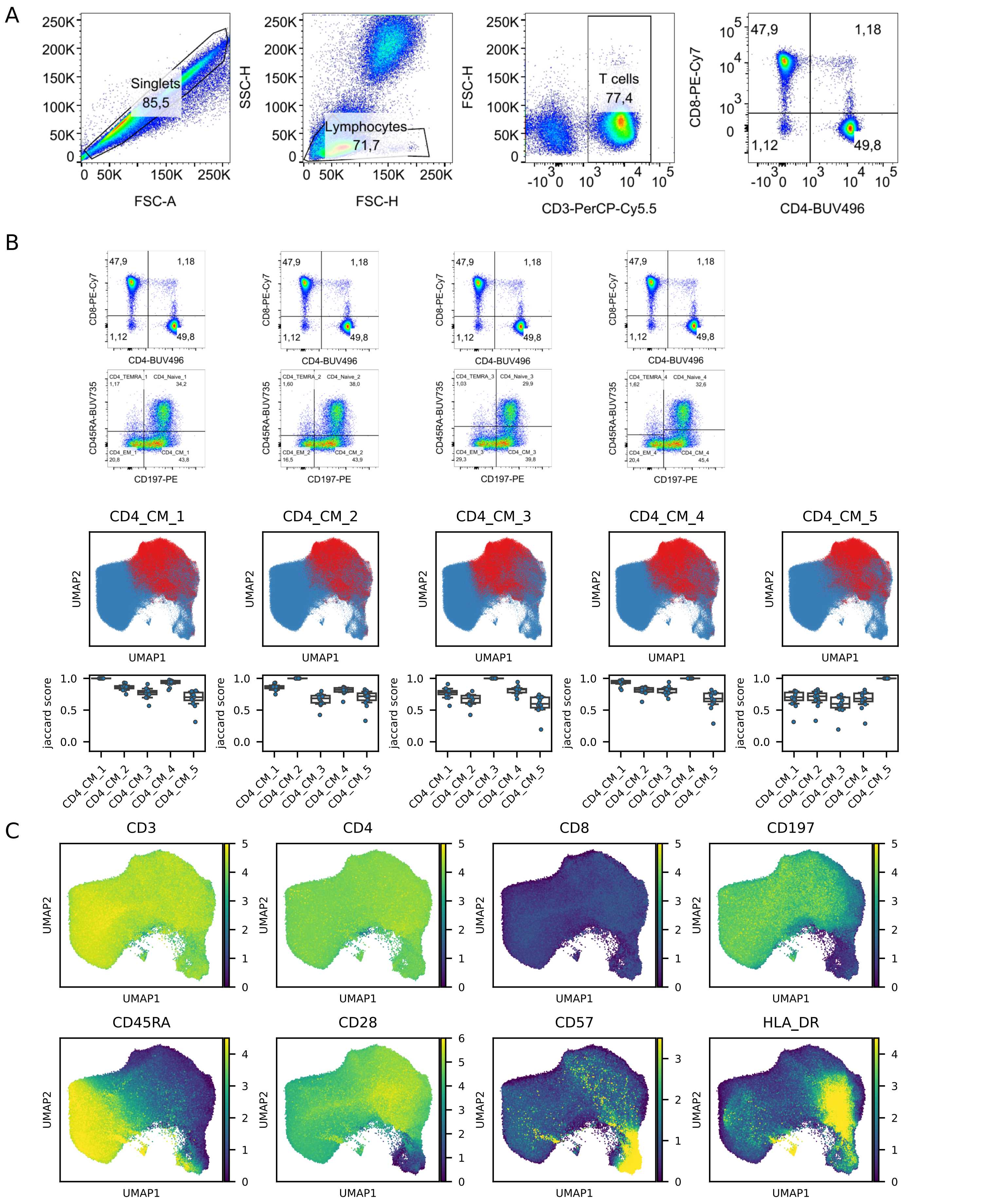


**Supplementary Figure S14: Discrepancy of cell type identification in continuous marker expressions. A** Gating Strategy. **B Top:** Four different gating strategies were applied to each sample manually. The first gating strategy aimed to split the populations by drawing the gates at the lowest point density between the points. The second gating strategy was aimed to represent a sensitive gating strategy for positive events while the third gating strategy was very specific for positive events. The fourth gating strategy used different positive-cutoffs in the CD45RA channel for the distinction of Naïve CD4+ T-cells and CD4-TEMRA. CD4_CM_5 denotes cells identified by the semi-supervised gating. **Mid**: UMAP representation of CD4+ T-cells colored by the identified CD4+ central memory T-cells identified by the respective approach. **Bottom**: Jaccard indices quantifying the overlap between the respective gating strategies. **C** UMAP representation of CD4+ T-cells colored by the indicated marker proteins.

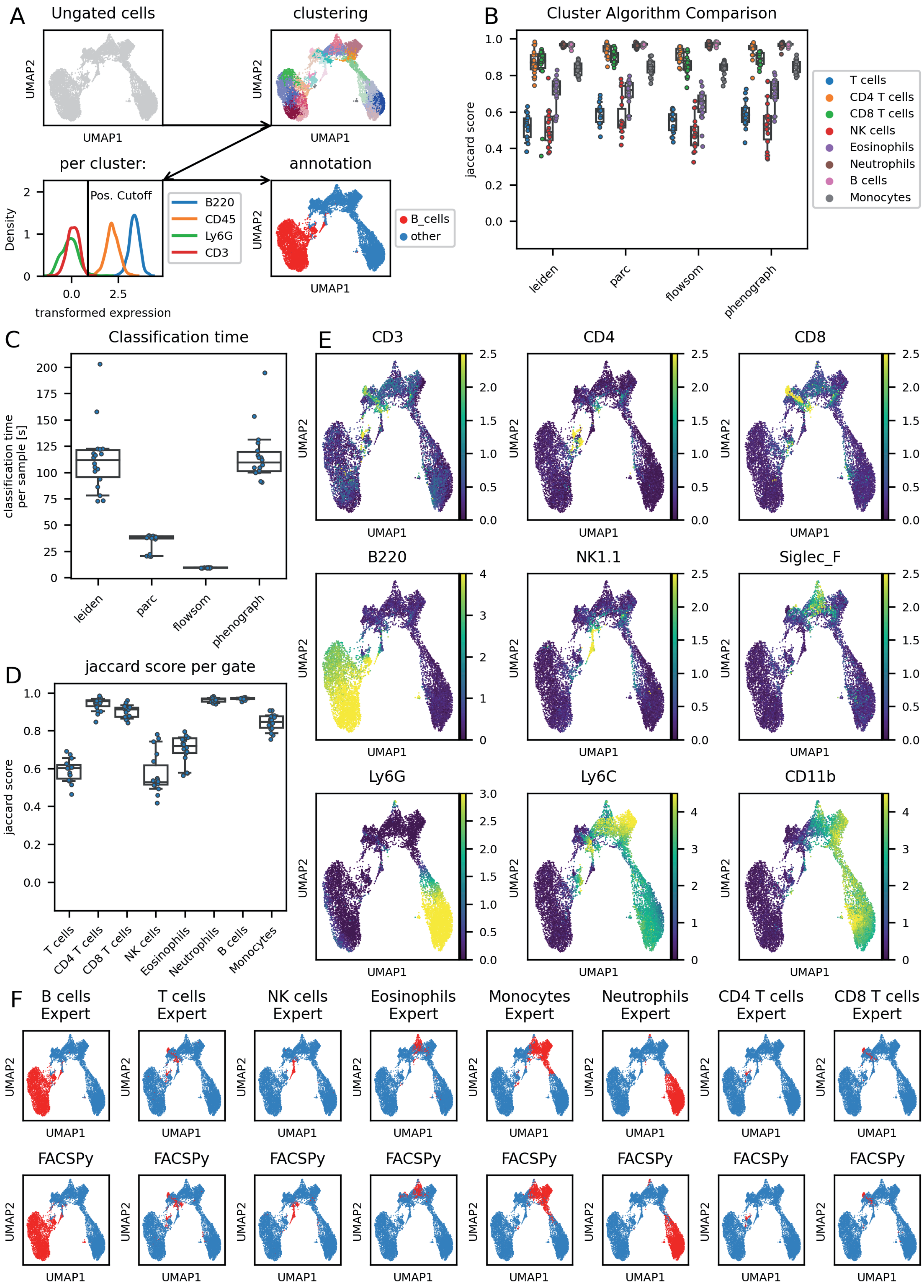


**Supplementary Figure S15: Semi-supervised gating.** The dataset consisted of 18 samples of mouse bone marrow cells (flow cytometry; Dataset 1). **A** Graphical abstract: ungated cells are clustered and each cluster is inspected for its median marker expression. Marker intensity is divided into positive, low, intermediate or high expression and cells are classified based on the user defined cell characterization. **B** Algorithm comparison. The indicated clustering algorithms were used and the marker expressions per cluster were used to gate cells as described in A. **C** Algorithm benchmark. Clustering time was compared for the indicated algorithms. The PARC algorithm showed comparable calculation times compared to FlowSOM, while maintaining high accuracy. **D** PARC algorithm was used with tuned hyperparameters (compare **Supplementary Figure S21**), and accuracy scores are plotted per gate. Each dot represents a single sample analyzed. **E** Marker expression on UMAP embedding. **F** High concordance of cell classification on a single cell level. UMAP embeddings are colored for the indicated cell type (red) for either manual expert gating (top row) or semi-supervised determined cell types (bottom row).

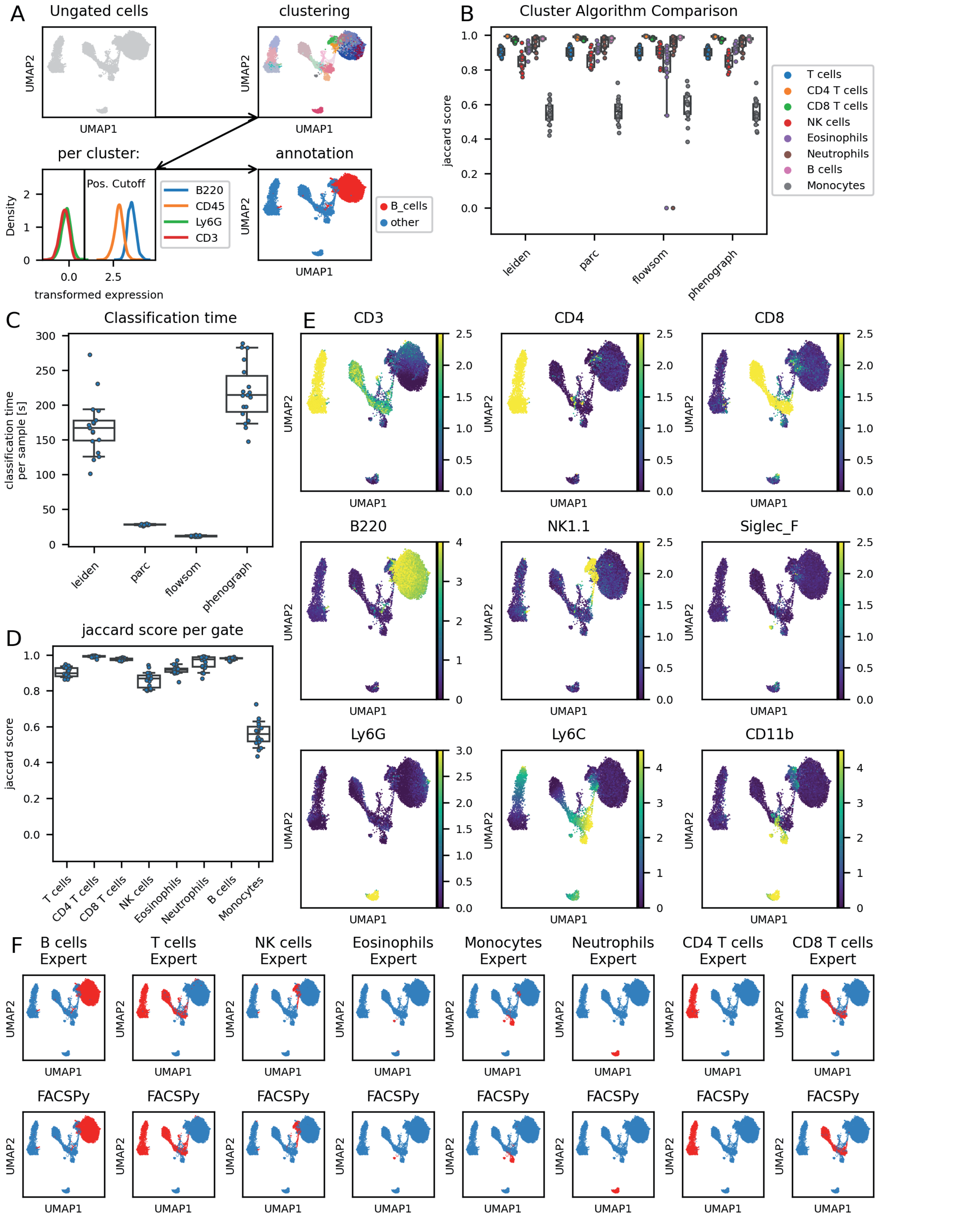


**Supplementary Figure S16: Semi-supervised gating.** The dataset consisted of 18 samples of mouse spleen cells (flow cytometry; Dataset 3). **A** Graphical abstract: ungated cells are clustered and each cluster is inspected for its median marker expression. Marker intensity is divided into positive, low, intermediate or high expression and cells are classified based on the user defined cell characterization. **B** Algorithm comparison. The indicated clustering algorithms were used and the marker expressions per cluster were used to gate cells as described in A. **C** Algorithm benchmark. Clustering time was compared for the indicated algorithms. The PARC algorithm showed comparable calculation times compared to FlowSOM, while maintaining high accuracy. **D** PARC algorithm was used with tuned hyperparameters (compare **Supplementary Figure S23**), and accuracy scores are plotted per gate. Each dot represents a single sample analyzed. **E** Marker expression on UMAP embedding. **F** High concordance of cell classification on a single cell level. UMAP embeddings are colored for the indicated cell type (red) for either manual expert gating (top row) or semi-supervised determined cell types (bottom row).

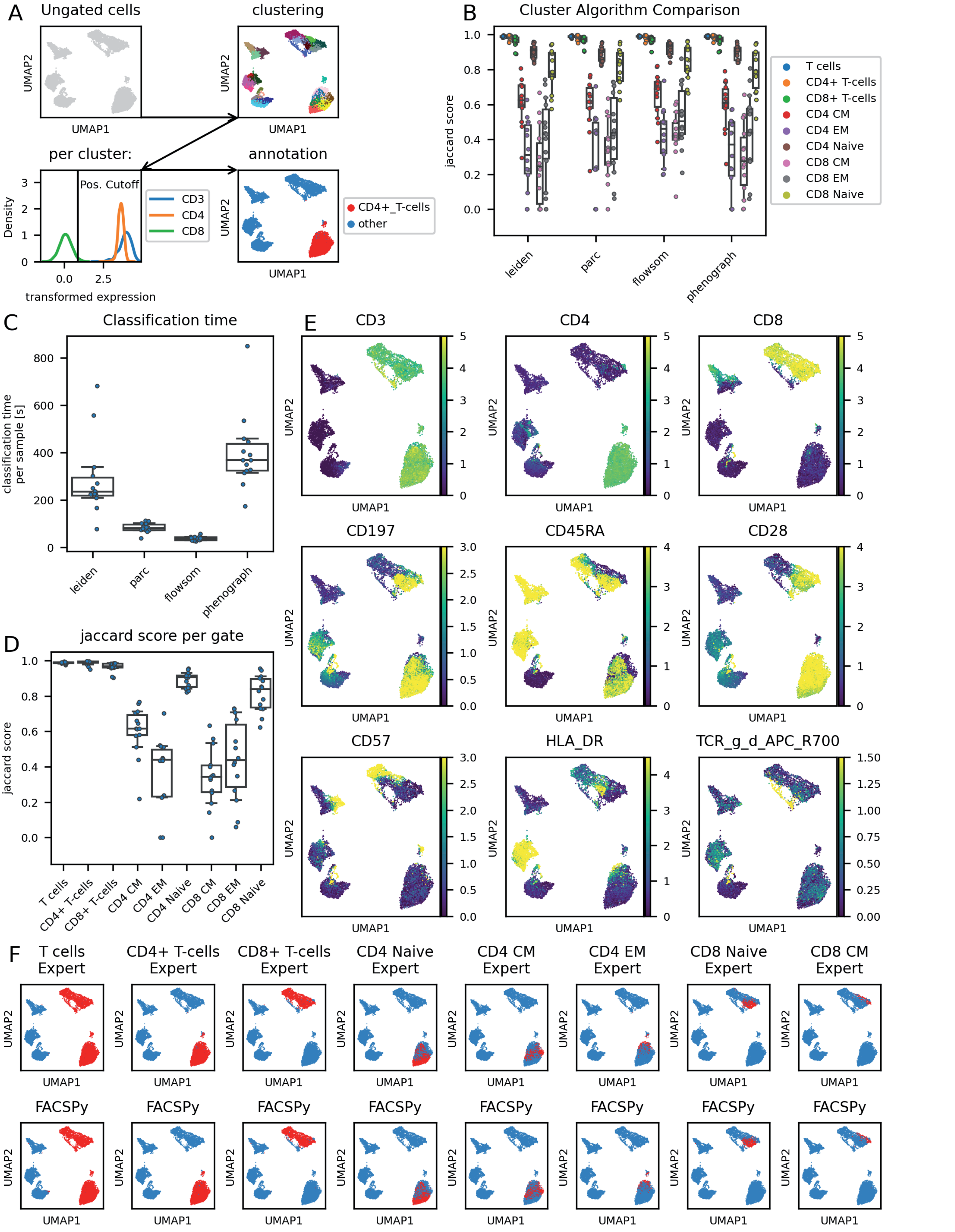


**Supplementary Figure S17: Semi-supervised gating.** The dataset consisted of 14 samples of human peripheral blood cells (flow cytometry; Dataset 4). **A** Graphical abstract: ungated cells are clustered and each cluster is inspected for its median marker expression. Marker intensity is divided into positive, low, intermediate or high expression and cells are classified based on the user defined cell characterization. **B** Algorithm comparison. The indicated clustering algorithms were used and the marker expressions per cluster were used to gate cells as described in A. **C** Algorithm benchmark. Clustering time was compared for the indicated algorithms. The PARC algorithm showed comparable calculation times compared to FlowSOM, while maintaining high accuracy. **D** PARC algorithm was used with tuned hyperparameters (compare **Supplementary Figure S24**), and accuracy scores are plotted per gate. Each dot represents a single sample analyzed. **E** Marker expression on UMAP embedding. **F** High concordance of cell classification on a single cell level. UMAP embeddings are colored for the indicated cell type (red) for either manual expert gating (top row) or semi-supervised determined cell types (bottom row).

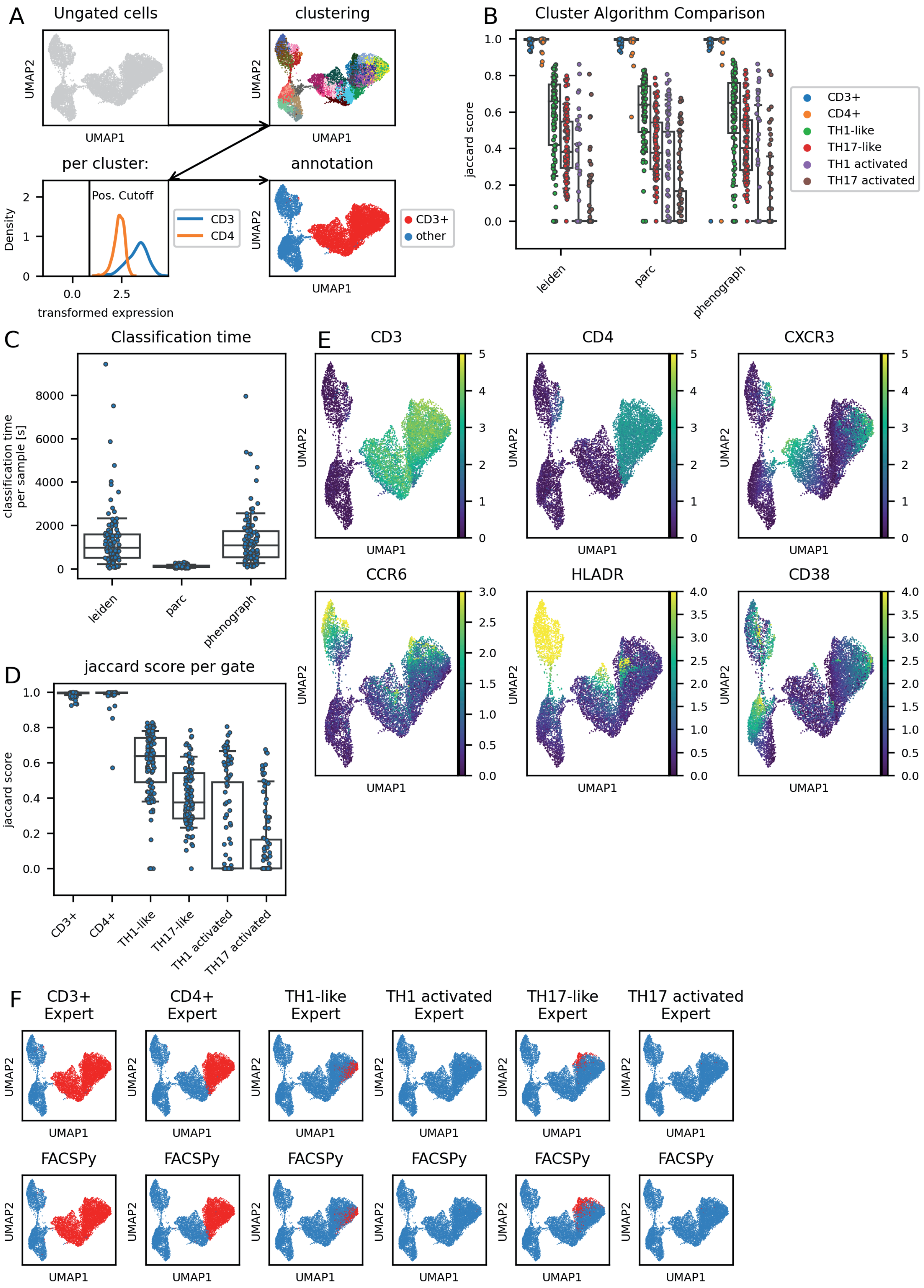


**Supplementary Figure S19: Semi-supervised gating.** The dataset consisted of 4 samples of human peripheral blood cells (flow cytometry; Dataset 6). **A** Graphical abstract: ungated cells are clustered and each cluster is inspected for its median marker expression. Marker intensity is divided into positive, low, intermediate or high expression and cells are classified based on the user defined cell characterization. **B** Algorithm comparison. The indicated clustering algorithms were used and the marker expressions per cluster were used to gate cells as described in A. **C** Algorithm benchmark. Clustering time was compared for the indicated algorithms. The PARC algorithm showed comparable calculation times compared to FlowSOM, while maintaining high accuracy. **D** PARC algorithm was used with tuned hyperparameters (compare **Supplementary Figure S27**), and accuracy scores are plotted per gate. Each dot represents a single sample analyzed. **E** Marker expression on UMAP embedding. **F** High concordance of cell classification on a single cell level. UMAP embeddings are colored for the indicated cell type (red) for either manual expert gating (top row) or semi-supervised determined cell types (bottom row).

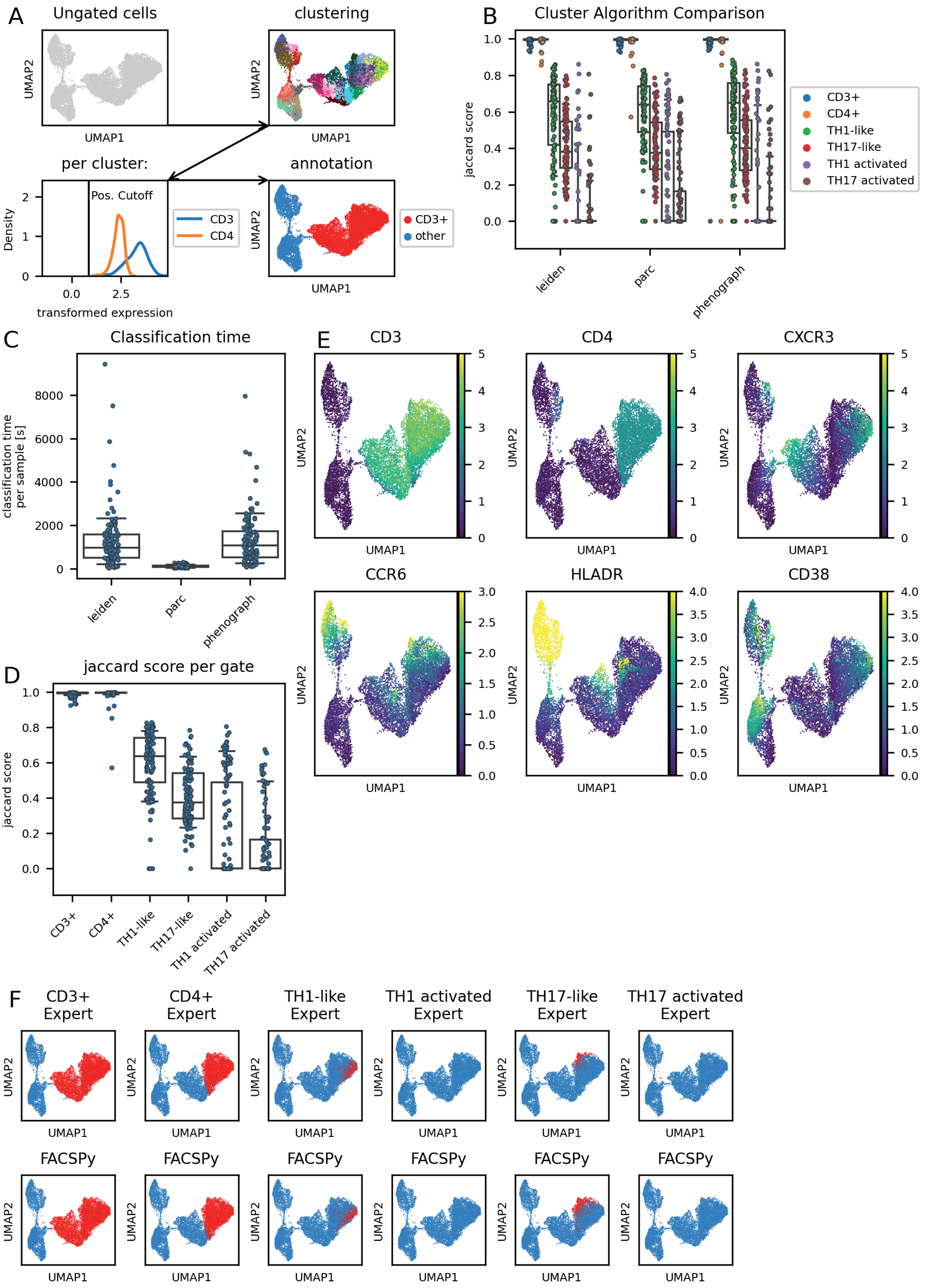


**Supplementary Figure S19: Semi-supervised gating.** The dataset consisted of 107 samples of human peripheral blood cells (flow cytometry; Dataset 5). **A** Graphical abstract: ungated cells are clustered and each cluster is inspected for its median marker expression. Marker intensity is divided into positive, low, intermediate or high expression and cells are classified based on the user defined cell characterization. **B** Algorithm comparison. The indicated clustering algorithms were used and the marker expressions per cluster were used to gate cells as described in A. **C** Algorithm benchmark. Clustering time was compared for the indicated algorithms. The PARC algorithm showed comparable calculation times compared to FlowSOM, while maintaining high accuracy. **D** PARC algorithm was used with tuned hyperparameters (compare **Supplementary Figure S26**), and accuracy scores are plotted per gate. Each dot represents a single sample analyzed. **E** Marker expression on UMAP embedding. **F** High concordance of cell classification on a single cell level. UMAP embeddings are colored for the indicated cell type (red) for either manual expert gating (top row) or semi-supervised determined cell types (bottom row).

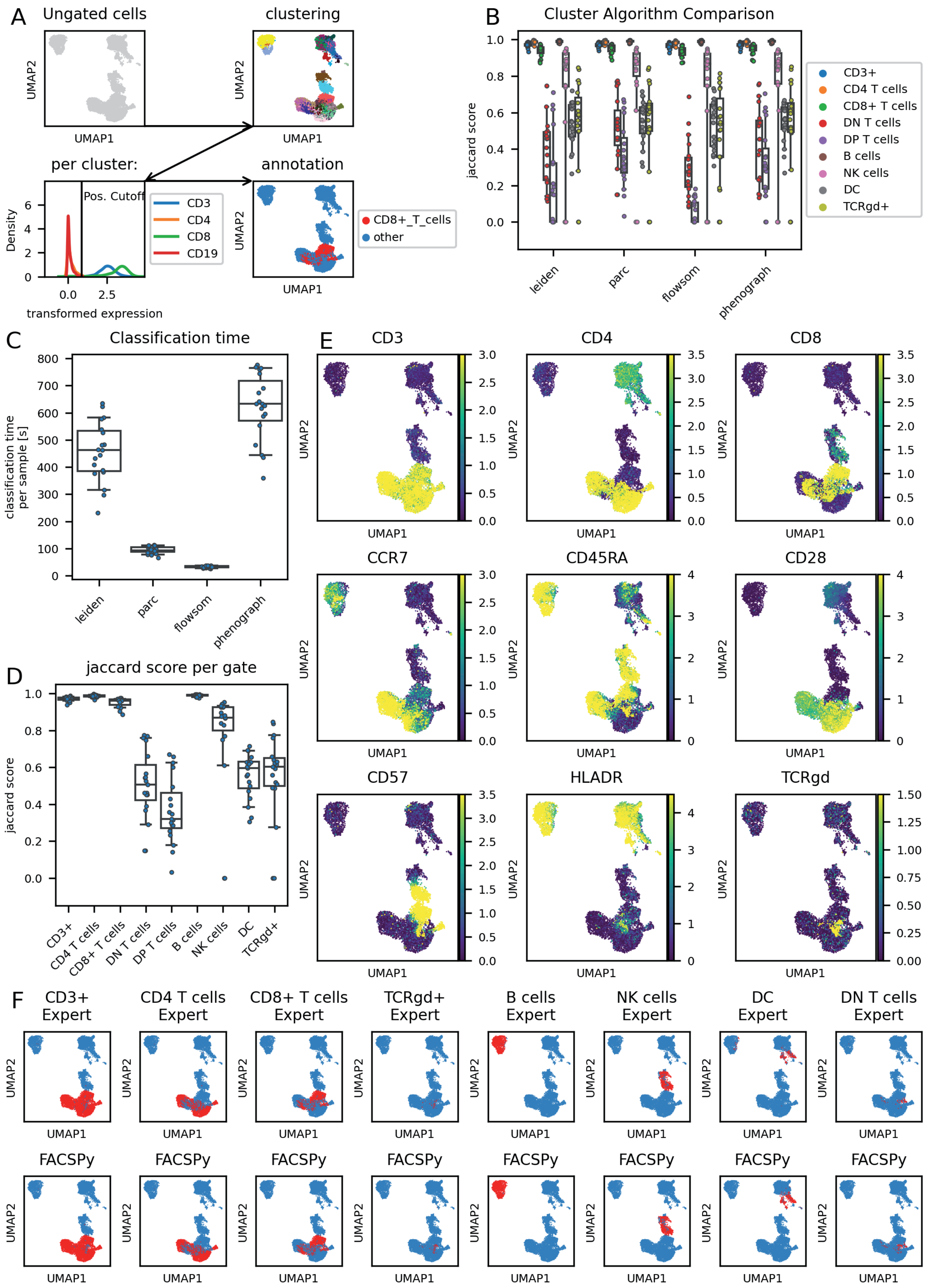


**Supplementary Figure S20: Semi-supervised gating.** The dataset consisted of 19 samples of human peripheral blood cells (mass cytometry; Dataset 7). **A** Graphical abstract: ungated cells are clustered and each cluster is inspected for its median marker expression. Marker intensity is divided into positive, low, intermediate or high expression and cells are classified based on the user defined cell characterization. **B** Algorithm comparison. The indicated clustering algorithms were used and the marker expressions per cluster were used to gate cells as described in A. **C** Algorithm benchmark. Clustering time was compared for the indicated algorithms. The PARC algorithm showed comparable calculation times compared to FlowSOM, while maintaining high accuracy. **D** PARC algorithm was used with tuned hyperparameters (compare **Supplementary Figure S27**), and accuracy scores are plotted per gate. Each dot represents a single sample analyzed. **E** Marker expression on UMAP embedding. **F** High concordance of cell classification on a single cell level. UMAP embeddings are colored for the indicated cell type (red) for either manual expert gating (top row) or semi-supervised determined cell types (bottom row).

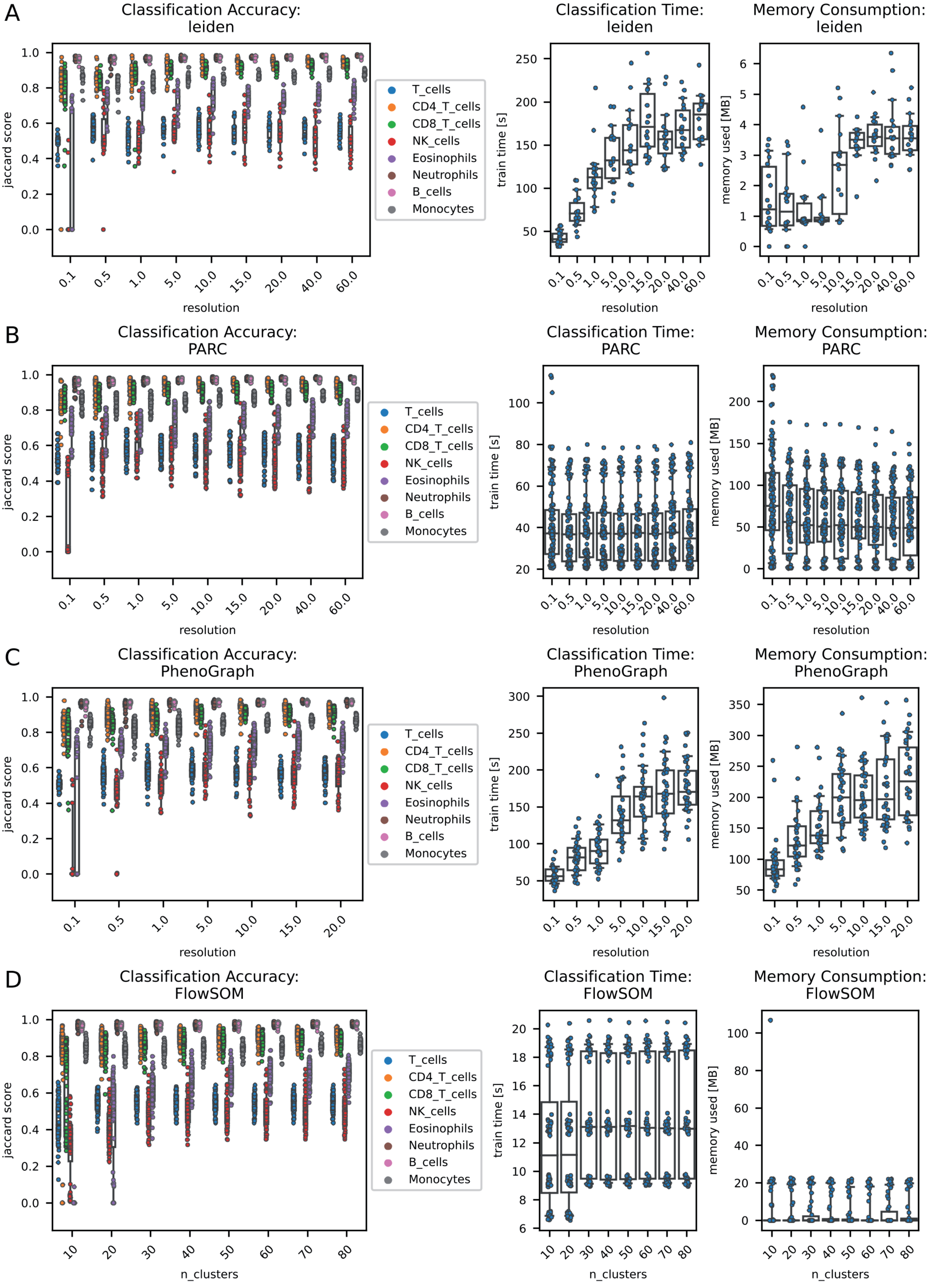


**Supplementary Figure S21: Classifier characterization semi-supervised gating.** The dataset consisted of 18 samples of mouse bone marrow cells (flow cytometry; Dataset 1). Cell line identification was performed as described in the methods section using the clustering algorithms were leiden (**A**), PARC (**B**), phenograph (**C**) and FlowSOM (**D**). In order to test the effect of the resolution parameter, the functions were ran concurrently with a different setting for the cluster resolution (left panel). For FlowSOM, the number of metaclusters was adjusted. The time needed for classification of one sample (mid panel) and the memory consumption during the classification (right panel) was measured.

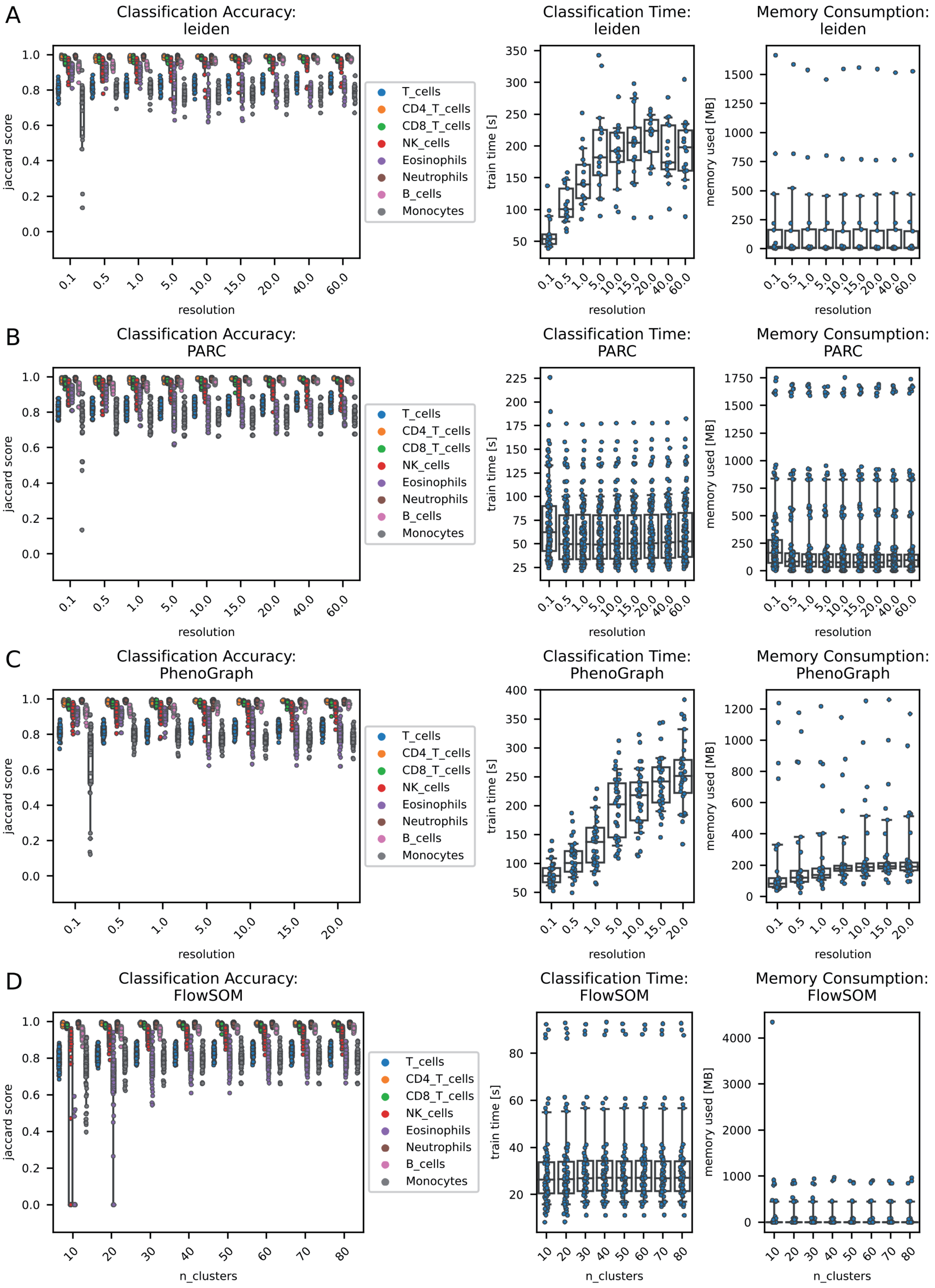


**Supplementary Figure S22: Classifier characterization semi-supervised gating.** The dataset consisted of 18 samples of mouse peripheral blood cells (flow cytometry; Dataset 2). Cell line identification was performed as described in the methods section using the clustering algorithms were leiden (**A**), PARC (**B**), phenograph (**C**) and FlowSOM (**D**). In order to test the effect of the resolution parameter, the functions were ran concurrently with a different setting for the cluster resolution (left panel). For FlowSOM, the number of metaclusters was adjusted. The time needed for classification of one sample (mid panel) and the memory consumption during the classification (right panel) was measured.

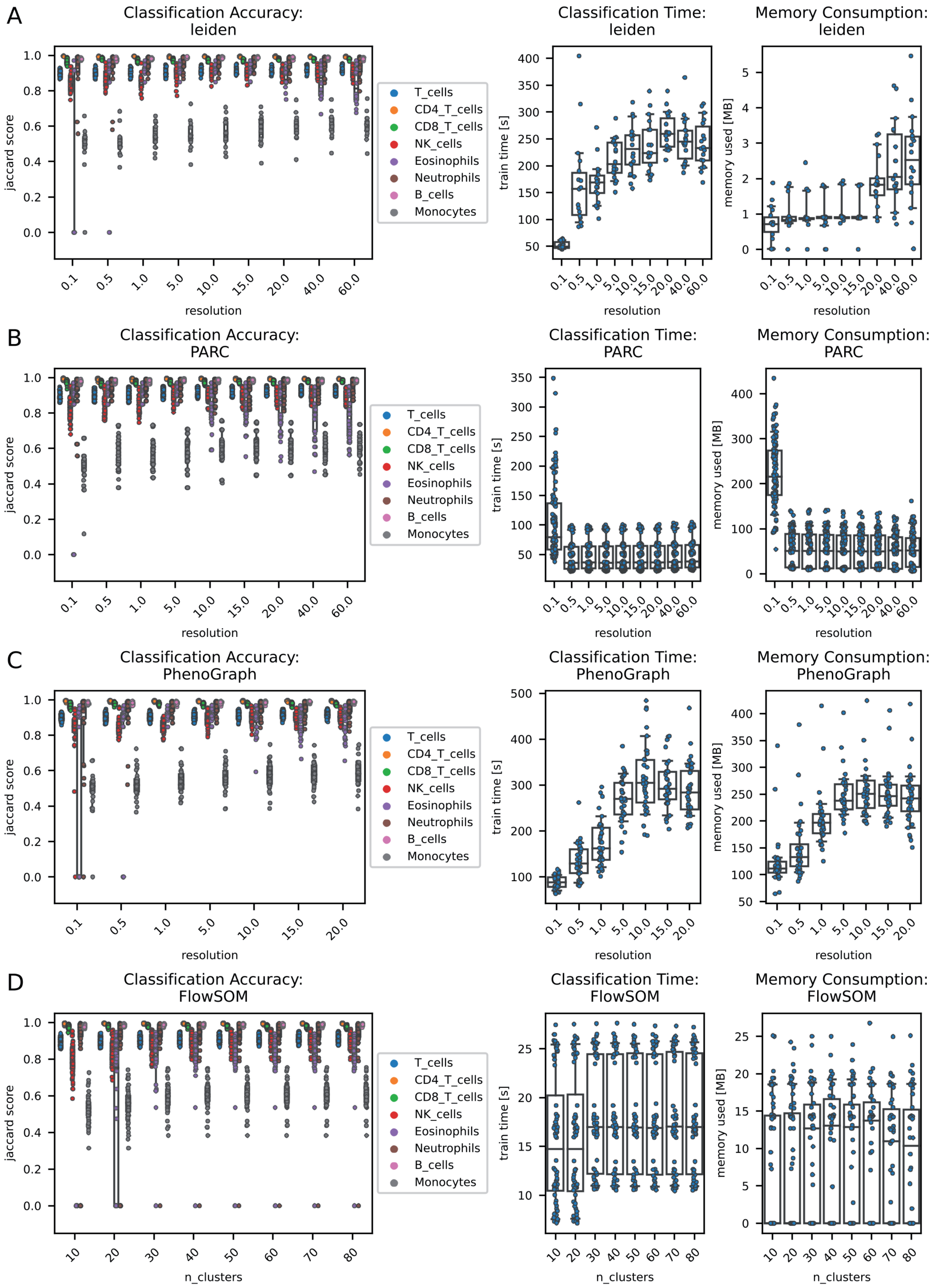


**Supplementary Figure S23: Classifier characterization semi-supervised gating.** The dataset consisted of 18 samples of mouse spleen cells (flow cytometry; Dataset 3). Cell line identification was performed as described in the methods section using the clustering algorithms were leiden (**A**), PARC (**B**), phenograph (**C**) and FlowSOM (**D**). In order to test the effect of the resolution parameter, the functions were ran concurrently with a different setting for the cluster resolution (left panel). For FlowSOM, the number of metaclusters was adjusted. The time needed for classification of one sample (mid panel) and the memory consumption during the classification (right panel) was measured.

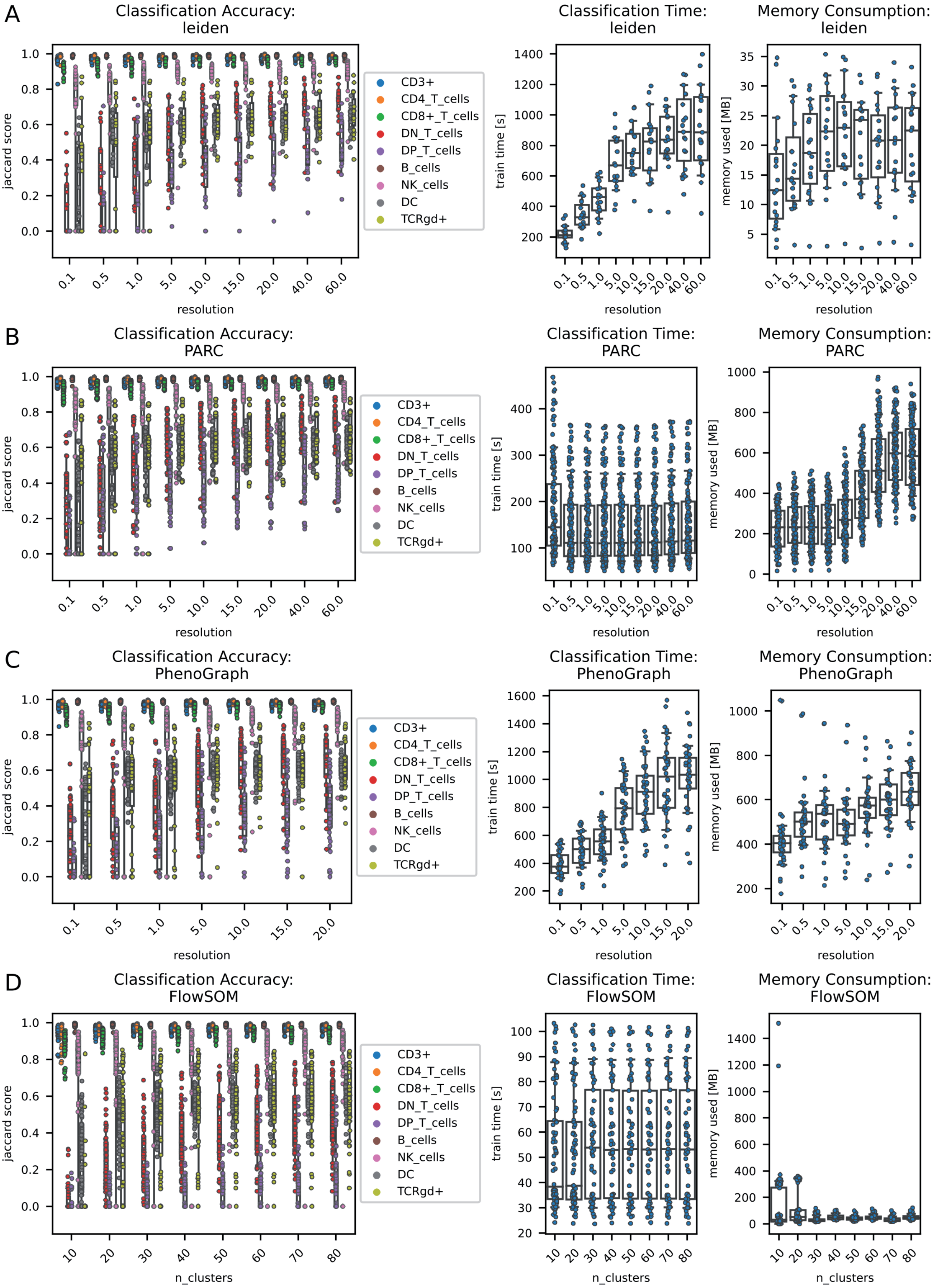


**Supplementary Figure S27: Classifier characterization semi-supervised gating.** The dataset consisted of 19 samples of human peripheral blood cells (mass cytometry; Dataset 7). Cell line identification was performed as described in the methods section using the clustering algorithms were leiden (**A**), PARC (**B**), phenograph (**C**) and FlowSOM (**D**). In order to test the effect of the resolution parameter, the functions were ran concurrently with a different setting for the cluster resolution (left panel). For FlowSOM, the number of metaclusters was adjusted. The time needed for classification of one sample (mid panel) and the memory consumption during the classification (right panel) was measured.

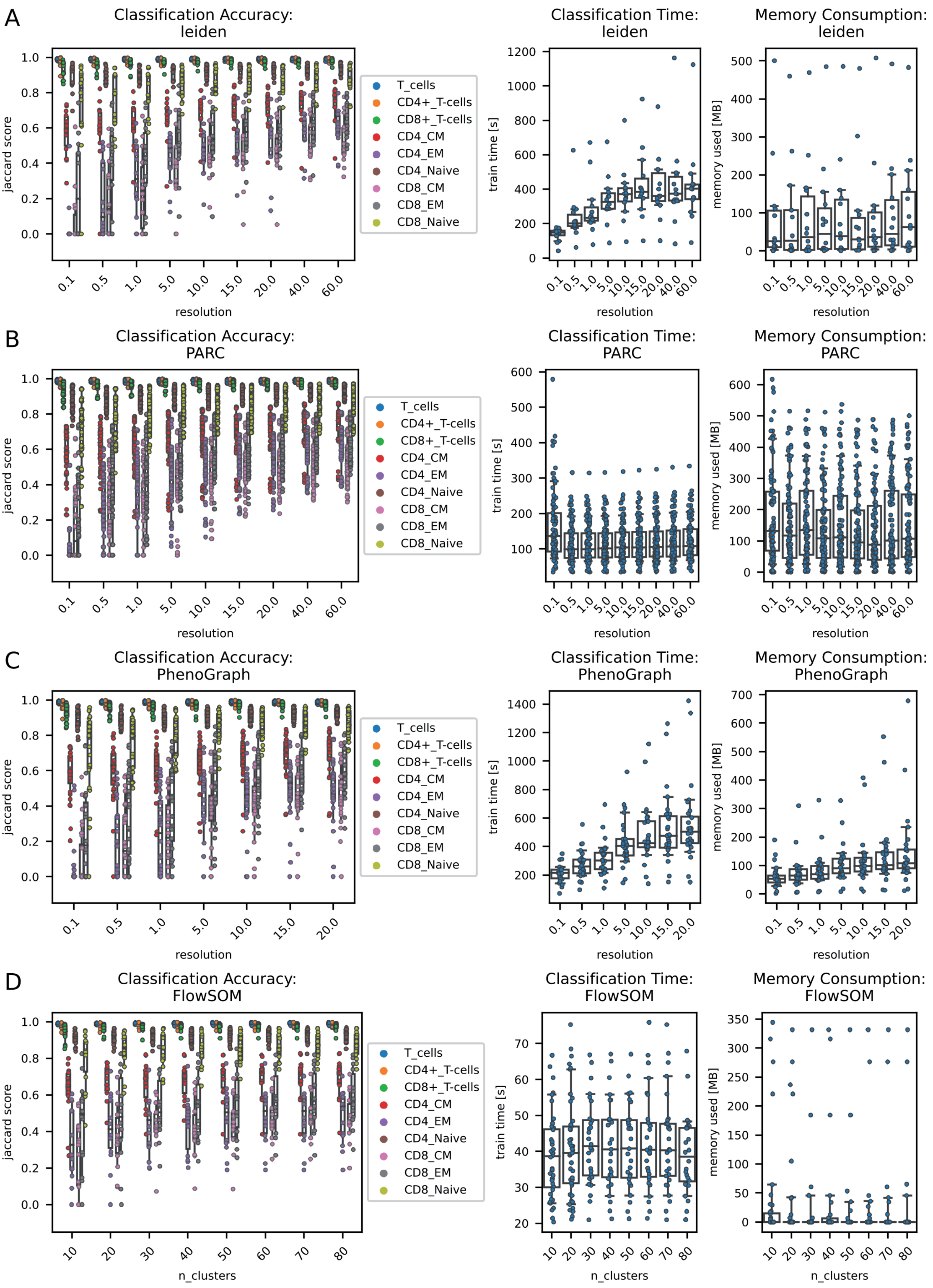


**Supplementary Figure S24: Classifier characterization semi-supervised gating.** The dataset consisted of 14 samples of human peripheral blood cells (flow cytometry; Dataset 4). Cell line identification was performed as described in the methods section using the clustering algorithms were leiden (**A**), PARC (**B**), phenograph (**C**) and FlowSOM (**D**). In order to test the effect of the resolution parameter, the functions were ran concurrently with a different setting for the cluster resolution (left panel). For FlowSOM, the number of metaclusters was adjusted. The time needed for classification of one sample (mid panel) and the memory consumption during the classification (right panel) was measured.

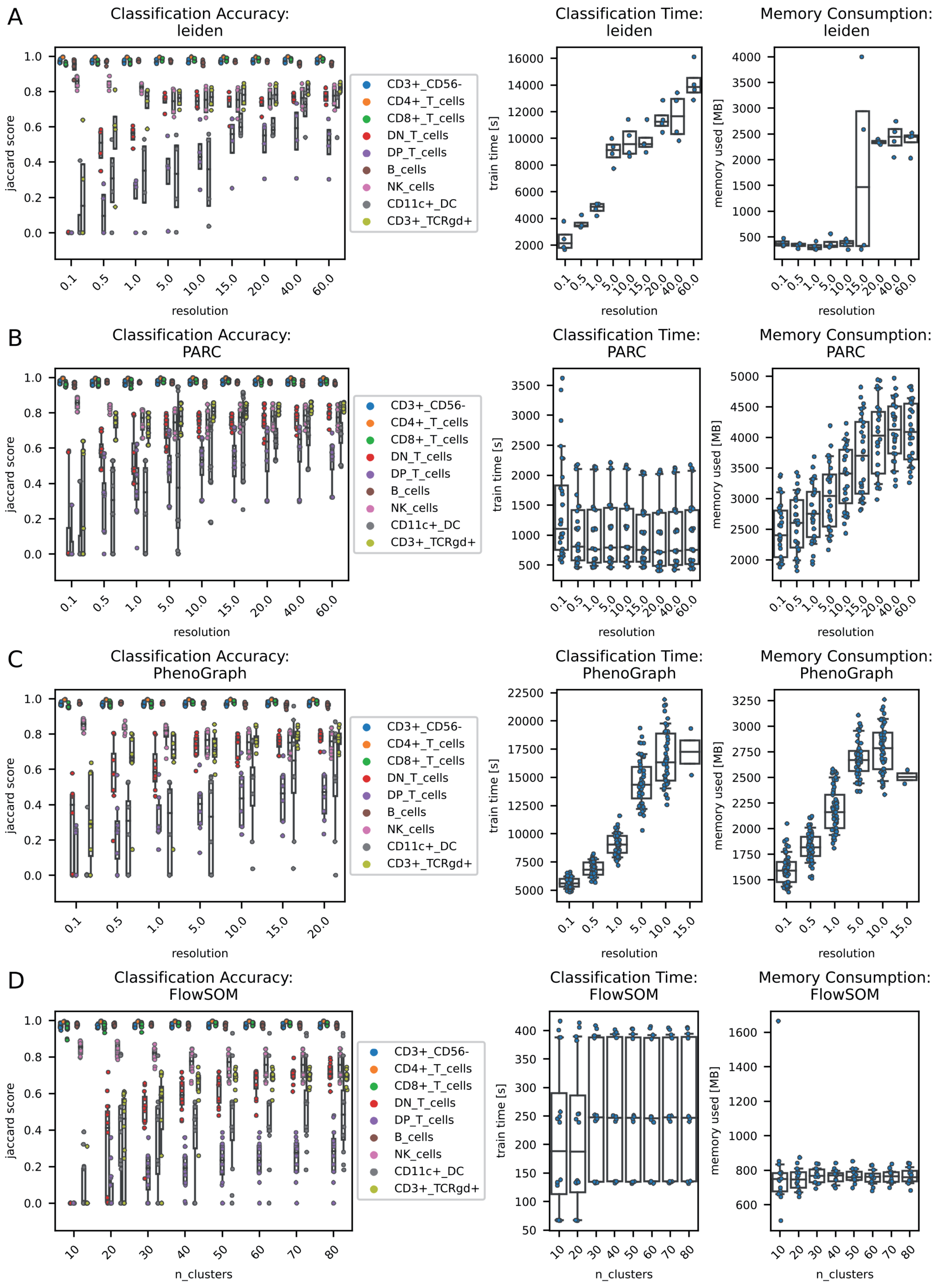


**Supplementary Figure S25: Classifier characterization semi-supervised gating.** The dataset consisted of 4 samples of human peripheral blood cells (flow cytometry; Dataset 6). Cell line identification was performed as described in the methods section using the clustering algorithms were leiden (**A**), PARC (**B**), phenograph (**C**) and FlowSOM (**D**). In order to test the effect of the resolution parameter, the functions were ran concurrently with a different setting for the cluster resolution (left panel). For FlowSOM, the number of metaclusters was adjusted. The time needed for classification of one sample (mid panel) and the memory consumption during the classification (right panel) was measured.

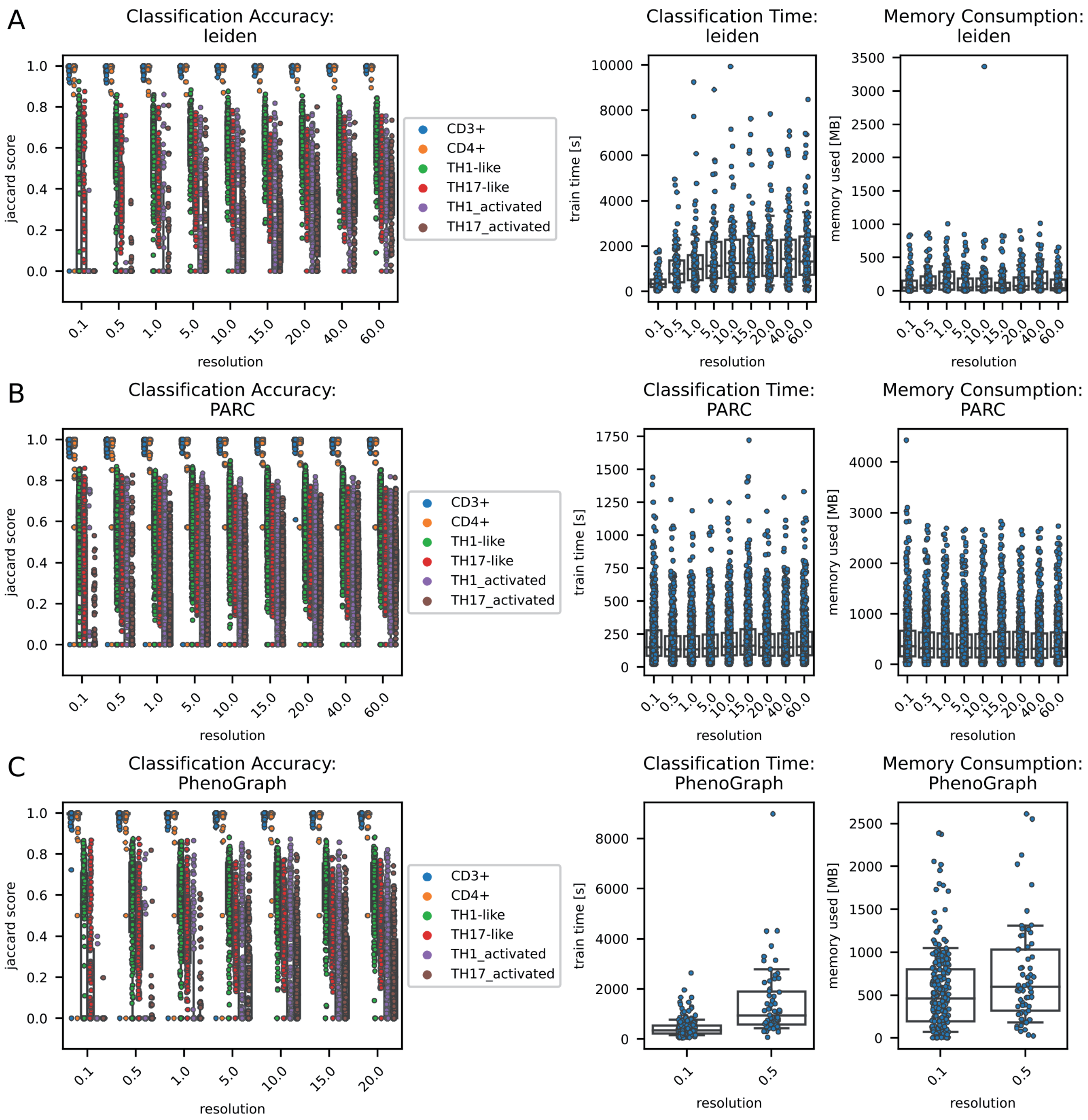


**Supplementary Figure S26: Classifier characterization semi-supervised gating.** The dataset consisted of 107 samples of human peripheral blood cells (flow cytometry; Dataset 5). Cell line identification was performed as described in the methods section using the clustering algorithms were leiden (**A**), PARC (**B**) and phenograph (**C**) . In order to test the effect of the resolution parameter, the functions were ran concurrently with a different setting for the cluster resolution (left panel). The time needed for classification of one sample (mid panel) and the memory consumption during the classification (right panel) was measured.

### Supplementary Tables

**Supplementary Table S1: Panel of datasets 1-3**

Panel for lineage analysis of mouse bone marrow, peripheral blood and spleen.

| Marker | Channel | Clone | Vendor | Catalog # | Dilution |
|---|---|---|---|---|---|
| **DAPI** | BUV496 | N/A | Sigma-Aldrich | D9542 | 1:1000 of 1 mg/ml |
| **B220** | APC | RA3-6B2 | BioLegend | 103211 | 1:100 |
| **CD4** | APC-Cy7 | RM4-5 | BioLegend | 100525 | 1:200 |
| **CD8** | BV510 | 53-6.7 | BioLegend | 100751 | 1:200 |
| **Siglec-F** | BV421 | S17007L | BioLegend | 155509 | 1:200 |
| **Ly6C** | BV605 | HK1.4 | BioLegend | 128035 | 1:400 |
| **Ly6G** | BUV395 | 1A8 | BD | 565964 | 1:200 |
| **NK1.1** | BV711 | PK136 | BioLegend | 108745 | 1:50 |
| **CD11b** | BV786 | M1/70 | BioLegend | 101243 | 1:200 |
| **CD3** | BUV737 | 145-2C11 | BD | 612771 | 1:50 |
| **F4/80** | PE-Cy7 | BM8 | BioLegend | 123113 | 1:400 |
| **CD45** | PerCP-Cy5.5 | 30-F11 | BioLegend | 103131 | 1:1000 |

**Supplementary Table S2: Panel of dataset 4.**

Panel for the diagnostic T-cell analysis dataset.

| Channel | Marker |
|---|---|
| **Alexa Fluor 488-A** | CD183 |
| **PerCP-Cy5-5-A** | CD3 |
| **PE-CF594-A** | CD57 |
| **PE-A** | CD197 |
| **PE-Cy5-A** | CD28 |
| **PE-Cy7-A** | CD8 |
| **APC-A** | CXCR5_(CD185) |
| **Alexa Fluor 700-A** | TCR_g_d_APC_R700 |
| **APC-Cy7-A** | CD45Ro |
| **BV421-A** | ICOS_(CD278) |
| **BV510-A** | HLA_DR |

| | |
|---|---|
| **BV605-A** | CD196 |
| **BV711-A** | PD_1_(CD279) |
| **BUV 735-A** | CD45RA |
| **BV786-A** | CD38 |
| **BUV 395-A** | CCR5_(CD195) |
| **BUV496-A** | CD4 |

**Supplementary Table S3: Panel of dataset 5**
Panel for the diagnostic dataset of the University of Tübingen

| Channel | Marker |
|---|---|
| **PE-Dazzle-A** | CD3 |
| **Alexa Fluor 700-A** | CD4 |
| **BV605-A** | CCR6 |
| **Alexa Fluor 647-A** | CXCR3 |
| **PE-Cy7-A** | CD38 |
| **BV421-A** | HLADR |
| **BV510-A** | Viability |

**Supplementary Table S4: Panel of dataset 6.**
Panel for the OMIP dataset.

| Channel | Marker |
|---|---|
| **BUV395-A** | CD45RA |
| **LIVE DEAD Blue-A** | live_dead |
| **BUV496-A** | CD16 |
| **BUV615-A** | CD314 |
| **BUV563-A** | CCR5 |
| **BUV661-A** | CD39 |
| **BUV737-A** | CD56 |
| **BUV805-A** | CD8 |
| **BV421-A** | CCR7 |
| **Super Bright 436-A** | CD123 |
| **eFluor 450-A** | CD11c |
| **BV480-A** | IgD |

| | |
|---|---|
| **BV510-A** | CD3 |
| **Pacific Orange-A** | CD20 |
| **BV570-A** | IgM |
| **BV605-A** | IgG |
| **BV650-A** | CD28 |
| **BV711-A** | CCR6 |
| **BV750-A** | CXCR5 |
| **BV785-A** | PD_1 |
| **BB515-A** | CD141 |
| **FITC-A** | CD57 |
| **Spark Blue 550-A** | CD14 |
| **PerCP-A** | CD45 |
| **PerCP-Cy5.5-A** | CD2 |
| **PerCP-eFluor 710-A** | TCRgd |
| **PE-Fire810-A** | HLADR |
| **PE-A** | CD159c |
| **CF568-A** | CD4 |
| **PE-Dazzle594-A** | CD337 |
| **PE-Alexa Fluor 610-A** | CD24 |
| **PE-Cy5-A** | CD95 |
| **PE-Alexa Fluor 700-A** | CD25 |
| **PE-Cy7-A** | CXCR3 |
| **APC-A** | CD159a |
| **Alexa Fluor 647-A** | CD1c |
| **Spark NIR685-A** | CD19 |
| **APC-R700-A** | CD127 |
| **APC-H7-A** | CD27 |
| **APC-Fire810-A** | CD38 |
| **BUV395-A** | CD45RA |

**Supplementary Table S5: Panel of dataset 7**
Panel for HIMC CyTOF dataset.

| Marker | Channel |
|---|---|
| **In113Di** | CD57 |
| **In115Di** | live_dead |
| **Sn120Di** | Sn120Di |
| **I127Di** | I127Di |
| **Xe131Di** | Xe131Di |
| **Ba138Di** | Ba138Di |
| **Ce140Di** | beads |
| **Nd142Di** | CD19 |
| **Nd143Di** | CD4 |
| **Nd144Di** | CD8 |
| **Nd146Di** | IgD |
| **Sm147Di** | CD85j |
| **Nd148Di** | CD11c |
| **Sm149Di** | CD16 |
| **Nd150Di** | CD3 |
| **Eu151Di** | CD38 |
| **Sm152Di** | CD27 |
| **Eu153Di** | CD11b |
| **Sm154Di** | CD14 |
| **Gd155Di** | CCR6 |
| **Gd156Di** | CD94 |
| **Gd157Di** | CD86 |
| **Gd158Di** | CXCR5 |
| **Tb159Di** | CXCR3 |
| **Gd160Di** | CCR7 |
| **Dy162Di** | CD45RA |
| **Dy164Di** | CD20 |
| **Ho165Di** | CD127 |
| **Er166Di** | CD33 |
| **Er167Di** | CD28 |
| **Er168Di** | CD24 |
| **Tm169Di** | ICOS |
| **Er170Di** | CD161 |

| | |
|---|---|
| **Yb171Di** | TCRgd |
| **Yb172Di** | PD1 |
| **Yb173Di** | CD123 |
| **Yb174Di** | CD56 |
| **Lu175Di** | HLADR |
| **Yb176Di** | CD25 |
| **BCKG190Di** | BCKG190Di |
| **Ir191Di** | DNA1 |
| **Ir193Di** | DNA2 |
